# AI-designed nuclease performs robust knock out, base editing and prime editing in plants

**DOI:** 10.64898/2026.01.20.700739

**Authors:** Priya Das, Romio Saha, Debasmita Panda, Chandana Ghosh, SP Avinash, Sonali Panda, Mirza J Baig, Kutubuddin A. Molla

## Abstract

Genome-editing applications in biological organisms rely on a limited set of naturally occurring enzymes. Artificial intelligence (AI) is transforming protein design. Here, building on the AI-designed nuclease OpenCRISPR-1, we establish the plant-optimized AI-designed editors (PAiD) as a versatile genome-editing platform in plants. PAiD supports NHEJ-mediated indel generation, adenine and cytosine base editing, and prime editing, with robust performance across multiple loci comparable to SpCas9. These results highlight the translational potential of AI-designed nucleases for plant genome engineering.

## Main text

RNA-guided programmable nucleases enable high-precision genome engineering for applications in agriculture, medicine and biotechnology. Widely used nucleases such as Cas9^1^ and Cas12a^2^, as well as emerging effectors including TnpB^3,4^ and IscB^5^, are derived from bacterial systems and often exhibit constraints in non-native eukaryotic contexts. Artificial intelligence-driven protein designing is now transforming nuclease engineering by leveraging large language models trained on the vast natural diversity of protein sequences^6^. Recently, OpenCRISPR-1 (OC1), an AI-generated RNA-guided nuclease comprising 1,380 amino acids and differing from the prototypical SpCas9 by 403 amino acid substitutions, was shown to support efficient genome editing in human cells^7^. However, its compatibility with diverse tracrRNA architectures, amenability to the development of advanced editing platforms such as cytosine and adenine base editors and prime editors, and functionality in plant systems have not yet been established. Here, we develop and validate a suite of OC1-based genome-editing tools for plants that support highly efficient gene knockout, base editing and prime editing, demonstrating the versatility of AI-designed nucleases in plant systems. For simplicity, we hereafter refer to this platform as plant-optimised AI-designed editors (PAiD).

We implemented a standardized workflow in which PAiD-based plasmid constructs and their SpCas9-based counterparts were transfected into rice protoplasts^8^, followed by genomic DNA isolation 72 h post-transfection and deep amplicon sequencing to quantify editing outcomes (Fig. 1-a1). Protoplast transfection efficiencies consistently exceeded 80%, as assessed by reporter expression (Fig. 1-a2). To establish a platform for targeted double-strand break (DSB) induction in plant genomes, we codon-optimized the OC1 sequence and cloned it into the pRGEB32 backbone^9^, replacing the SpCas9 coding region to generate the PAiD nuclease expression vector (Fig. 1-b1-b2, supplementary fig. 1). For an initial assessment, two guide RNAs spaced 573 bp apart were cloned into a polycistronic tRNA–gRNA (PTG) cassette^9^ and expressed under the OsU3 promoter in the PAiD vector to target the *OsCYP75B4* gene (Supplementary figure 2, 3). Protoplast transfection followed by PCR and gel analysis revealed a ~570-bp deletion (Fig. 1-b3), indicating that PAiD induced DSBs at both target sites, resulting in excision of the intervening region. The smaller PCR amplicon was cloned, and the targeted deletion was confirmed by Sanger sequencing (Extended data 1). We next evaluated PAiD-mediated editing at the *OsSWEET11, OsSWEET14* and *OsZ3* loci in rice protoplasts. Deep amplicon sequencing revealed robust indel formation at all three targets, with editing efficiencies ranging from 10.0% to 16.8% (Fig. 1-b4-b6, extended data 2-7). PAiD and SpCas9 exhibited comparable indel frequencies at all tested loci.

**Fig. 1.**
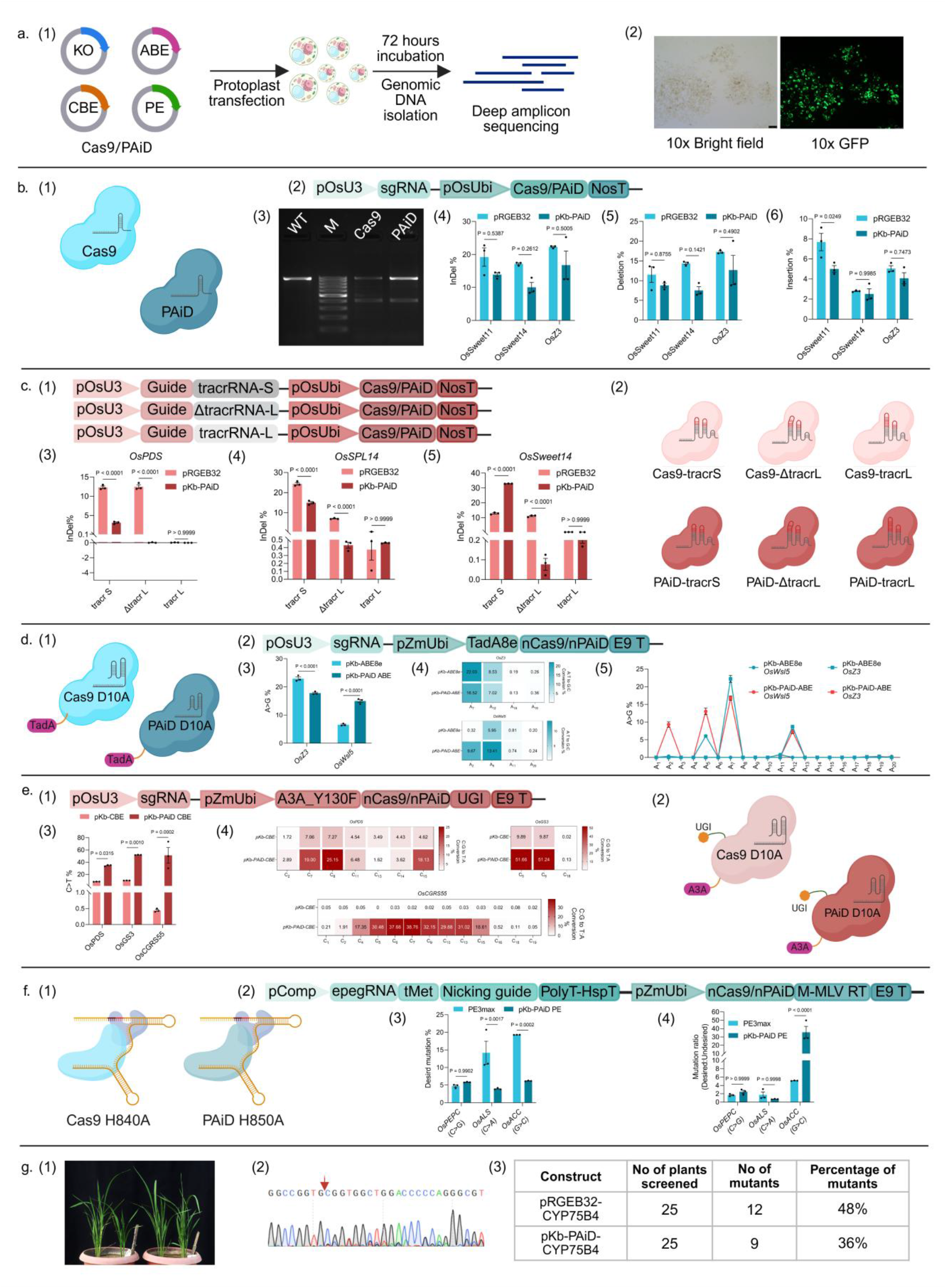
Plant optimized AI-designed editor (PAiD) mediates versatile genome editing in rice. **a1**, Schematic of the workflow for rapid validation of genome editing reagents in-vivo. **a2**, GFP visualization from the transfected protoplast after 16 hours of transfection. *Ubi::GFP* cassette was used as a control to track transfection efficiency. **b1** and **b2**, Cartoons and schematic of the pRGEB32 & pKb-PAiD editing cassettes. **b3**, Gel profile showing a percentage of protoplasts population harbours deletion at the target *OsCYP75B4* locus. WT, PCR amplicon from negative control rice protoplasts (transfected with *Ubi::GFP*). PAiD, lanes with PCR amplicon from PAiD+CYP57B4 PTG transfected protoplasts. Cas9+CYP57B4 PTG was used as positive control. **b4, b5, and b6**, Comparative indel, deletion and insertion efficiencies of Cas9 and PAiD. **c1**, Schematic showing the components of the pRGEB32 and pKb–PAiD vectors containing different tracrRNA variants. **c2**, Cartoon depicting the different vector configurations with tracrRNA variants. **c3, c4, and c5**, Comparative InDel efficiency at three rice loci by both Cas9 and PAiD with canonical and alternative tracrRNA components. **d1 and d2**, Cartoons and schematic of the pKb-ABE & pKb-PAiD-ABE cassettes. **d3**, Comparative A-to-G editing percentage. **d4 and d5**, Editing window at both targets for pKb-ABE & pKb-PAiD-ABE. **e1 and e2**, Schematic and cartoons of the pKb-CBE & pKb-PAiD-CBE cassettes. **e3**, Comparative C-to-T editing percentage. **e4**, Editing window of CBEs at three targets. **f1 and f2**, Cartoons and schematic of the PE3max and pKb-PAiD-PE cassettes. **f3**, Efficiency of Intended edits. **f4**, Ratio of intended to unintended editing efficiencies. **g1**, *cyp75b4* knockout plant lines generated using PAiD. **g2**, Sanger chromatogram of a PAiD-mediated cyp75b4 knockout plant showing the target locus with overlapping peaks indicative of biallelic heterozygous mutations. **g3**, Comparison of editing efficiency in T0 plant lines for *OsCYP75B4* target.

In most SpCas9-mediated genome-editing applications, target recognition is guided by a chimeric single-guide RNA (sgRNA) generated by fusing the crRNA and tracrRNA via an artificial linker^1^. In addition to this canonical short tracrRNA (tracrRNA-S), *Streptococcus pyogenes* encodes a longer tracrRNA variant (tracrRNA-L) that folds into a natural single-guide RNA and directs Cas9 to repress its own promoter, thereby preventing autoimmunity^10^. We recently showed that tracrRNA-L and a truncated derivative (ΔtracrRNA-L) can efficiently support genome editing in eukaryotic systems^11^. Here, we generated PAiD expression vectors incorporating tracrRNA-L and ΔtracrRNA-L modules for guide RNA expression to assess DSB induction and indel formation (Fig. 1-c1-c2, Supplementary fig. 4, 5). We compared PAiD and canonical SpCas9 systems using tracrRNA-S, tracrRNA-L and ΔtracrRNA-L at the *OsPDS, OsSPL14* and *OsSWEET14* loci in rice protoplasts. SpCas9 supported efficient indel formation at all three loci with tracrRNA-S and ΔtracrRNA-L, whereas tracrRNA-L alone consistently showed minimal activity (Fig. 1-c3-c5, extended data 8-25, 42-44). In contrast, PAiD-mediated editing was strongly dependent on the canonical tracrRNA-S, with robust indel formation at *PDS, SPL* and *SWEET14*, while tracrRNA-L and ΔtracrRNA-L resulted in markedly reduced or negligible editing in a locus-dependent manner (Fig. 1-c3-c5, extended data 8-25 and 42-44). The restricted compatibility of PAiD with non-canonical tracrRNA variants likely reflects altered RNA-protein interfaces arising from its extensive AI-driven sequence divergence from SpCas9.

Next, we evaluated the compatibility of PAiD for base editor development. Base editing enables precise installation of targeted single-nucleotide substitutions without introducing genomic DSB^12^. Adenine base editors (ABE) mediate A·T-to-G·C conversions^13^, whereas cytosine base editors (CBE) install C·G-to-T·A substitutions^14^. Modern ABEs are typically generated by fusing a Cas9(D10A) nickase to the laboratory-evolved adenosine deaminase TadA8e^15^. Following this strategy, we engineered D10A nickase variants of both Cas9 and PAiD and fused TadA8e to their N termini via a flexible linker to generate Cas9-ABE and PAiD-ABE, respectively (Fig. 1-d1-d2, Supplementary fig. 6). PAiD-ABE efficiently mediated A·T-to-G·C substitutions at both tested loci, *OsZ3* and *OsWSL5*. Notably, PAiD-ABE exhibited significantly higher editing efficiency at the *OsWSL5* locus (15%) compared with Cas9-ABE (6.6%), whereas editing at the *OsZ3* locus was modestly reduced relative to Cas9-ABE (17.8% versus 23%) (Fig. 1-d3, extended data 26-29). PAiD-ABE exhibits an expanded editing window, as evidenced by an approximately 30-fold increase in editing efficiency at an adenine located at protospacer position 2 of *OsWSL5* relative to Cas9-ABE (Fig. 1-d4-d5, extended data 28-29).

To generate PAiD-CBE and Cas9-CBE, we fused a human APOBEC3A variant (A3A-Y130F) and an uracil glycosylase inhibitor (UGI) to the N and C termini of PAiD(D10A) and Cas9(D10A), respectively (Fig. 1-e1-e2, supplementary fig. 7). The cytidine deaminase A3A-Y130F has previously been shown to confer high C-to-T editing efficiency in plant CBE architectures^16^. Across all three target sites tested, *OsPDS, OsGS3 and OsCGRS55*, PAiD-CBE induced significantly higher C·G-to-T·A substitutions than Cas9-CBE (Fig. 1-e3, extended data 30-35). Across these targets, PAiD-CBE exhibited an activity window spanning protospacer positions 2-15 (Fig. 1-e4, extended data 30-35). Although negligible, PAiD-ABE generated similar or reduced Indels byproduct, while PAiD-CBE yielded similar Indels profile except for *OsGS3* (Extended data 45). Collectively, these results demonstrate that PAiD is compatible with both adenine and cytosine base editor platforms and supports highly efficient base editing.

Prime editing enables precise installation of all types of base substitutions as well as predefined insertions and deletions at target loci without the need for a donor template or the induction of DSB^17^. To construct prime editors (PEs), a Cas9(H840A) nickase is typically fused to a Moloney murine leukemia virus reverse transcriptase (M-MLV-RT)^17^. Following this strategy, we generated a corresponding PAiD(H850A) nickase and fused M-MLV-RT to its C terminus to generate PAiD-PE (Fig.1-f1-f2, supplementary fig. 8). Following the PE3max strategy, we designed engineered pegRNAs (epegRNAs) and nicking guide RNAs to install three distinct edits: a C·G-to-G·C substitution in *OsPEPC*, a C·G-to-A·T substitution in *OsALS*, and a G·C-to-C·G substitution at the *OsACC* locus. Compared with Cas9-PE3max, PAiD-PE exhibited modestly higher editing efficiency at the *OsPEPC* target, although this difference did not reach statistical significance (Fig. 1-f3, extended data 36-37). At the *OsALS* and *OsACC* loci, PAiD-PE achieved 4.0% and 6.18% precise edits, respectively (Fig. 1-f3, extended data 38-41). Interestingly, PAiD-PE exhibited comparable ratios of desired to undesired edits at the *OsPEPC* and *OsALS* loci, whereas a significantly improved ratio was observed at the *OsACC* locus (Fig. 1-f4).

Finally, we evaluated the performance of PAiD for editing an endogenous target gene in regenerated rice plants. The *OsCYP57B4* locus was targeted using a PAiD vector harbouring the PTG cassette constructed in the preliminary experiment (Supplementary fig. 2, 3). Rice plants were regenerated following *Agrobacterium*-mediated transformation of calli^18^. Among the hygromycin-resistant plants obtained (Fig. 1-g1), Sanger sequencing of the target locus in 25 independent lines per construct revealed mutation frequencies of 36% (9 of 25 plants) for PAiD and 48% (12 of 25 plants) for pRGEB32 (Fig. 1-g2, g3). We recovered biallelically edited plants harboring a 573-bp deletion that encompassed the intervening sequence between the two induced DSBs (Fig. 1-g2, extended data 46). These results demonstrate that PAiD supports efficient genome editing in regenerated rice plants.

In summary, we developed PAiD-mediated versatile genome-editing platforms and demonstrated efficient NHEJ-driven indel formation, adenine and cytosine base editing, and prime editing in plants. Our results establish that an AI-designed nuclease is compatible with multiple genome-editing modalities and can achieve comparable—and in some contexts superior—performance relative to the naturally occurring SpCas9 across diverse rice target loci. The expanding repertoire of AI-designed effectors^19^, including nucleases and deaminases, is poised to unlock new opportunities in eukaryotic genome engineering by overcoming intrinsic limitations of naturally evolved systems.

## Acknowledgement

P.D. acknowledges the fellowship from the Department of Biotechnology (DBT), Government of India-JRF program. R.S. and C.G. appreciate the fellowship received from Ignite Life Science Foundation, Bangalore, India. S.P.A. acknowledges the support from the University Grants Commission (UGC), Government of India-JRF program. The pRGEB32 and pG3H-PE3max vectors used in this study were gifts from Yinong Yang and Qijun Chen, respectively.

## Author Contribution

K.M. and M.J.B. conceived and designed the study. K.M. supervised the entire project. K.M., P.D. and R.S. planned the experiments. P.D. and R.S. performed experiments and collected all data. D.P. performed the protoplast transfection and prepared the PAiD-based alternative tracr vectors. C.G. prepared the pKb-PAiD mother vector and assisted in cloning. P.D., R.S., D.P., S.P.A. and S.P. performed data analysis. S.P.A. and S.P. constructed the control vectors for ABE and PE experiments. P.D., R.S., D.P., and S.P. analysed the mutant plant lines. K.M., P.D. and R.S. prepared the manuscript. R.S. and P.D. prepared figures. K.M. and M.J.B. revised and finalized the manuscript.

## Data availability

All data supporting the findings of this study are given in the main text and its supplementary files. Amplicon sequencing data were submitted to gene bank (NCBI Bioproject ID-PRJNA1401667).

