## Supplemental information for "AI-designed nuclease performs robust knock out, base editing and prime editing in plants"

#### Table of contents

Extended data

Supplemental Materials and Methods

Supplementary tables

- ❖ Supplemental Table 1. Different vectors, polynucleotides, polypeptide sequences, and their sequence IDs
- ❖ Supplemental Table 2. Target genes with locus ID and guide sequence
- ❖ Supplemental Table 3. List of primers used for cloning different guides
- ❖ Supplemental Table 4. List of primers used for screening of mutants
- ❖ Supplemental Table 5. List of primers used for Deep amplicon sequencing (1st round)
- ❖ Supplemental Table 6. List of primers used for Deep amplicon sequencing (2nd round)

Supplementary sequences

Supplementary figures

#### Extended data

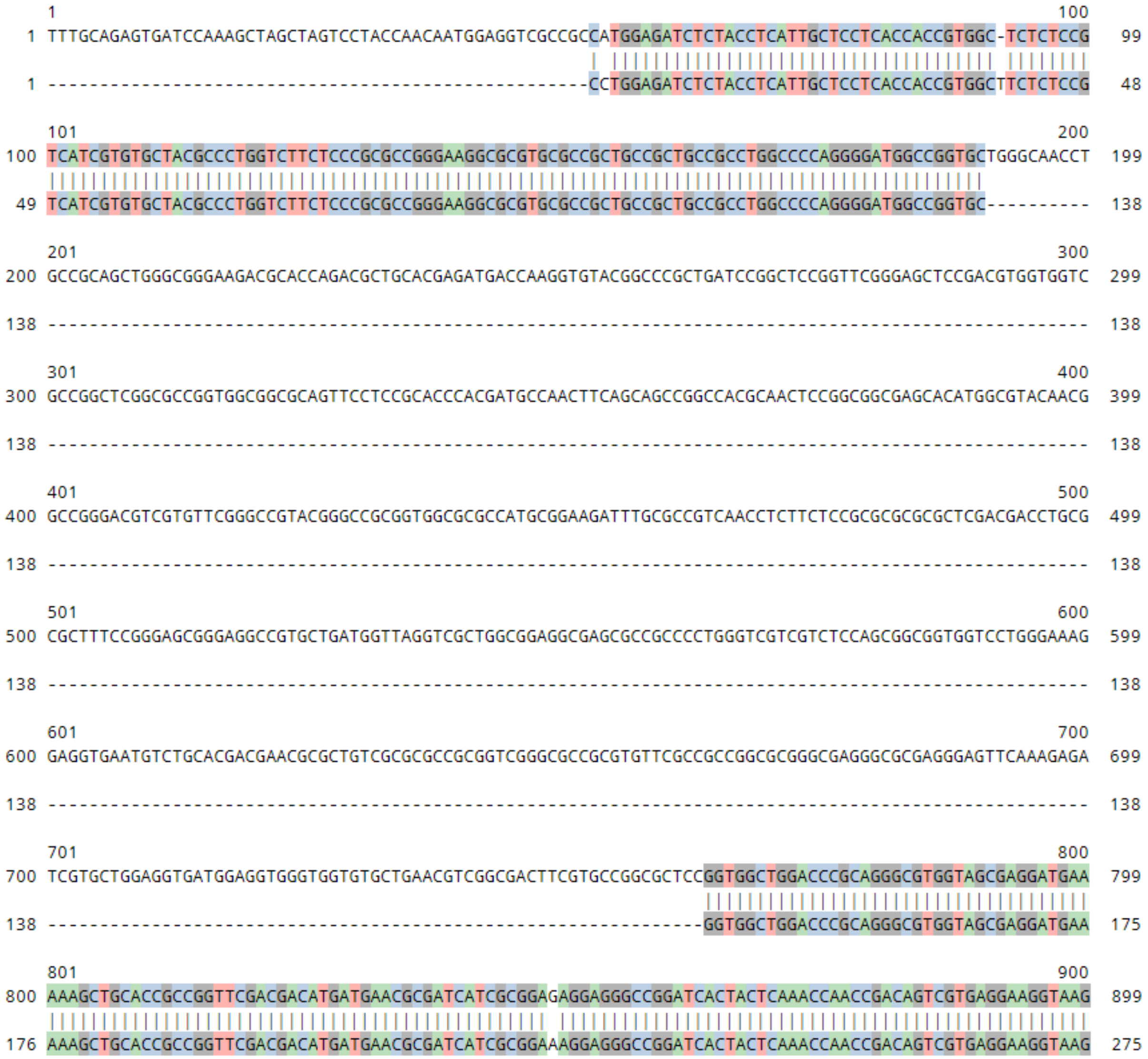


**Extended data 1:** This image depicts the CRISP-Id analysis for *OsCYP75B4*, from the Sanger sequencing data of a single clone from the lower band obtained from protoplast sample. The top strand denotes the control sequence and the lower strand denotes edited allele.

##### *OsSweet11* promoter editing using Cas9

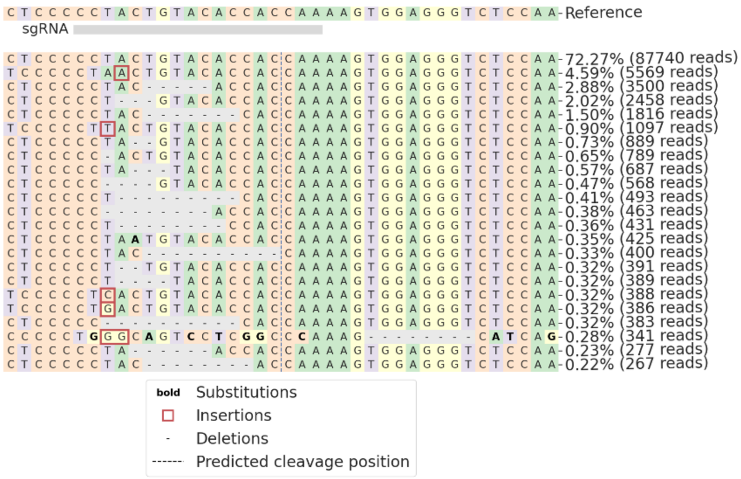


**Extended data 2:** Representative snapshot of CRISPResso2 analysis showing mutations at the target *OsSweet11* promoter locus using the pRGEB32 vector. The "-" marks the deletion in the representative sample compared to the wild-type sequence. The red square box denotes insertion, whereas substitution is marked by **bold letters**.

##### *OsSweet11* promoter editing using PAiD

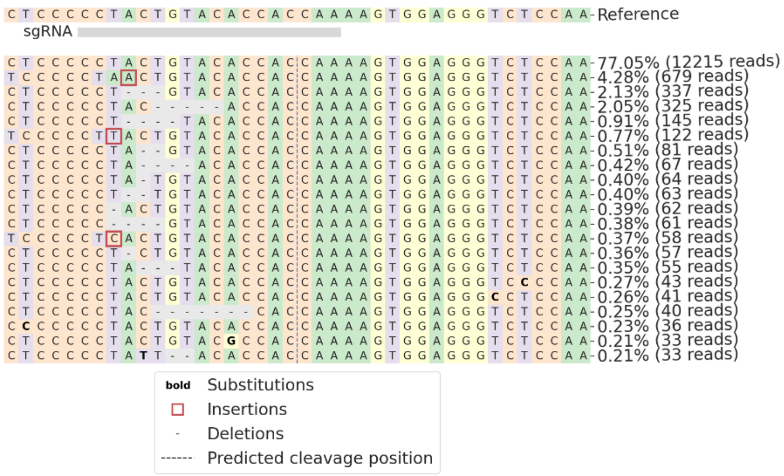


**Extended data 3:** Representative snapshot of CRISPResso2 analysis showing mutations at the target *OsSweet11* promoter locus using the pKb-PAiD vector.

##### *OsSweet14* promoter editing using Cas9

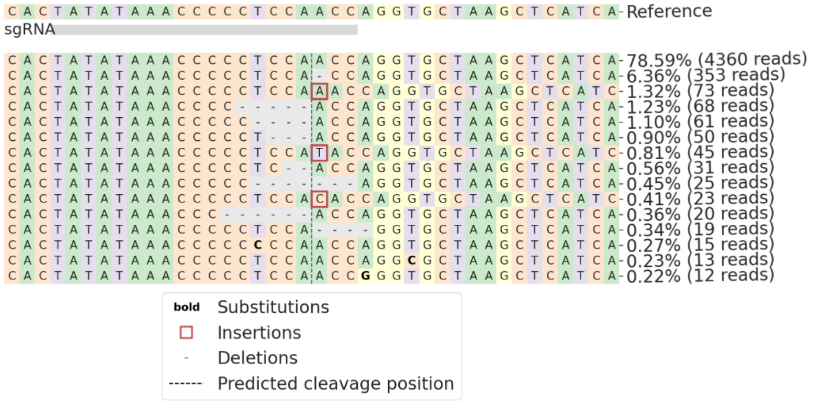


**Extended data 4:** Representative snapshot of CRISPResso2 analysis showing mutations at the target *OsSweet14* promoter locus using the pRGEB32 vector.

##### *OsSweet14* promoter editing using PAiD

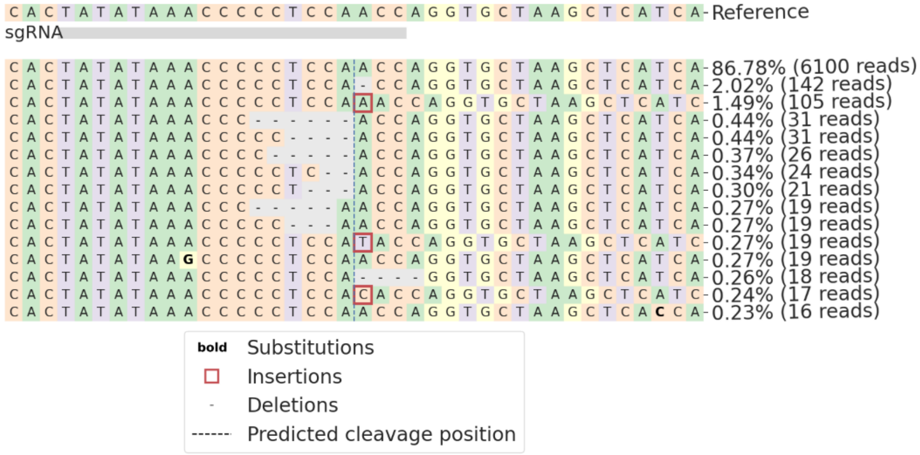


**Extended data 5:** Representative snapshot of CRISPResso2 analysis showing mutations at the target *OsSweet14* promoter locus using the pKb-PAiD vector.

***OsZ3* Knockout by Cas9**

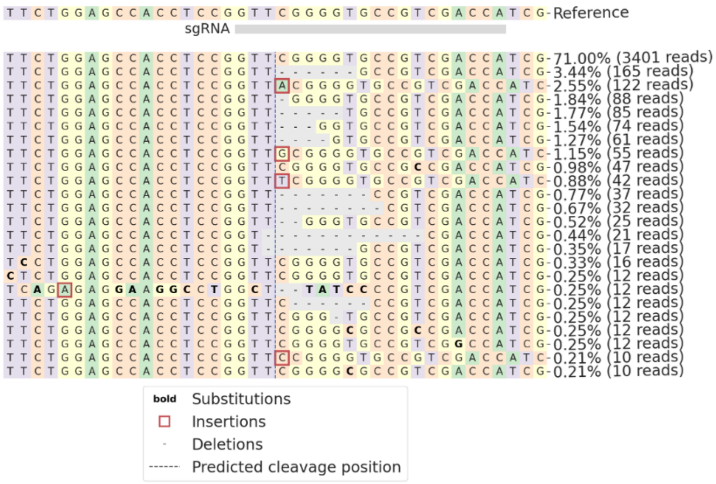

**Extended data 6:** Representative snapshot of CRISPResso2 analysis showing mutations at the target *OsZ3* locus using the pRGEB32 vector.

***OsZ3* Knockout by PAiD**

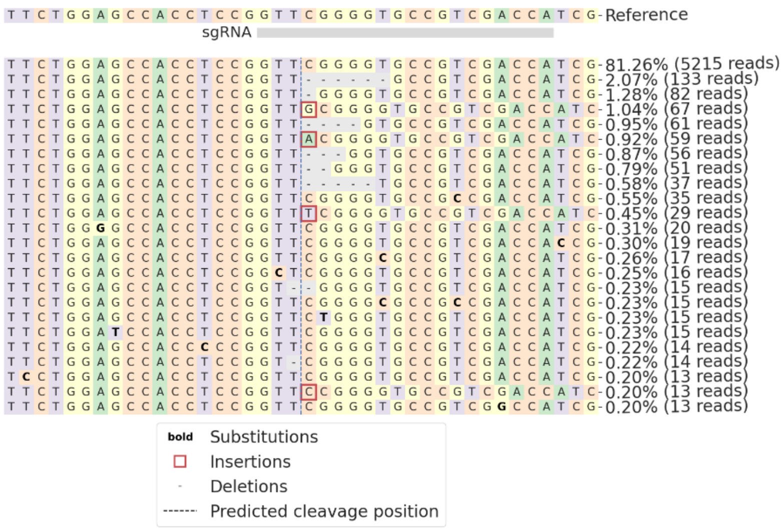

**Extended data 7:** Representative snapshot of CRISPResso2 analysis showing mutations at the target *OsZ3* locus using the pKb-PAiD vector.

##### *OsPDS* Knockout by pKb-Cas9-tracrS

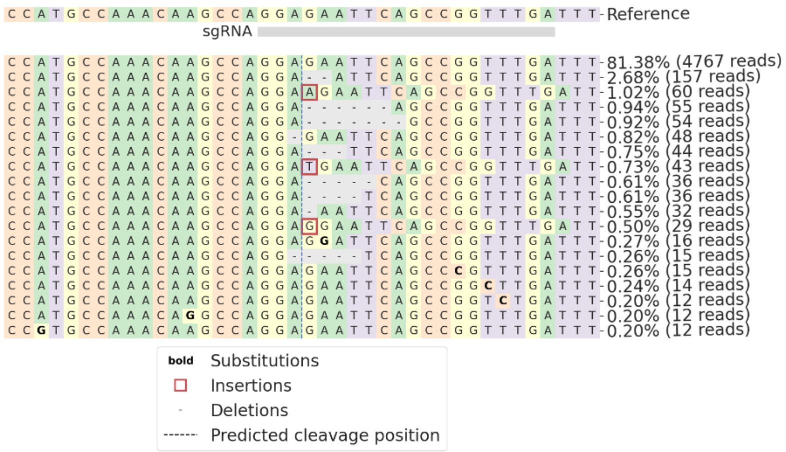


**Extended data 8:** Representative snapshot of CRISPResso2 analysis showing mutations at the target Os*PDS* locus using the pKb-Cas9-tracrS vector. tracrS represents the conventional scaffold for SpCas9.

##### *OsPDS* Knockout by pKb-PAiD-tracrS

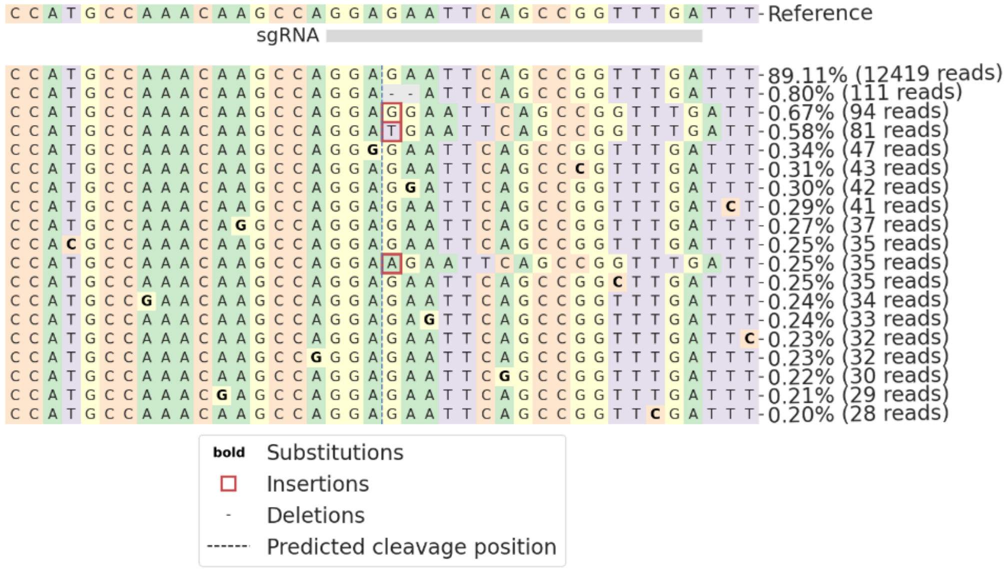


**Extended data 9:** Representative snapshot of CRISPResso2 analysis showing mutations at the target Os*PDS* locus using the pKb-PAiD-tracrS vector. tracrS represents the conventional scaffold for SpCas9.

***OsPDS* Knockout by pKb-Cas9-ΔtracrL**

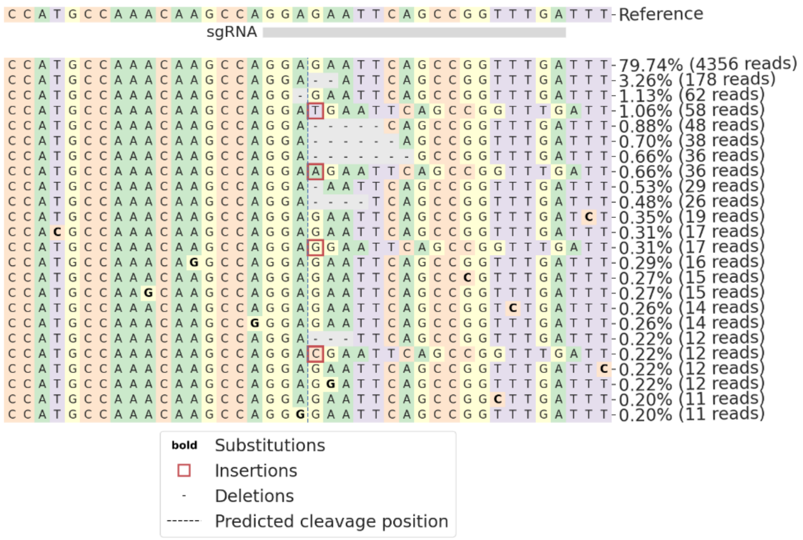


**Extended data 10:** Representative snapshot of CRISPResso2 analysis showing mutations at the target Os*PDS* locus using the pKb-Cas9-ΔtracrL vector. ΔtracrL represents the truncated long tracrRNA

***OsPDS* Knockout by pKb-PAiD-ΔtracrL**

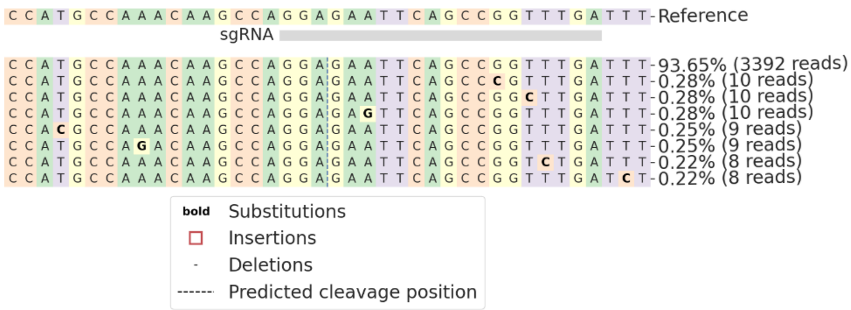


**Extended data 11:** Representative snapshot of CRISPResso2 analysis showing mutations at the target Os*PDS* locus using the pKb-PAiD-ΔtracrL vector. ΔtracrL represents the truncated long tracrRNA.

***OsPDS* Knockout by pKb-Cas9-tracrL**

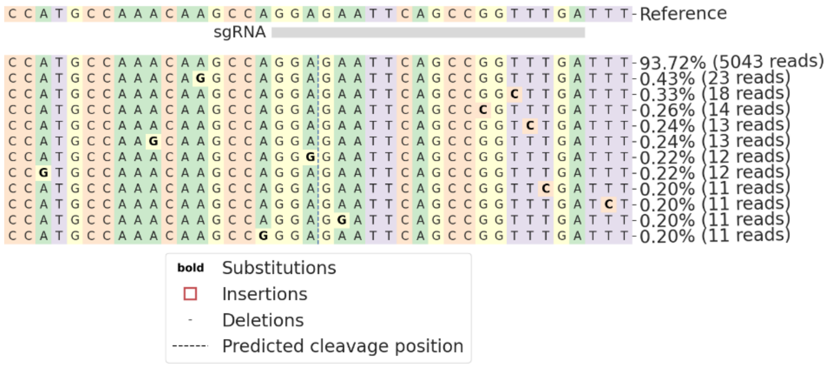

**Extended data 12:** Representative snapshot of CRISPResso2 analysis showing mutations at the target Os*PDS* locus using the pKb-Cas9-tracrL vector. tracrL represents the wild type long tracrRNA.

***OsPDS* Knockout by pKb-PAiD-tracrL**

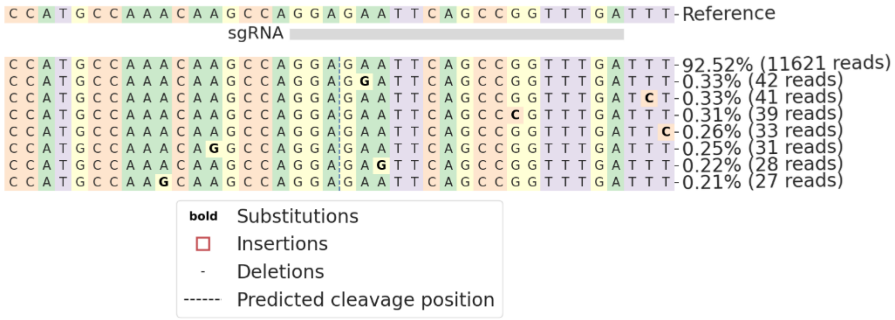

**Extended data 13:** Representative snapshot of CRISPResso2 analysis showing mutations at the target Os*PDS* locus using the pKb-PAiD-tracrL vector. tracrL represents the wild type long tracrRNA.

##### *OsSPL14* Knockout by pKb-Cas9-tracrS

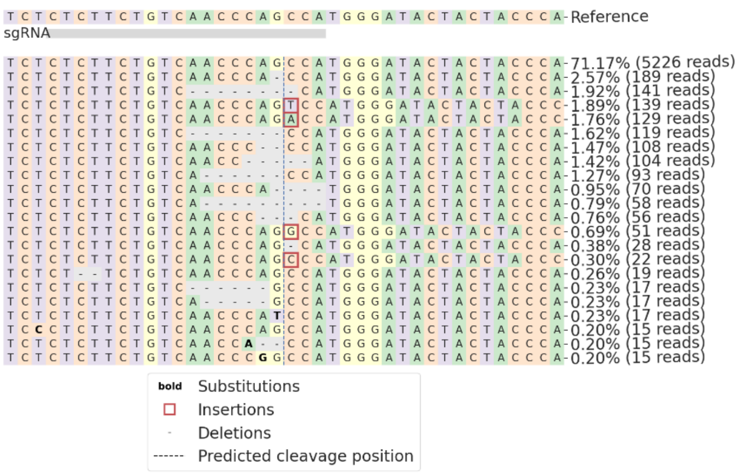

**Extended data 14:** Representative snapshot of CRISPResso2 analysis showing mutations at the target *OsSPL14* locus using the pKb-Cas9-tracrS vector. tracrS represents the conventional scaffold for SpCas9.

##### *OsSPL14* Knockout by pKb-PAiD-tracrS

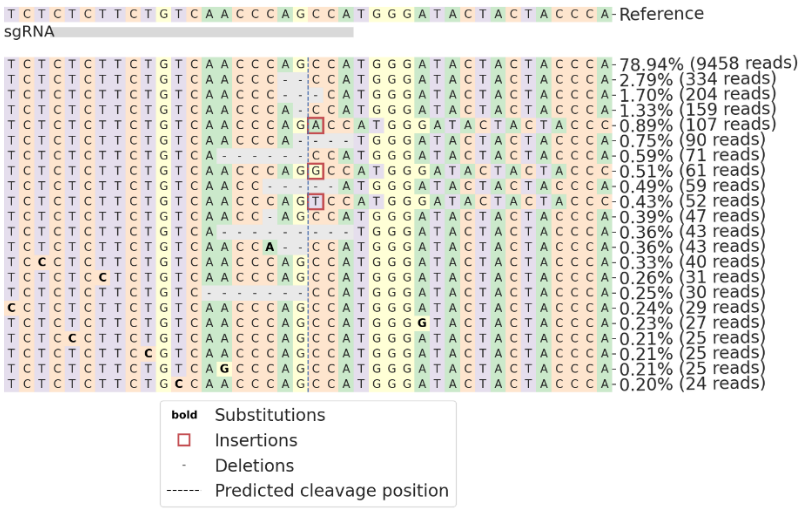

**Extended data 15:** Representative snapshot of CRISPResso2 analysis showing mutations at the target *OsSPL14* locus using the pKb-PAiD-tracrS vector. tracrS represents the conventional scaffold for SpCas9.

##### *OsSPL14* Knockout by pKb-Cas9-ΔtracrL

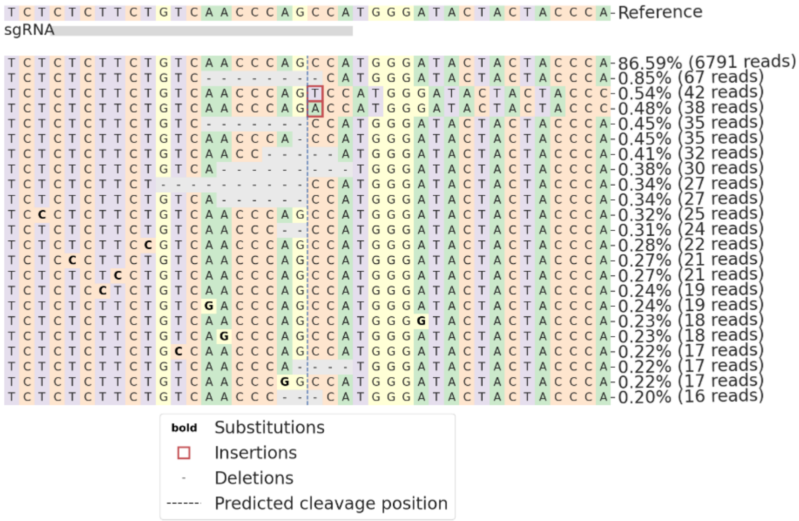

**Extended data 16:** Representative snapshot of CRISPResso2 analysis showing mutations at the target *OsSPL14* locus using the pKb-Cas9-ΔtracrL vector. ΔtracrL represents the truncated long tracrRNA.

##### *OsSPL14* Knockout by pKb-PAiD-ΔtracrL

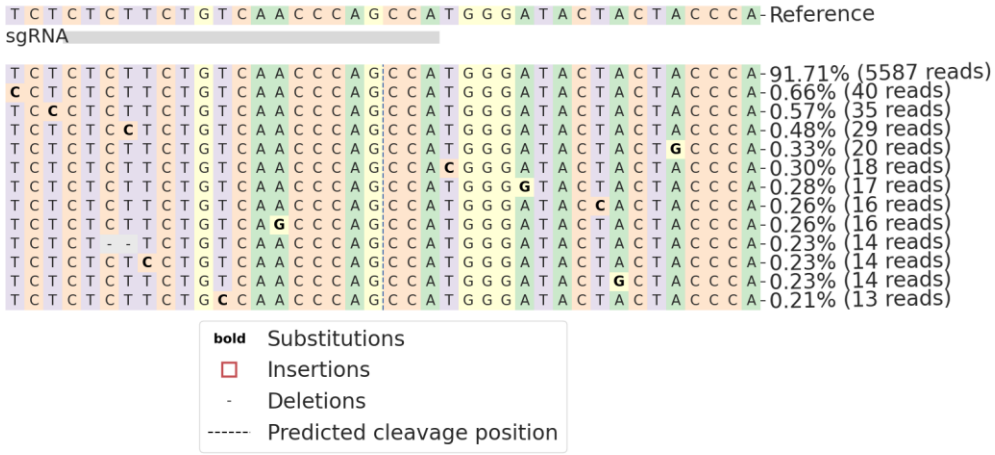

**Extended data 17:** Representative snapshot of CRISPResso2 analysis showing mutations at the target *OsSPL14* locus using the pKb-PAiD-ΔtracrL vector. ΔtracrL represents the truncated long tracrRNA.

***OsSPL14* Knockout by pKb-Cas9-tracrL**

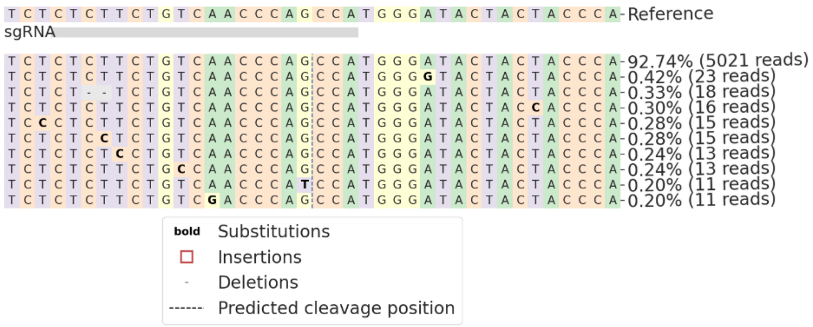


**Extended data 18:** Representative snapshot of CRISPResso2 analysis showing mutations at the target *OsSPL14* locus using the pKb-Cas9-tracrL vector. tracrL represents the wild type long tracrRNA.

***OsSPL14* Knockout by pKb-PAiD-tracrL**

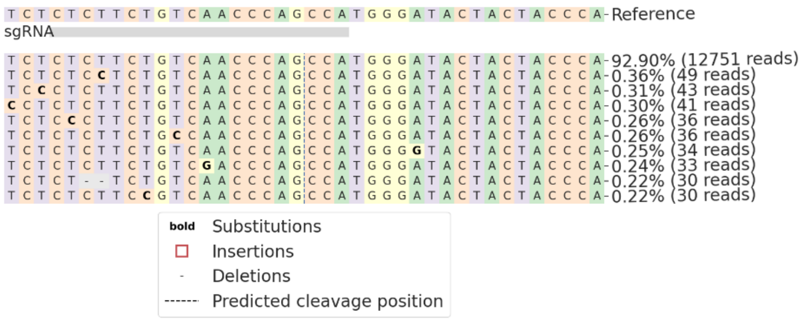


**Extended data 19:** Representative snapshot of CRISPResso2 analysis showing mutations at the target *OsSPL14* locus using the pKb-PAiD-tracrL vector. tracrL represents the wild type long tracrRNA.

***OsSweet14* promoter editing by pKb-Cas9-tracrS**

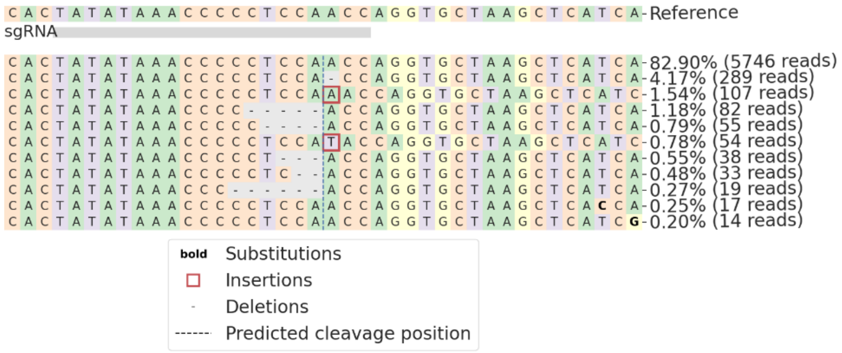

**Extended data 20:** Representative snapshot of CRISPResso2 analysis showing mutations at the target *OsSWEET14* promoter locus using the pKb-Cas9-tracrS vector. tracrS represents the conventional scaffold for SpCas9.

***OsSweet14* promoter editing by pKb-PAiD-tracrS**

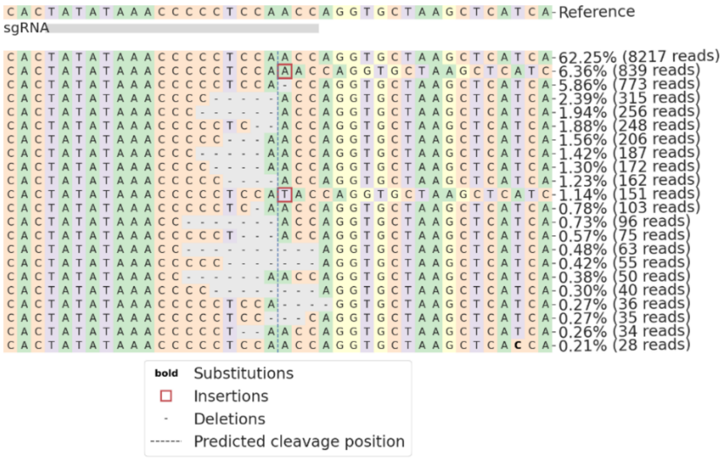

**Extended data 21:** Representative snapshot of CRISPResso2 analysis showing mutations at the target *OsSWEET14* promoter locus using the pKb-PAiD-tracrS vector. tracrS represents the conventional scaffold for SpCas9.

##### *OsSweet14* promoter editing by pKb-Cas9-ΔtracrL

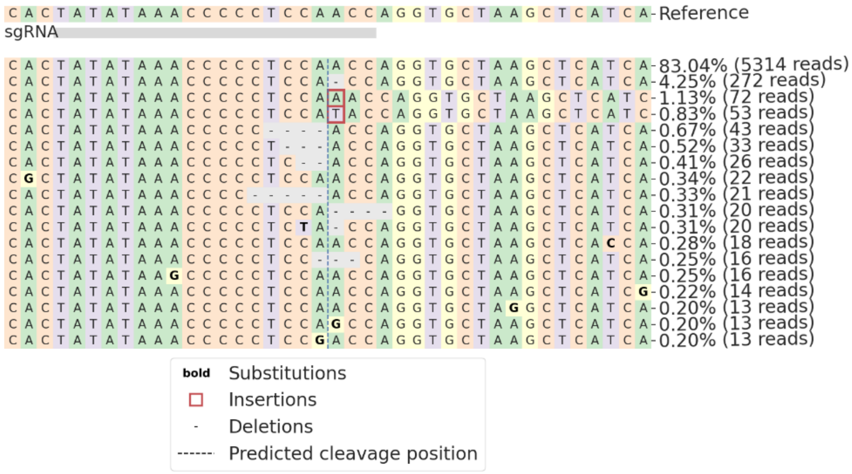


**Extended data 22:** Representative snapshot of CRISPResso2 analysis showing mutations at the target *OsSWEET14* promoter locus using the pKb-Cas9-ΔtracrL vector. ΔtracrL represents the truncated long tracrRNA.

##### *OsSweet14* promoter editing by pKb-PAiD-ΔtracrL

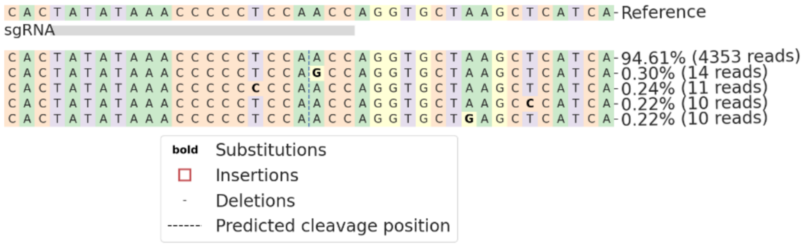


**Extended data 23:** Representative snapshot of CRISPResso2 analysis showing mutations at the target *OsSWEET14* promoter locus using the pKb-PAiD-ΔtracrL vector. ΔtracrL represents the truncated long tracrRNA.

**_OsSweet14_ promoter editing by pKb-Cas9-tracrL**

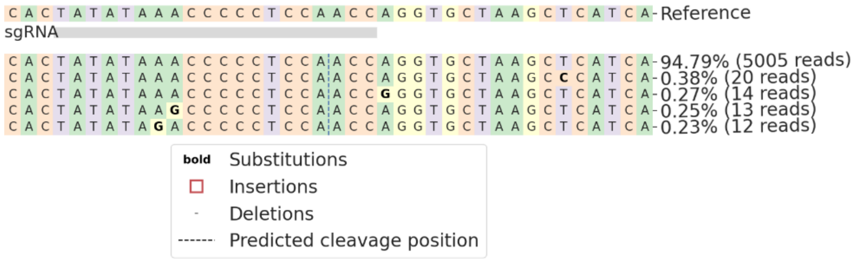


**Extended data 24:** Representative snapshot of CRISPResso2 analysis showing mutations at the target *OsSweet14* promoter locus using the pKb-Cas9-tracrL vector. tracrL represents the wild type long tracrRNA.

**_OsSweet14_ promoter editing pKb-PAiD-tracrL**

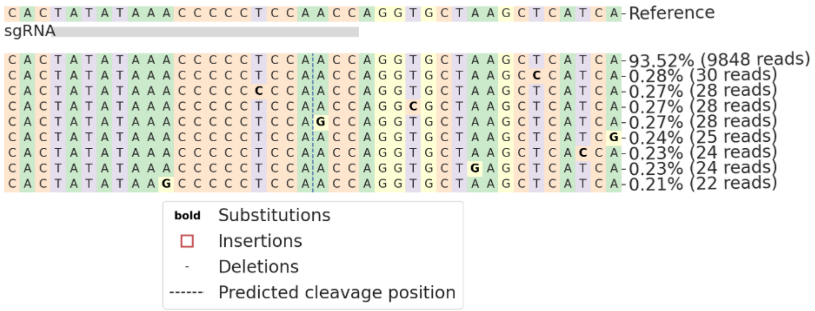


**Extended data 25:** Representative snapshot of CRISPResso2 analysis showing mutations at the target *OsSweet14* promoter locus using the pKb-PAiD-tracrL vector. tracrL represents the wild type long tracrRNA.

***OsZ3*-ABE**

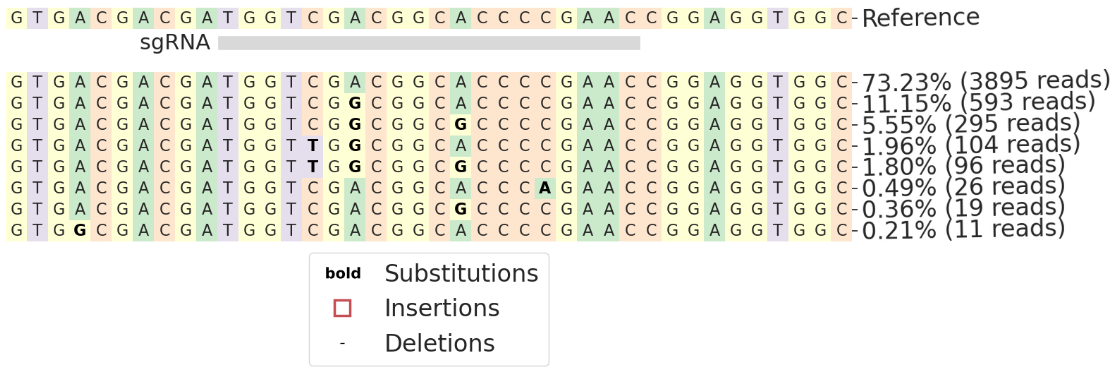

**Extended data 26:** Representative snapshot of CRISPResso2 analysis showing mutations at the target *OsZ3* locus using the pKb-ABE8e vector. The "**bold letters**" indicate substitutions in the representative sample compared to the wild type sequence.

***OsZ3*-PAiD-ABE**

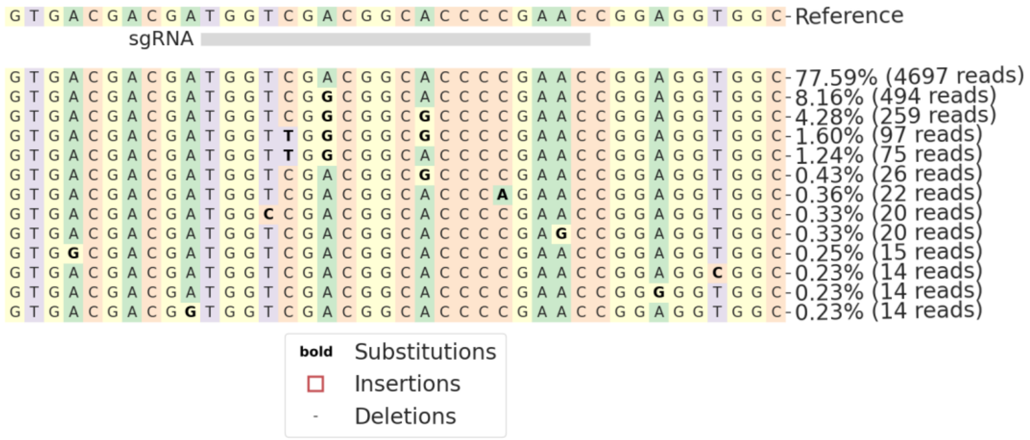

**Extended data 27:** Representative snapshot of CRISPResso2 analysis showing mutations at the target *OsZ3* locus using the pKb-PAiD-ABE vector.

***OsWsl5*-ABE**

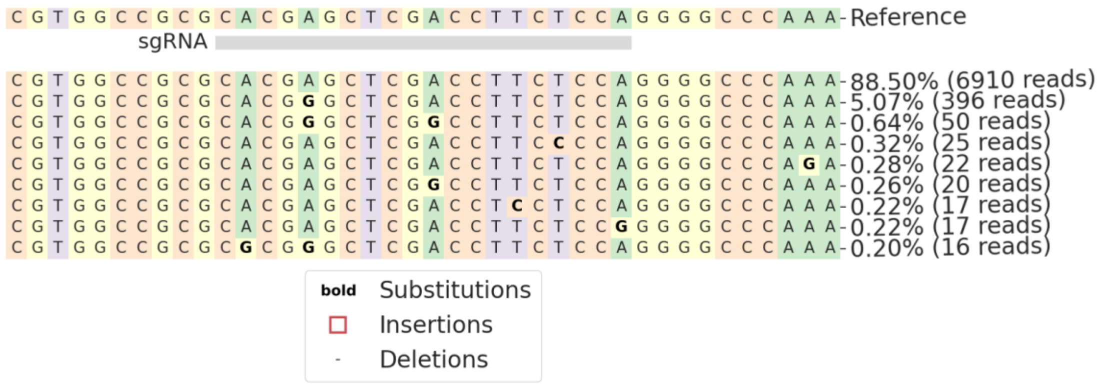

**Extended data 28:** Representative snapshot of CRISPResso2 analysis showing mutations at the target *OsWsl5* locus using the pKb-ABE8e vector.

***OsWsl5*-PAiD-ABE**

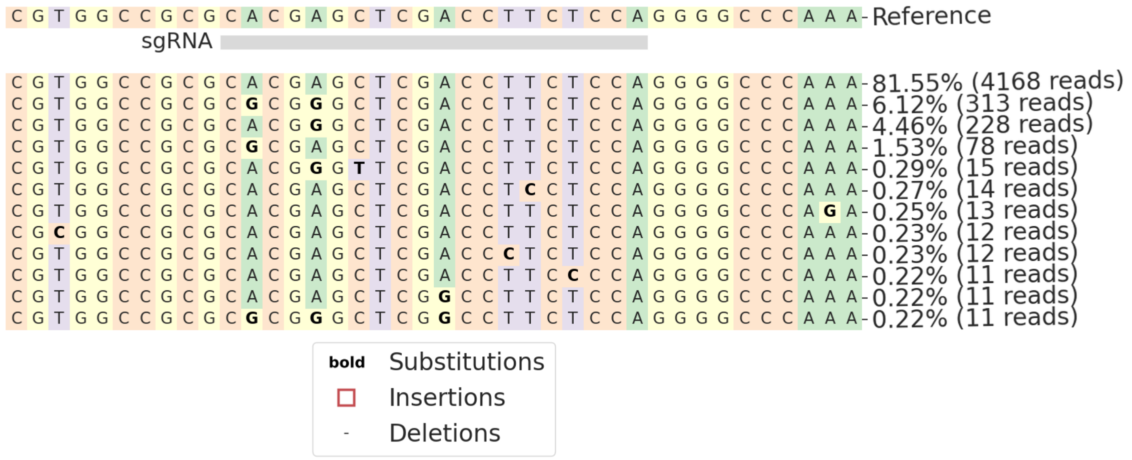

**Extended data 29:** Representative snapshot of CRISPResso2 analysis showing mutations at the target *OsWsl5* locus using the pKb-PAiD-ABE vector.

***OsPDS*-CBE**

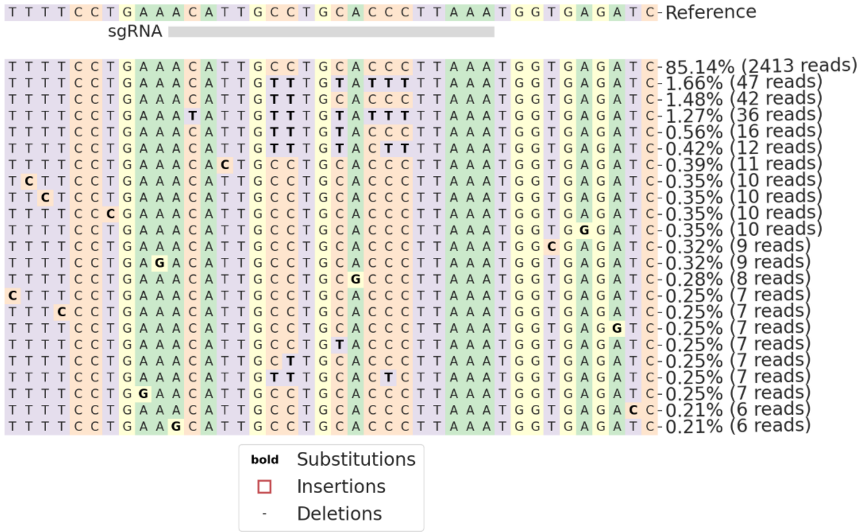


**Extended data 30:** Representative snapshot of CRISPResso2 analysis showing mutations at the target *OsPDS* locus using the pKb-CBE vector.

***OsPDS*-PAiD-CBE**

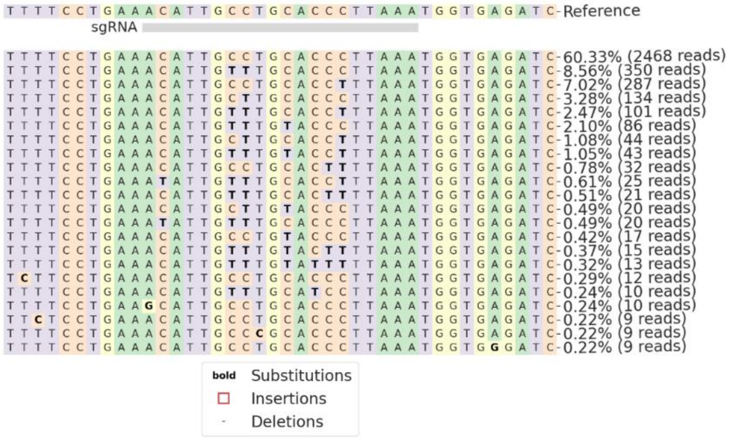


**Extended data 31:** Representative snapshot of CRISPResso2 analysis showing mutations at the target *OsPDS* locus using the pKb-PAiD-CBE vector.

***OsGS3*-CBE**

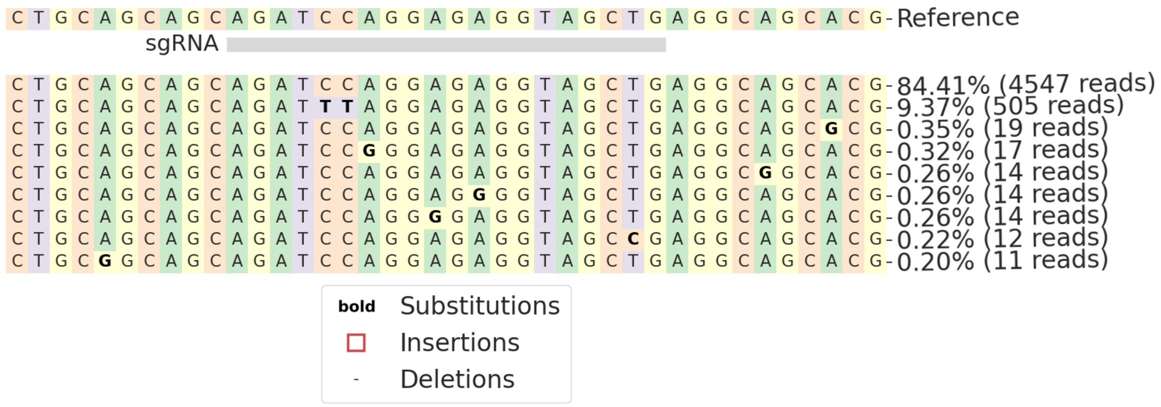


**Extended data 32:** Representative snapshot of CRISPResso2 analysis showing mutations at the target *OsGS3* locus using the pKb-CBE vector.

***OsGS3*-PAiD-CBE**

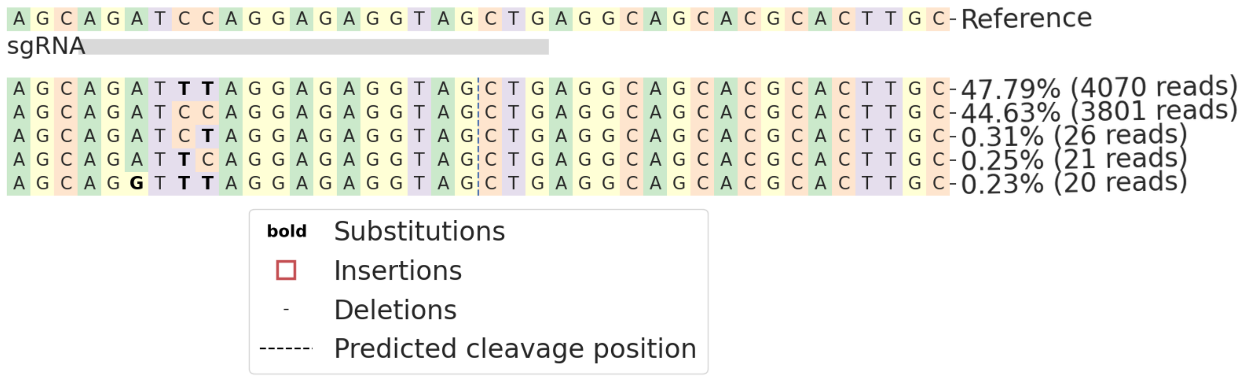


**Extended data 33:** Representative snapshot of CRISPResso2 analysis showing mutations at the target *OsGS3* locus using the pKb-PAiD-CBE vector.

***OsCGRS55*-CBE**

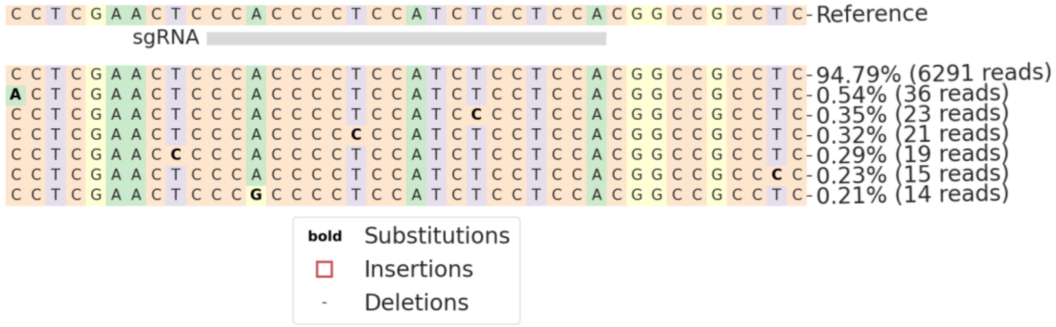


**Extended data 34:** Representative snapshot of CRISPResso2 analysis showing mutations at the target *OsCGRS55* locus using the pKb-CBE vector.

***OsCGRS55*-PAiD-CBE**

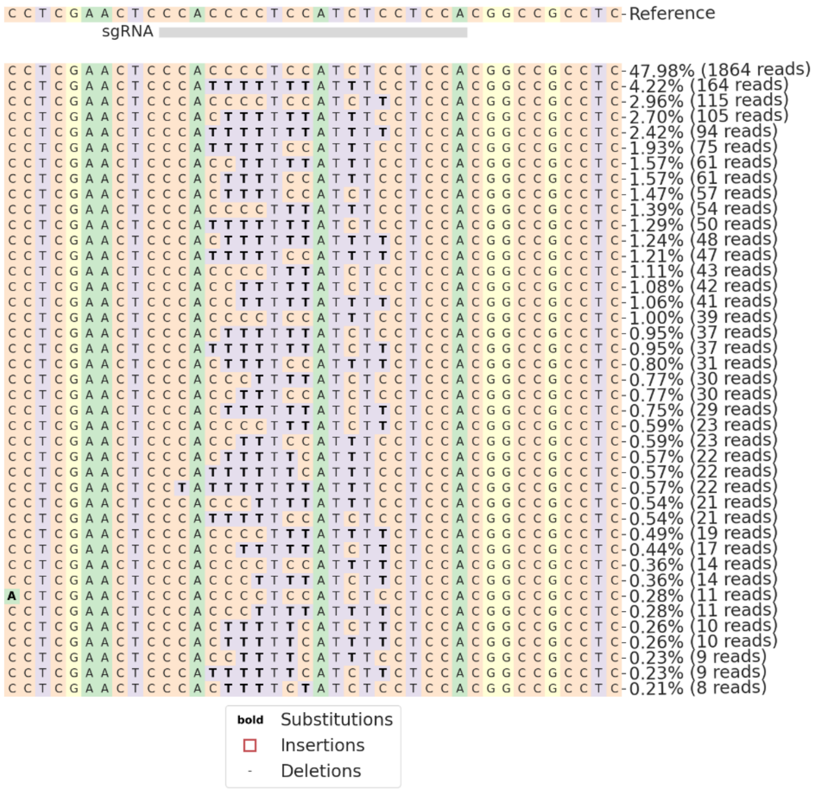


**Extended data 35:** Representative snapshot of CRISPResso2 analysis showing mutations at the target *OsCGRS55* locus using the pKb-PAiD-CBE vector

***OsPEPC*-PE3max (C>G)**

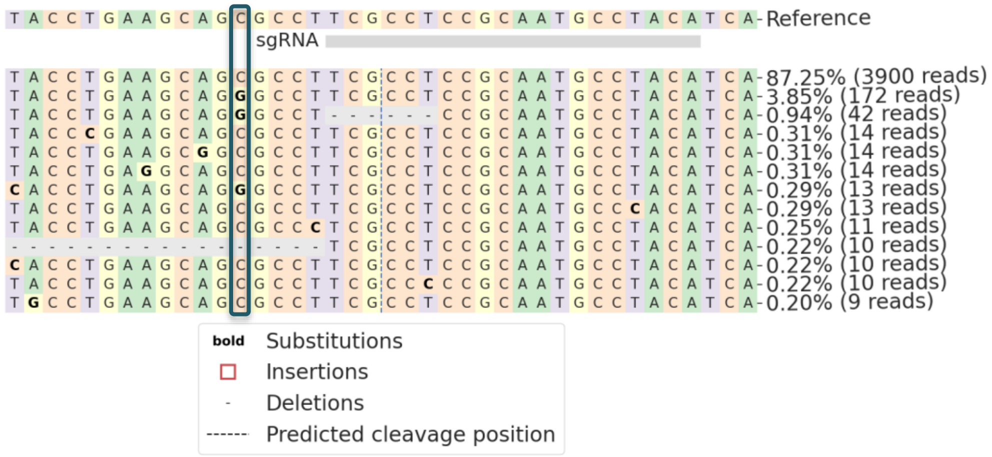


**Extended data 36:** Representative snapshot of CRISPResso2 analysis showing mutations at the target mutations across the reverse transcriptase template sequence for *OsPEPC* locus using the PE3max vector. The "blue vertical box" indicates the desired mutation (C>G) in the representative sample comparative to the wild type sequence.

***OsPEPC*-PAiD PE (C>G)**

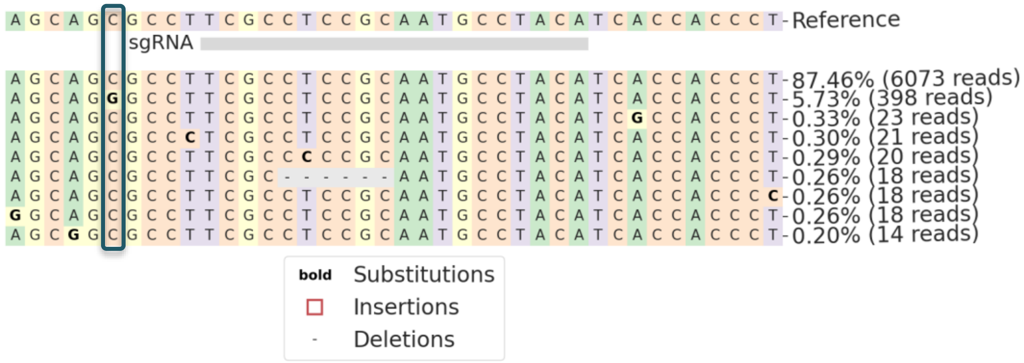


**Extended data 37:** Representative snapshot of CRISPResso2 analysis showing mutations at the target the reverse transcriptase template sequence for *OsPEPC* locus using the pKb-PAiD PE vector.

***OsALS*-PE3max (C>A)**

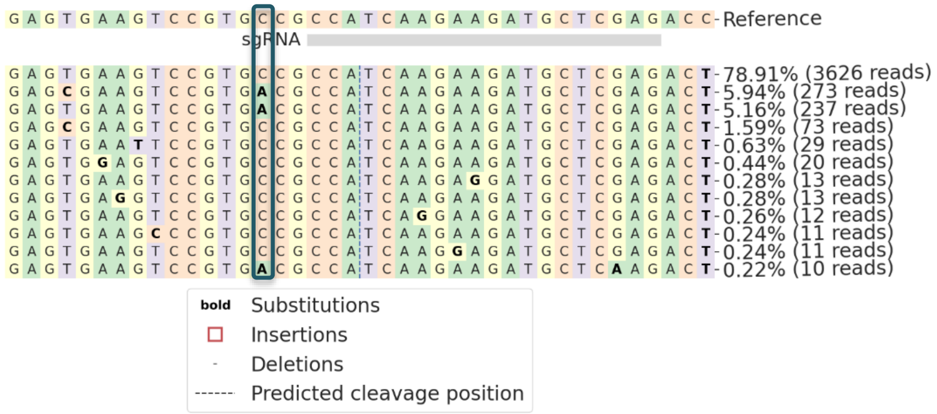


**Extended data 38:** Representative snapshot of CRISPResso2 analysis showing mutations at the target reverse transcriptase template sequence for *OsALS* locus using the PE3max vector.

***OsALS*-PAiD PE (C>A)**

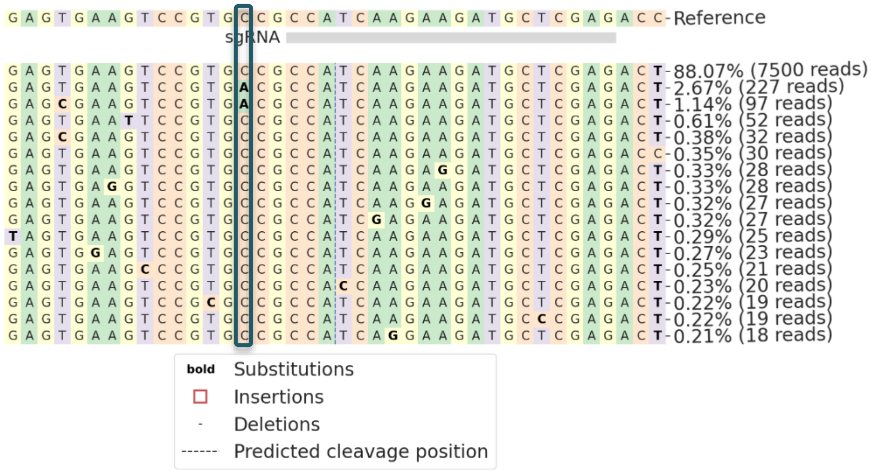


**Extended data 39:** Representative snapshot of CRISPResso2 analysis showing mutations at the target reverse transcriptase template sequence for *OsALS* locus using the pKb-PAiD PE vector.

***OsACC*-PE3max (G>C)**

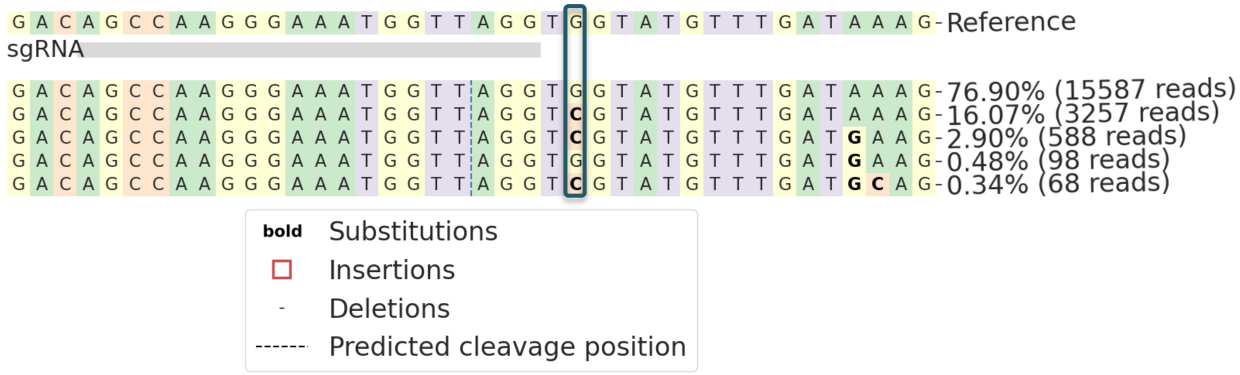


**Extended data 40:** Representative snapshot of CRISPResso2 analysis showing mutations at the target reverse transcriptase template sequence for *OsACC* locus using the PE3max vector.

***OsACC*-PAiD PE (G>C)**

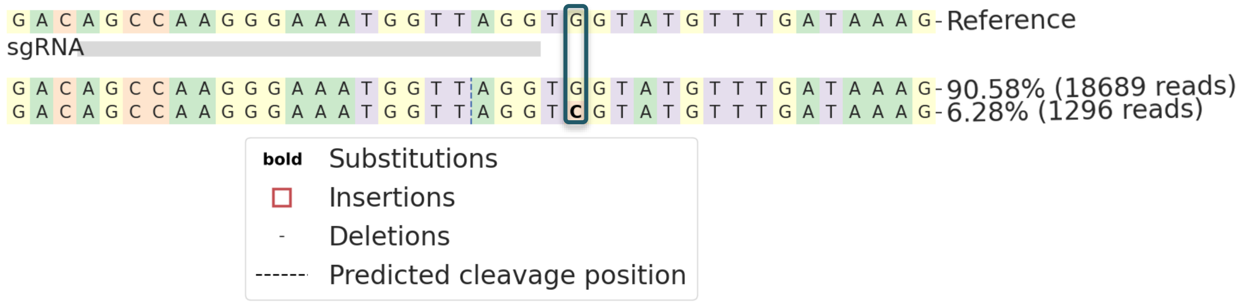


**Extended data 41:** Representative snapshot of CRISPResso2 analysis showing mutations at the target reverse transcriptase template sequence for *OsACC* locus using the pKb-PAiD PE vector.

(a) (b)

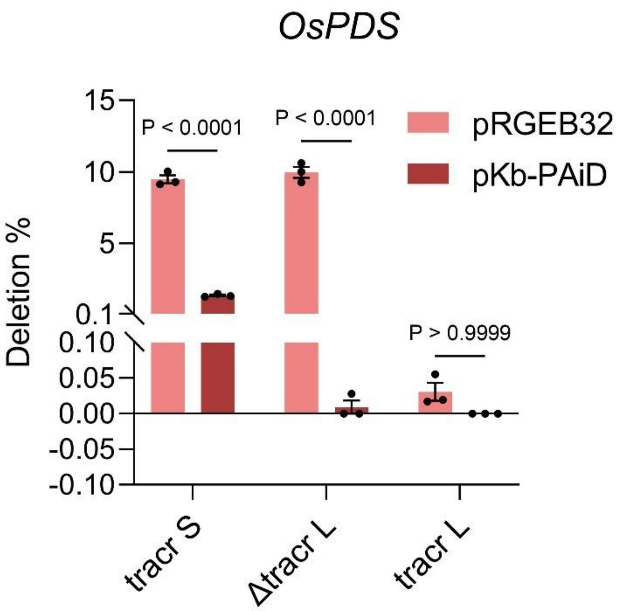


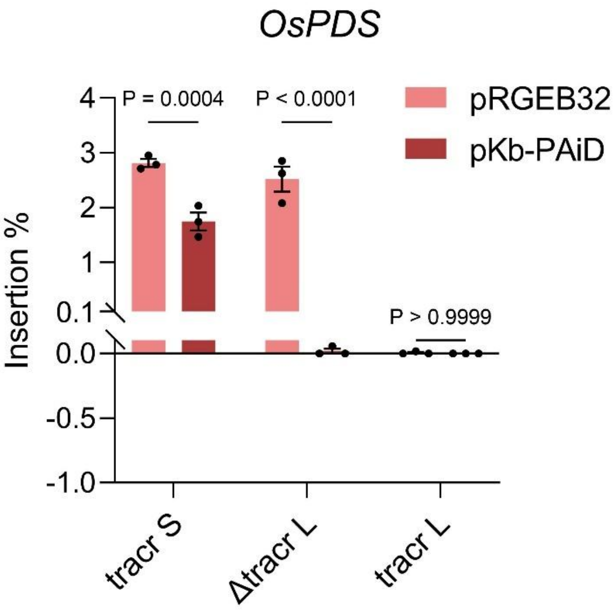


**Extended data 42:** Comparative study of three different tracrRNAs in combination with both Cas9 and PAiD for the *OsPDS* locus. **(a)** The bar graph presents the deletion percentage for both Cas9 and PAiD in combination with tracr S (conventional tracrRNA used for SpCas9), Δtracr L (truncated long tracrRNA of SpCas9) and tracr L (wildtype long tracrRNA of SpCas9) for the *OsPDS* locus. **(b)** The bar graph presents the insertion percentage for both Cas9 and PAiD in combination with tracr S (conventional tracrRNA used for SpCas9), Δtracr L (truncated long tracrRNA of SpCas9) and tracr L (wildtype long tracrRNA of SpCas9) for the *OsPDS* locus.

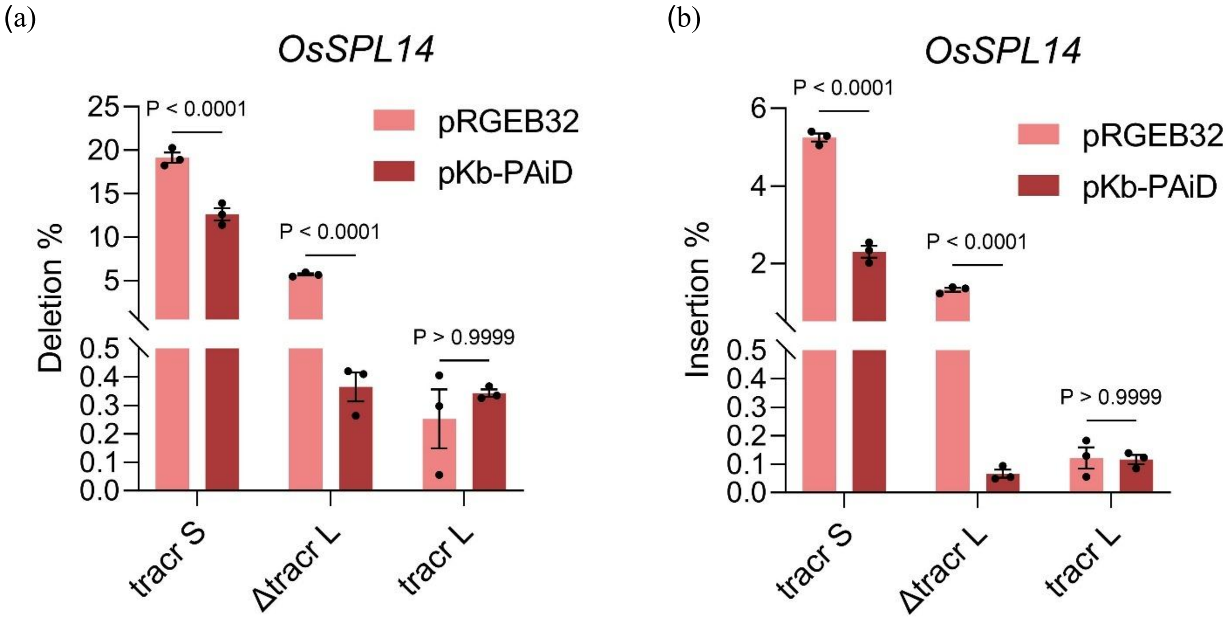


**Extended data 43:** Comparative study of three different tracrRNAs in combination with both Cas9 and PAiD for the *OsSPL14* locus. **(a)** The bar graph presents the deletion percentage for both Cas9 and PAiD in combination with tracr S (conventional tracrRNA used for SpCas9), Δtracr L (truncated long tracrRNA of SpCas9) and tracr L (wildtype long tracrRNA of SpCas9) for the *OsSPL14* locus. **(b)** The bar graph presents the insertion percentage for both Cas9 and PAiD in combination with tracr S (conventional tracrRNA used for SpCas9), Δtracr L (truncated long tracrRNA of SpCas9) and tracr L (wildtype long tracrRNA of SpCas9) for the *OsSPL14* locus.

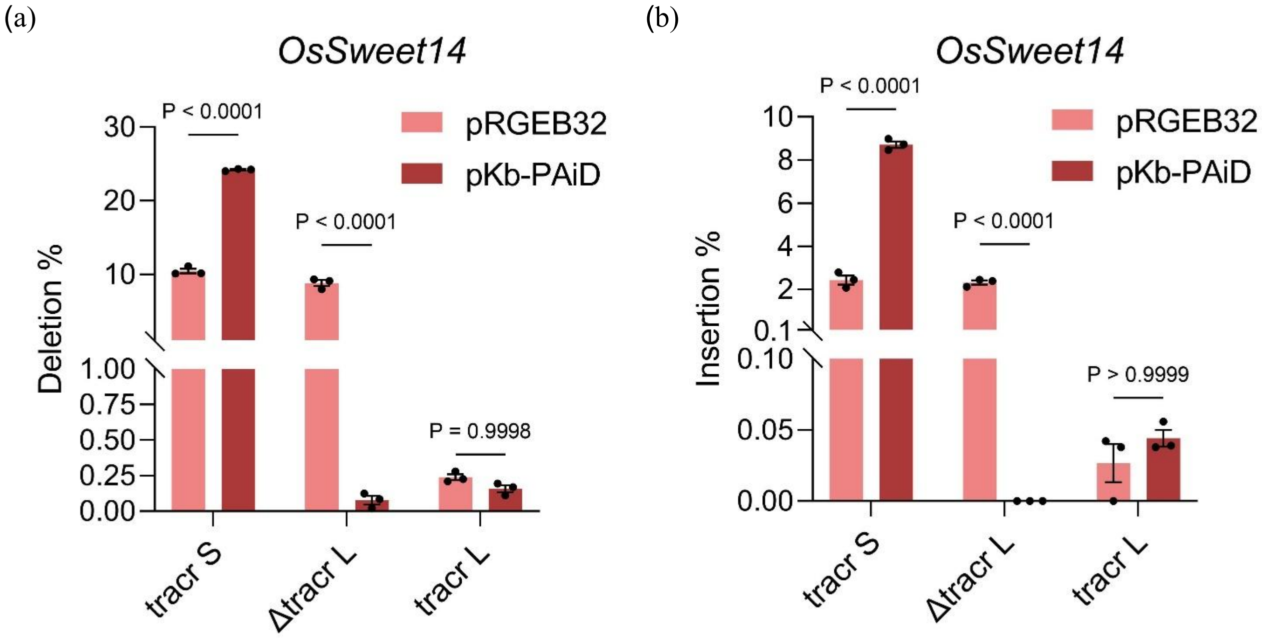


**Extended data 44:** Comparative study of three different tracrRNAs in combination with both Cas9 and PAiD for the *OsSweet14* locus. **(a)** The bar graph presents the deletion percentage for both Cas9 and PAiD in combination with tracr S (conventional tracrRNA used for SpCas9), Δtracr L (truncated long tracrRNA of SpCas9) and tracr L (wildtype long tracrRNA of SpCas9) for the *OsSweet14* promoter locus. **(b)** The bar graph presents the insertion percentage for both Cas9 and PAiD in combination with tracr S (conventional tracrRNA used for SpCas9), Δtracr L (truncated long tracrRNA of SpCas9) and tracr L (wildtype long tracrRNA of SpCas9) for the *OsSweet14* promoter locus.

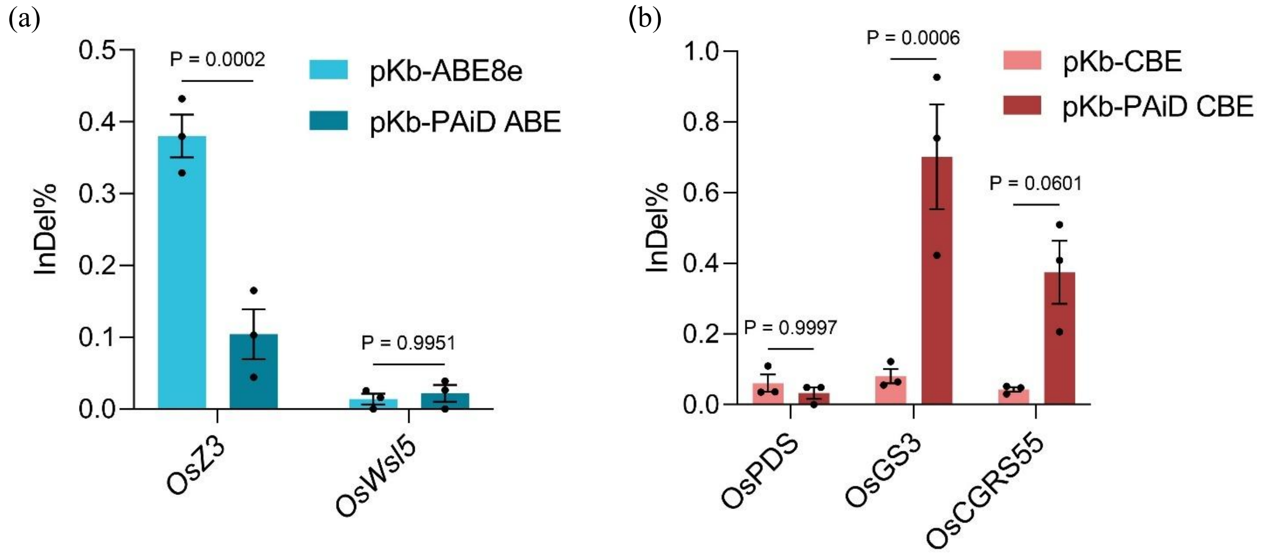


**Extended data 45:** Comparative analysis of unintended InDels in base editing experiments. (a) This bar diagram shows the comparative proportions of unwanted InDels in Adenine base editing experiment. In case of *OsZ3* the InDel percentage is significantly lower with PAiD compared to Cas9 where as in *OsWsl5*, both the PAiD and Cas9 showed similar percentage of unwanted InDels. (b) The bar diagram comprises the comparison of unwanted InDels in the Cytosine base editing experiment. In case of *OsPDS* and *OsCGRS55*, InDel percentage is almost similar with both Cas9 and PAiD, whereas for *OsGS3*, InDel percentage showed significant increase with PAiD as compared to Cas9.

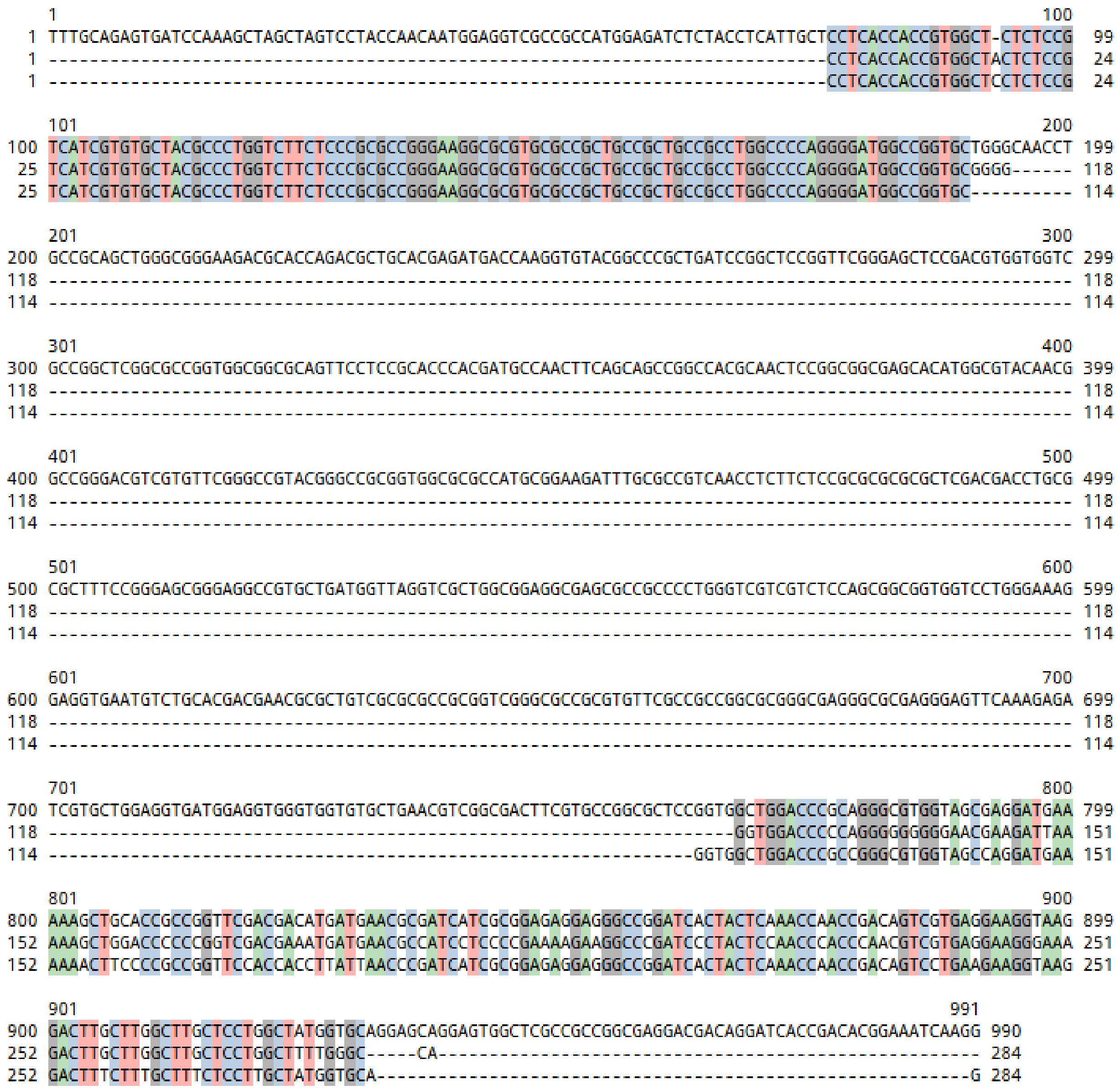

**Extended data 46:** Image depicts the CRISP-Id analysis from the Sanger sequencing data of a regenerated T0 plant for *cyp75b4* knockout induced by PAiD. The top strand denotes the control sequence and the lower two strands denote two alleles from the plant. The two alleles show different type of deletion patterns, indicating the plant as a biallelic heterozygous one.

### Supplemental Material and Methods

#### Plant material and growth condition

Fresh seeds of Kitaake (*Oryza sativa* L. subsp. japonica cultivar) were used for this study. Kitaake plants were maintained at ICAR–Central Rice Research Institute (ICAR-CRRI), Cuttack.

#### Vector construction for OC1-mediated Knock-out in rice

To generate pKb_PAiD (Supplementary Table 1), rice codon-optimized PAiD (Seq ID NO: 14) was synthesized from Twist Bioscience (USA). The synthesized PAiD nuclease coding sequence (KM#27) was flanked by XbaI (New England Biolabs, R0145S) and BstBI (New England Biolabs, R0519S) restriction sites. The rice codon optimized nuclease was excised from the KM#27 using BstBI and XbaI restriction enzymes and ligated into the same sites of pRGEB32 vector (Addgene Plasmid #63142) using T4 DNA ligase (New England Biolabs, M0202S) to generate pKb-PAiD (Supplementary Fig. 1). The final vector was confirmed through restriction digestion.

To generate the multiplexed editing cassette (Supplementary Table 1), three DNA fragments, Fragment 1 (128 bp), Fragment 2 (195 bp), and Fragment 3 (138 bp), were PCR-amplified using distinct primer pairs. All fragments were amplified from the plasmid pGTR (Addgene #63143) using Q5® High-Fidelity DNA Polymerase (New England Biolabs, M0491S). Specifically, Fragment 1 was amplified with primers 74 and 1086/Gr3R, Fragment 2 with primers 1085/Gr3F and 1088/Gr4R, and Fragment 3 with primers 1087/Gr4F and 75.

The three PCR products were assembled into a single cassette (PTG) using a Golden Gate assembly workflow. The assembled product was subsequently re-amplified with Q5® High-Fidelity DNA Polymerase using primers 76 and 77 to obtain sufficient material for downstream cloning. The purified amplicon was polyadenylated using Taq DNA Polymerase (New England Biolabs, M0273S) to facilitate TA cloning and subsequently ligated into the pGEM®-T Easy vector (Promega, A1360). Positive clones were verified by Sanger sequencing.

Following sequence confirmation, the final PTG cassette was excised and inserted into the pKb-PAiD and pRGEB32 via BsaI (New England Biolabs, R3733S) mediated restriction cloning using T4 DNA Ligase.

#### Preparation of tracrL based vectors

The constructs pKb-Cas9-tracrL, pKb-Cas9-ΔtracrL, and pKb-Cas9-tracrS, harbouring three different rice targets, *OsPDS*, *OsSPL14*, and the promoter region of *OsSWEET14* were used in this study. All vector construction details have been described in our previous report (Karmakar et al., 2024). To generate pKb-PAiD-tracr-L, pKb-PAiD-Δtracr-L, and pKb-PAiD-tracr-S using the same guide RNAs, the corresponding Cas9 containing vectors were independently digested with BstBI and SacI (New England Biolabs, R3156S) and used as backbone vectors. The synthesized PAiD fragment (KM#27) was digested with the same restriction enzymes and used as the insert. Ligation was performed using T4 DNA ligase, followed by transformation into E coli. Positive clones were confirmed by restriction digestion analysis.

##### Vector construction for PAiD based adenine base editor: pKb-PAiD-ABE

To create pKb-PAiD-ABE (Seq ID No. 3), two fragments were synthesized. The synthesized fragment KM#41 contains ABE8e monomer and a portion of rice codon-optimised PAiD having D>A mutation. The KM#41 was flanked by BstBI and MfeI (New England Biolabs, R3589S) restriction sites. Another synthesized product KM#44 contained OsU3 promoter, guide RNA cloning site and scaffold. The KM#44 was flanked by HindIII-HF (New England Biolabs, R3104S) and NheI (New England Biolabs, R3131S) restriction sites.

We first generated pUC19-PAiD. The pKb-PAiD (Supplementary Fig. 1) was digested using restriction enzyme SbfI-HF (New England Biolabs, R3642S) and EcoRI-HF (New England Biolabs, R3101S) to release PAiD. PAiD was ligated into the same sites in pUC19 (Addgene Plasmid #50005). Next, KM#41 was digested with BstBI and MfeI to release clone in the same sites into pUC19-PAiD for generating pUC19-ABE8e-nPAiD (n stands for nickase).

We, then, generated pG3H-KM#44. The KM#44 was digested using HindIII-HF and NheI-HF and ligated into the same sites in PE3max_pG3H (Chen et al., 2021) to generate PE3max-KM#44. Next, we digested pUC19-ABE8e-nPAiD with AvrII (New England Biolabs, R0174S) and SacI-HF to release ABE8e-nPAiD fragment. The released fragment was then ligated into same sites of PE3max-KM#44 to generate pKb-PAiD-ABE (Supplementary Fig. 3). The final vector was confirmed through restriction digestion.

Following a similar fashion, pKb-ABE8e was generated.

##### Vector construction for PAiD based cytosine base editor: pKb-PAiD-CBE

We synthesized KM#42 and KM#43 to generate pKb-PAiD-CBE (Seq. Id. No. 5). KM#42 contains A3A/Y130F cytidine deaminase and a portion of PAiD having D>A mutation. KM#43 contains uracil DNA glycosylase inhibitor (UGI) and linker. KM#42 was digested with BstBI and MfeI and ligated into same sites in pUC19-ABE8e-nPAiD to generate pUC19-A3A-nPAiD. One of the two SphI (New England

Biolabs, R3182S) sites in pUC19-A3A-nPAiD was removed through an intermediary step. KM#43 was digested by Sph1 and Xba1 and cloned in the same sites in pUC19-A3A-PAiD to generate pUC19-PAiD-CBE. Next, PAiD-CBE fragment was released from pUC19-PAiD-CBE by digesting with AvrII and SacI-HF and cloned into the same sites in PE3max-KM#44 to generate **pKb-PAiD-CBE** (Supplementary Fig. 5). Following a similar fashion, pKb-CBE was generated.

#### Vector construction for PAiD based prime editor: pKb-PAiD PE

pUC19 vector was digested with BamHI-HF (New England Biolabs, R3136S) and SphI and the 2666 bp was purified as a vector. KM#27, harbouring synthesized PAiD, was digested with BamHI and SphI and a 4077 bp was purified and ligated into the above generated vector to generate pUC19-KM27. For prime editing, we need H850A modification in the endonuclease. To achieve this, KM#45, harbouring the H850A mutation, was synthesized. KM#45 was digested with BsrGI (New England Biolabs, R3575S) and AgeI (New England Biolabs, R3552S), 445 bp purified, and ligated into pUC19-KM27 in the same sites for generating pUC19-KM27-H850A. Next, KM#27 was digested with BsrGI and BamHI-HF and a 3123 bp was purified. The fragment was then ligated into pUC19-KM27-H850A in BsrGI and BamHI-HF sites to generate KM27-H850A.

KM#46, harbouring a portion of MMLV reverse transcriptase pentamutant (D200N/L603W/T330P/T306K/W313F), was synthesized. A 1383 bp fragment was released from PE3max-pG3H (Chen et al., 2021) by digesting it by KpnI (New England Biolabs, R3142S) and SacI-HF and ligated to KM#46 to generate pUC57-mini-complete RT. pUC57-mini-complete RT was digested with SphI and SacI-HF to release a 2706 bp fragment. The fragment was then cloned in the same sites into pTWIST-KM27-H850A to generate pTWIST-PAiD-RT. pTWIST-PAiD-RT was digested with AvrII and SacI-HF and a 6520 bp band was purified. The fragment was then ligated to the same sites in PE3max-pG3H to generate pKb-PAiD PE.

#### Protoplast isolation, transfection of constructs

Rice protoplast isolation and transfection were carried out following our previously published protocol (Panda et al., 2024). Approximately 150–200 rice seedlings were grown in darkness at 30 °C for 10 days. Etiolated stems were then cut perpendicularly into 0.5–1 mm strips using sterile razor blades. The cut strips were transferred to a sterile 100 mL conical flask containing 10 mL of 0.6 M mannitol and incubated in the dark for 10 minutes. The mannitol solution was removed, and 10 mL of protoplast isolation buffer was added. The strips were incubated in the enzymatic digestion buffer for 5 hours at 25 °C in the dark with gentle shaking (40 rpm) and vacuum infiltration. The digestion mixture was filtered through a 40 μm nylon mesh, and the filtrate was centrifuged at 100 × g for 7 minutes. The supernatant was discarded, and the pellet was resuspended in W5 solution. After gentle mixing, the suspension was centrifuged again, and the pellet was resuspended in 6 mL of 0.55 M sucrose. Next, 2 mL of W5 buffer was carefully layered on top of the sucrose–protoplast suspension. The mixture was centrifuged at 100 × g for 30 minutes. A distinct band of protoplasts formed at the interface and was carefully collected into a fresh round-bottom tube. The isolated protoplasts were washed with 1–5 mL of W5 buffer and centrifuged at 100 × g for 5 minutes. Finally, the pellet was resuspended in MMG buffer to obtain a final concentration of $2 \times 10^6$ protoplasts/mL. For transfection, approximately 30 µg of plasmid DNA was mixed with the protoplast suspension ($2 \times 10^6$ protoplasts/mL) in MMG buffer. An equal volume of PEG solution was added, and the mixture was incubated for 20 minutes. The reaction was diluted with W5 buffer to a final volume of 1 mL and incubated for an additional 20 minutes. Subsequently, 900 µL of W5 buffer was added, and the mixture was gently inverted four times. The suspension was centrifuged at 100 × g for 5 minutes. For incubation, protoplasts were plated by adding 80 µL of 5% calf serum and 1 mL of W5 buffer per well in a 12-well plate. After 72 hours of incubation at 32 °C in the dark with gentle shaking (25 rpm), protoplasts were collected for genomic DNA isolation.

#### Genomic DNA isolation, amplification of the targets and deep sequencing

Genomic DNA was isolated from the collected protoplast samples using NucleoSpin Plant II, Mini kit (Catalogue No. 740770.250). For amplicon sequencing, we have designed Hi-TOM sequencing primers for all the target genes following the Hi-TOM Rapid Sequencing Service Protocol (Liu et al., 2018). The FASTA files of PCR amplicons were then demultiplexed and then analysed though CRISPResso22 (https://CRISPResso2.pinellolab.org/submission) (Clement et al., 2019). Editing efficiencies were calculated as the percentage of reads with mutations relative to total mapped reads.

#### Stable rice transformation

Transgenic Kitaake rice lines were generated using Agrobacterium-mediated transformation following the protocol described by (Molla et al., 2020). Mature embryo-derived calli were induced on N6-D callus induction medium and maintained for 14 days at 32 °C under cool white light. Binary vectors, pRGEB32-CYP M and pKb-PAiD-CYPM were introduced into *Agrobacterium tumefaciens* strain LBA4404. Transformed *Agrobacterium* cultures were grown at 28 °C for 2 days and then resuspended in liquid callus induction medium ($OD_{600}$ = 0.15) supplemented with 100 µM acetosyringone. Calli were co-cultivated with *Agrobacterium* for 72 hrs, washed with liquid CIM containing 200 mg/L Timentin, and transferred to selection medium supplemented with 200 mg/L Timentin and 50 mg/L Hygromycin. After 3 weeks of selection, surviving calli were moved to regeneration medium containing 200 mg/L Timentin. Regenerated shoots were transferred to the rooting medium and subsequently moved to the greenhouse for hardening and further growth.

##### Mutation analysis from plants

Genomic DNAs were extracted from leaf samples of T0 plant lines. The target loci were amplified through PCR using respective primer sets (Supplementary table 4), and the resulting amplicons were sent for Sanger sequencing. The Sanger sequencing data were analysed on CRISP-ID webtool (Dehairs et al., 2016).

##### Statistical Analysis

All experiments in this study were performed with three biological replicates for each target. Statistical analyses were carried out using GraphPad Prism (version 9.4.1). Data are presented as mean ± s.e.m. Multiple comparisons were analysed using one-way or two-way ANNOVA, as appropriate. For comparisons between two sample groups, Tukey's unpaired t-test was used.

#### Supplementary Tables

**Supplemental Table 1. Different vectors, polynucleotides, polypeptide sequences and their sequence IDs**

| Sl No. | Vector, polynucleotide, polypeptide Name | Seq. ID |
|---|---|---|
| 1 | pKb-PAiD<br>CaMV Poly(A) Signal-HPT-CaMV35s Promoter (enhanced)-OsU3 Promoter-crRNA cloning site-tracrRNA-PolyT-OsUbi10 Promoter-SV40 NLS-PAiD-Nucleoplasmin NLS-NOS T | Seq ID No.1 |
| 2 | pKb-PAiD-Δ*tracrL*<br>CaMV Poly(A) Signal-HPT-CaMV35s Promoter (enhanced)-OsU3 Promoter-crRNA cloning site-ΔtracrRNA L-PolyT-OsUbi10 Promoter-SV40 NLS-PAiD-Nucleoplasmin NLS-NOS T | Seq ID No.2 |
| 3 | pKb-PAiD-*tracrL*<br>CaMV Poly(A) Signal-HPT-CaMV35s Promoter (enhanced)-OsU3 Promoter-crRNA cloning site-tracrRNA L-PolyT-OsUbi10 Promoter-SV40 NLS-PAiD-Nucleoplasmin NLS-NOS T | Seq ID No.3 |
| 4 | pKb-PAiD-ABE<br>OsU3 Promoter-crRNA cloning site-tracrRNA-PolyT-ZmUbi Promoter-SV40 NLS (Bipartite)-Adenine Deaminase-Linker-PAiD D10A-Nucleoplsmin NLS-E9 terminator-CaMV35S Promoter (enhanced)-HPT-CaMV Poly(A) Signal | Seq ID No.4 |
| 5 | pKb-PAiD-CBE<br>OsU3 Promoter-crRNA cloning site-tracrRNA-PolyT-ZmUbi Promoter-Cytidine deaminase-SV40 NLS-PAiD D10A-Uridine glycosylase inhibitor-SV40 NLS-E9 terminator-CaMV35S Promoter (enhanced)-HPT-CaMV Poly(A) Signal | Seq ID No. 5 |
| 6 | pKb-PAiD PE<br>Composit Promoter-tGly-AmpR Promoter-AmpR-sgRNA2m-HDV ribozyme-PolyT HSPt-ZmUbi Promoter-SV40 NLS-PAiD H850A-Linker-M MLV RT-NLSc Myc-E9 terminator-CaMV35S Promoter (enhanced)-HPT-CaMV Poly(A) Signal | Seq ID No. 6 |
| 7 | pKb-PAiD M<br>CaMV Poly(A) Signal-HPT-CaMV35s Promoter (enhanced)-OsU3 Promoter-tRNA-crRNA1-tracrRNA-tRNA-crRNA2-tracrRNA-PolyT-OsUbi10 Promoter-SV40 NLS-PAiD-Nucleoplasmin NLS-NOS T | Seq ID No. 7 |
| 8 | Rice codon optimized PAiD (KM#27) | Seq ID No. 8 |
| 9 | Adenine deaminase | Seq ID No. 9 |
| 10 | Cytidine deaminase | Seq ID No. 10 |

| | | |
|---|---|---|
| 11 | Uridine Glycosylase Inhibitor | Seq ID No. 11 |
| 12 | SV40 NLS | Seq ID No. 12 |
| 13 | Nucleoplasmin NLS | Seq ID No. 13 |
| 14 | Rice codon optimized PAiD D10A | Seq ID No. 14 |
| 15 | Ric codon optimized PAiD H850A | Seq ID No. 15 |
| 16 | Prime editor linker | Seq ID No. 16 |
| 17 | M MLV RT | Seq ID No. 17 |
| 18 | Δ*tracrL* | Seq ID No. 18 |
| 19 | *tracrL* | Seq ID No. 19 |

**Supplementary Table 2**: **Target genes with locus ID and guide sequence**

| SL No | Gene name | Locus ID | Guide sequence |
|---|---|---|---|
| 1 | *OsSWEET11* | LOC_Os08g42350 | TTGGTGGTGTACAGTAGG |
| 2 | *OsSWEET14* | LOC_Os11g31190 | TATATAAACCCCCTCCAACC |
| 3 | *OsZ3* | LOC_Os03g05390 | TGGTCGACGGCACCCCGAAC |
| 4 | *OsSWEET14* | LOC_Os11g31190 | AGGGCATGCATGTCAGCAGC (guide1)<br>TATATAAACCCCCTCCAACC (guide2)<br>Used in multiplexed system |
| 5 | *CYP75B3* | LOC_Os10g17260 | TGCGGCAGGTTGCCCAGCAC (guide1)<br>ACTTCGTGCCGGCGCTCCGG (guide2)<br>Used in multiplexed system |
| 6 | *OsWSL5* | LOC_Os04g58780 | CACGAGCTCGACCTTCTCCA |
| 7 | *OsGS3* | Os03g0407400 | AGATCCAGGAGAGGTAGCTG |
| 8 | *OsCGRS55* | PRJNA1123306<br>NC_089038.1 | CCACCCCTCCATCTCCTCCA |
| 9 | *OsPDS* | LOC_Os03g08570 | ACATTGCCTGCACCCTTAAA<br>(for CBE experiment)<br>TCAAACCGGCTGAATTCTCC<br>(for alternative tracr experiment) |
| 10 | *OsSPL14* | OsKitaake08g2077<br>00 | CTCTTCTGTCAACCCAGCCA |
| 11 | *OsPEPC* | OsKitaake08g1319<br>00 | TGTAGGCATTGCGGAGGCGA |
| 12 | *OsALS* | LOC_Os02g30630 | CTCGAGCATCTTCTTGATGG |
| 13 | *OsACC* | LOC_Os05g22940 | AGCCAAGGGAAATGGTTAGG |

**Supplemental Table 3. List of primers used for cloning different guides**

| Primer name | Primer sequence | Purpose |
|---|---|---|
| 261 Oligo1 *OsSweet11* | ggcaAGTTTTGGTGGTGTACAGTAGG | Cloning of guide for *OsSweet11* for InVitro Protoplast validation |
| 262 Oligo2 *OsSweet11* | aaacCCTACTGTACACCACCAAAACT | |
| 753 Oligo1 *OsSweet14* | ggcaTATATAAACCCCCTCCAACC | Cloning of guide for *OsSweet14* for InVitro Protoplast validation |
| 754 Oligo2 *OsSweet14* | aaacGGTTGGAGGGGGTTTATATA | |
| 778 Oligo1 *OsZ3* | ggcaTGGTCGACGGCACCCCGAAC | Cloning of guide for *OsZ3* for InVitro Protoplast validation |
| 777 Oligo2 *OsZ3* | aaacGTTCGGGGTGCCGTCGACCA | |
| 909 Oligo1 *OsWSL5* | ggcaCACGAGCTCGACCTTCTCCA | Cloning of guide for *OsWsl5* for InVitro Protoplast validation |
| 910 Oligo2 *OsWsl5* | aaacTGGAGAAGGTCGAGCTCGTG | |
| a71 Oligo1 *OsGS3* | ggcaAGATCCAGGAGAGGTAGCTG | Cloning of guide for *OsGS3* for InVitro Protoplast validation |
| a72 Oligo2 *OsGS3* | aaacCAGCTACCTCTCCTGGATCT | |
| a73 Oligo1 *OsCGRS55* | ggcaCCACCCCTCCATCTCCTCCA | Cloning of guide for *OsCGRS55* for InVitro Protoplast validation |
| a74 Oligo2 *OsCGRS55* | aaacTGGAGGAGATGGAGGGGTGG | |
| a77 Oligo1 *OsPDS* | ggcaACATTGCCTGCACCCTTAAA | Cloning of guide for *OsPDS* for InVitro Protoplast validation for CBE |
| a78 Oligo2 *OsPDS* | aaacTTTAAGGGTGCAGGCAATGT | |
| 74-L5AD5 F | CGGGTCTCAGGCAGGATGGGCAGTCTGGGCAACAAAGCACCAGTGG | Cloning of multiplexed guide using pGTR system for *OsCYP75B4* and *OsSweet14* for InVitro Protoplast |
| 75-L3AD5 R | TAGGTCTCCAAACGGATGAGCGACAGCAAACAAAAAAAAAAGCACCGACTCG | |
| 76-S5AD5 F | CGGGTCTCAGGCAGGATGGGCAGTCTGGGCA | |

| | | |
|---|---|---|
| 77-S3AD5 R | TAGGTCTCCAAACGGATGAGCGACAGCAAAC | validation and stable plant transformation |
| 1085-CYP75B4 G1 F | TAGGTCTCCGTTGCCCAGCACgttttagagctagaa | Cloning of multiplexed guide using pGTR system for *OsCYP75B4* for InVitro Protoplast validation and stable plant transformation |
| 1086-CYP75B4 G1 R | CGGGTCTCACAACCTGCCGCAtgcaccagccggg | |
| 1087-CYP75B4 G2 F | TAGGTCTCCCCGGCGCTCCGGgttttagagctagaa | |
| 1088-CYP75B4 G2 R | CGGGTCTCACCGGCACGAAGTtgcaccagccggg | |
| Gr3F | TAGGTCTCCCATGTCAGCAGCgttttagagctagaa | Cloning of multiplexed guide using pGTR system for *OsSweet14* for InVitro Protoplast validation and stable plant transformation |
| Gr3R | CGGGTCTCACATGCATGCCCTtgcaccagccggg | |
| Gr4F | TAGGTCTCCCCCCCTCCAACCgttttagagctagaa | |
| Gr4R | CGGGTCTCAGGGGTTTATATAtgcaccagccggg | |
| 751-Oligo1 OsPDS | ggcaTCAAACCGGCTGAATTCTCC | Guide for *OsPDS* for in vivo validation in protoplast |
| 752-Oligo2 OsPDS | aaacGGAGAATTCAGCCGGTTTGA | |
| 291-Oligo1 OsSPL14 | ggcaCTCTTCTGTCAACCCAGCCA | Cloning of guide for *OsSPL14* for in vivo validation in protoplast |
| 292-Oligo2 OsSPL14 | aaacTGGCTGGGTTGACAGAAGAG | |

**Supplementary Table 4: List of primers used for screening of mutant plants**

| Primer name | Primer sequence | Purpose |
|---|---|---|
| 1083 F | TTTGCAGAGTGATCCAAAGCTAG | Screening of *OsCYP75B4* mutant lines |
| 1084 R | CCTTGATTTCCGTGTCGGTGA | |

**Supplementary Table 5: List of primers used for Deep amplicon sequencing (1st round)**

| Primer name | Primer sequence | Purpose |
|---|---|---|
| a33-F | ggagtgagtacggtgtgcTGGAGAGAGGGACAGATCTAGA | 1st round amplification for *OsSWEET11* |
| a34-R | gagttggatgctggatggCAGTGAGAAGGTTAGGAAGAGGA | |
| 1522-F | ggagtgagtacggtgtgcAGTTTGTGTGTGCAGCTATATTG | 1st round amplification for *OsSWEET14* |
| 1523-R | gagttggatgctggatggGGGAGGAGATCAATGAGGCACAGT | |
| b18-F | ggagtgagtacggtgtgcCTTTGTCTTTCAGTACTACTGCTCG | 1st round amplification for *OsZ3* |
| b19-R | gagttggatgctggatggCTTATTGATCCAAAGGGTTCGTTC | |
| a37-F | ggagtgagtacggtgtgcGAGCAGCCAGAGCCTTCTAC | 1st round amplification for *OsWSL5* |
| a38-R | gagttggatgctggatggAGTCAACCCCGTCAACCTC | |
| a79-F | ggagtgagtacggtgtgcCCATGCTCACACTGTTTTGTCGT | 1st round amplification for *OsPDS* |
| a80-R | gagttggatgctggatggACGTTGCTAGTAATATACCCCCTA | |
| a69-F | ggagtgagtacggtgtgcCCCACAAAACCATCAACTTGTTAAT | 1st round amplification for *OsGS3* |
| a70-R | gagttggatgctggatggGACACGGACTCTTCGTTAACGCC | |
| a75-F | ggagtgagtacggtgtgcGCTTGCATTATTAATTGCCGACG | 1st round amplification for *OsCGRS55* |
| a76-R | gagttggatgctggatggACAACGACCTACCTCCCTAGA | |
| a3-F | ggagtgagtacggtgtgcGAAACTGAGGGCCAACTGTG | 1st round amplification for *OsPEPC* |
| a4-R | gagttggatgctggatggCGGCTTAGACCAGTCCATGA | |
| a13-F | ggagtgagtacggtgtgcGTAACAACACCGTTGGACCC | 1st round amplification for *OsACC* |
| a14-R | gagttggatgctggatggCCAGCACGAGGAACAGATTG | |

| | | |
|---|---|---|
| a15-F | ggagtgagtacggtgtgcGAGTTGGCATTGATCCGCAT | 1st round amplification for *OsALS* |
| a16-R | gagttggatgctggatggATCATGTCCTTGAATGCGCC | |

*Red marks indicate the bridge sequences to be used in the second round PCR

**Supplementary Table 6: List of primers used for Deep amplicon sequencing (2nd round)**

| Primer name | Primer sequence | Purpose |
|---|---|---|
| a49-F1 | ACACTCTTTCCCTACACGACGCTCTTCCGATCTgcttGCGTtggagtgagtacggtgtgc | 2nd round barcoding amplification |
| a50-F2 | ACACTCTTTCCCTACACGACGCTCTTCCGATCTgcttGTAGtggagtgagtacggtgtgc | |
| a51-F3 | ACACTCTTTCCCTACACGACGCTCTTCCGATCTgcttACGCtggagtgagtacggtgtgc | |
| a52-F4 | ACACTCTTTCCCTACACGACGCTCTTCCGATCTgcttCTCGtggagtgagtacggtgtgc | |
| a53-F5 | ACACTCTTTCCCTACACGACGCTCTTCCGATCTgcttGCTCtggagtgagtacggtgtgc | |
| a54-F6 | ACACTCTTTCCCTACACGACGCTCTTCCGATCTgcttAGTCtggagtgagtacggtgtgc | |
| a55-F7 | ACACTCTTTCCCTACACGACGCTCTTCCGATCTgcttCGACtggagtgagtacggtgtgc | |
| a56-F8 | ACACTCTTTCCCTACACGACGCTCTTCCGATCTgcttGATGtggagtgagtacggtgtgc | |
| a57-F9 | ACACTCTTTCCCTACACGACGCTCTTCCGATCTgcttATACtggagtgagtacggtgtgc | |
| a58-F10 | ACACTCTTTCCCTACACGACGCTCTTCCGATCTgcttCACAtggagtgagtacggtgtgc | |
| a59-F11 | ACACTCTTTCCCTACACGACGCTCTTCCGATCTgcttGTGCtggagtgagtacggtgtgc | |
| a60-F12 | ACACTCTTTCCCTACACGACGCTCTTCCGATCTgcttACTAtggagtgagtacggtgtgc | |
| a61-RA | GACTGGAGTTCAGACGTGTGCTCTTCCGATCTctgtGCGTtgagttggatgctggatgg | |
| a62-RB | GACTGGAGTTCAGACGTGTGCTCTTCCGATCTctgtGTAGtgagttggatgctggatgg | |
| a63-RC | GACTGGAGTTCAGACGTGTGCTCTTCCGATCTctgtACGCtgagttggatgctggatgg | |
| a64-RD | GACTGGAGTTCAGACGTGTGCTCTTCCGATCTctgtCTCGtgagttggatgctggatgg | |

| | | |
|---|---|---|
| a65-RE | GACTGGAGTTCAGACGTGTGCTCTTCCGATCTctgtGCTCtgagttggatgctggatgg | 2nd round barcoding amplification |
| a66-RF | GACTGGAGTTCAGACGTGTGCTCTTCCGATCTctgtAGTCtgagttggatgctggatgg | |
| a67-RG | GACTGGAGTTCAGACGTGTGCTCTTCCGATCTctgtCGACtgagttggatgctggatgg | |
| a68-RH | GACTGGAGTTCAGACGTGTGCTCTTCCGATCTctgtGATGtgagttggatgctggatgg | |

*Red marks indicate the bridge sequences from the first round PCR template.

#### Supplementary Sequences

Polynucleotides

Seq ID No. 1

(CaMV Poly(A) Signal-HPT-CaMV35s Promoter (enhanced)-OsU3 Promoter-crRNA cloning site-tracrRNA-PolyT-OsUbi10 Promoter-SV40 NLS-PAiD-Nucleoplasmin NLS-NOS T)

ccgaattaattcgggggatctggattttagtactggattttggttttaggaattagaaattttattgatagaagtattttacaaatacaaatacatactaagggtttc
ttatatgctcaacacatgagcgaaaccctataggaaccctaattcccttatctgggaactactcacacattattatggagaaactcgagcttgtcgatcgaca
gatcccggtcggcatctactctatttctttgccctcggacgagtgctggggcgtcggtttccactatcggcgagtacttctacacagccatcggtccagacg
gccgcgcttctgcgggcgatttgtgtacgcccgacagtcccggctccggatcggacgattgcgtcgcatcgaccctgcgcccaagctgcatcatcgaaa
ttgccgtcaaccaagctctgatagagttggtcaagaccaatgcggagcatatacgcccggagtcgtggcgatcctgcaagctccggatgcctccgctcg
aagtagcgcgtctgctgctccatacaagccaaccacggcctccagaagaagatgttggcgacctcgtattgggaatccccgaacatcgcctcgctccag
tcaatgaccgctgttatgcggccattgtccgtcaggacattgttggagccgaaatccgcgtgcacgaggtgccggacttcggggcagtcctcggcccaa
agcatcagctcatcgagagcctgcgcgacggacgcactgacggtgtcgtccatcacagtttgccagtgatacacatggggatcagcaatcgcgcatatg
aaatcacgccatgtagtgtattgaccgattccttgcggtccgaatgggccgaacccgctcgtctggctaagatcggccgcagcgatcgcatccatagcct
ccgcgaccggttgtagaacagcgggcagttcggtttcaggcaggtcttgcaacgtgacaccctgtgaacggcgggagatgcaataggtcaggctctcg
ctaaactccccaatgtcaagcacttccggaatcgggagcgcggccgatgcaaagtgccgataaacataacgatctttgtagaaaccatcggcgcagctat
ttacccgcaggacatatccacgccctcctacatcgaagctgaaagcacgagattcttcgccctccgagagctgcatcaggtcggagacgctgtcgaactt
ttcgatcagaaacttctcgacagacgtcgcggtgagttcaggctttttcatatctcattgccccccggatctgcgaaagctcgagagagatagatttgtagag
agagactggtgatttcagcgtgtcctctccaaatgaaatgaacttccttatatagaggaagggtcttgcgaaggatagtgggattgtgcgtcatcccttacgt
cagtggagatatcacatcaatccacttgctttgaagacgtggttggaacgtcttctttttccacgatgctcctcgtgggtgggggtccatctttgggaccactg
tcggcagaggcatcttgaacgatagcctttcctttatcgcaatgatggcatttgtaggtgccaccttccttttctactgtccttttgatgaagtgacagatagctg
ggcaatggaatccgaggaggtttcccgatattaccctttgttgaaaagtctcaatagccctttggtcttctgagactgtatctttgatattcttggagtagacga
gagtgtcgtgctccaccatgttcacatcaatccacttgctttgaagacgtggttggaacgtcttctttttccacgatgctcctcgtgggtgggggtccatctttg
ggaccactgtcggcagaggcatcttgaacgatagcctttcctttatcgcaatgatggcatttgtaggtgccaccttccttttctactgtccttttgatgaagtga
cagatagctgggcaatggaatccgaggaggtttcccgatattaccctttgttgaaaagtctcaatagccctttggtcttctgagactgtatctttgatattcttgg
agtagacgagagtgtcgtgctccaccatgttggcaagctgctctagccaatacgcaaaccgcctctccccgcgcgttggccgattcattaatgcagctgg
cacgacaggtttcccgactggaaagcgggcagtgagcgcaacgcaattaatgtgagttagctcactcattaggcaccccaggctttacactttatgcttcc
ggctcgtatgttgtgtggaattgtgagcggataacaatttcacacaggaaacagctatgacatgattacgccaagcttaaggaatctttaaacatacgaaca
gatcacttaaagttcttctgaagcaacttaaagttatcaggcatgcatggatcttggaggaatcagatgtgcagtcagggaccatagcacaagacaggcgt
cttctactggtgctaccagcaaatgctggaagccgggaacactgggtacgttggaaaccacgtgatgtgaagaagtaagataaactgtaggagaaaagc
atttcgtagtgggccatgaagcctttcaggacatgtattgcagtatgggccggcccattacgcaattggacgacaacaaagactagtattagtaccacctcg
gctatccacatagatcaaagctgatttaaaagagttgtgcagatgatccgtggcaggagaccgaggtctcggttttagagctagaaatagcaagttaaaat
aaggctagtccgttatcaacttgaaaaagtggcaccgagtcggtgcttttttgttttagagctagaaatagcaagttaaaataaggctagtccgtttttagcgc
gtgcatgcctgcaggtccacaaattcgggtcaaggcggaagccagcgcgccaccccacgtcagcaaatacggaggcgcggggttgacggcgtcacc
cggtcctaacggcgaccaacaaaccagccagaagaaattacagtaaaaaaaaagtaaattgcactttgatccaccttttattacctaagtctcaatttggatc
acccttaaacctatcttttcaatttgggccgggttgtggtttggactaccatgaacaacttttcgtcatgtctaacttccctttcagcaaacatatgaaccatatat
agaggagatcggccgtatactagagctgatgtgtttaaggtcgttgattgcacgagaaaaaaaaatccaaatcgcaacaatagcaaatttatctggttcaaa
gtgaaaagatatgtttaaaggtagtccaaagtaaaacttatagataataaaatgtggtccaaagcgtaattcactcaaaaaaaatcaacgagacgtgtacca
aacggagacaaacggcatcttctcgaaatttcccaaccgctcgctcgcccgcctcgtcttcccggaaaccgcggtggtttcagcgtggcggattctccaa
gcagacggagacgtcacggcacgggactcctcccaccacccaaccgccataaataccagcccccctcatctcctctcctcgcatcagctccaccccccga
aaaatttctccccaatctcgcgaggctctcgtcgtcgaatcgaatcctctcgcgtcctcaaggtacgctgcttctcctctcctcgcttcgtttcgattcgatttcg
gacgggtgaggttgttttgttgctagatccgattggtggttagggttgtcgatgtgattatcgtgagatgtttaggggttgtagatctgatggttgtgatttgggc
acggttggttcgataggtggaatcgtggttaggttttgggattggatgttggttctgatgattgggggaatttttacggttagatgaattgttggatgattcgat
tggggaaatcggtgtagatctgttggggaattgtggaactagtcatgcctgagtgattggtgcgatttgtagcgtgttccatcttgtaggccttgttgcgagca
tgttcagatctactgttccgctcttgattgagttattggtgccatgggttggtgcaaacacaggctttaatatgttatatctgttttgtgtttgatgtagatctgtagg
gtagttcttcttagacatggttcaattatgtagcttgtgcgtttcgatttgatttcatatgttcacagattagataatgatgaactcttttaattaattgtcaatggtaa

ataggaagtcttgtcgctatatctgtcataatgatctcatgttactatctgccagtaatttatgctaagaactatattagaatatcatgttacaatctgtagtaatatc
atgttacaatctgtagttcatctatataatctattgtggtaatttctttttactatctgtgtgaagattattgccactagttcattctacttatttctgaagttcaggatac
gtgtgctgttactacctatctgaatacatgtgtgatgtgcctgttactatctttttgaatacatgtatgttctgttggaatatgtttgctgtttgatccgttgttgtgtcc
ttaatcttgtgctagttcttaccctatctgtttggtgattatttcttgcagatagttatcaacaagtttgtacaaaaaagcaggcttCGAAGGCCCTAG
GAAGCCACCATGGATTATAAAGATCACGACGGCGACTACAAAGATCATGACATCGATTACA
AGGACGACGACGACAAGATGAAACGTACAGCTGATGGCAGCGAATTTGAATCGCCGAAGAA
GAAGCGGAAGGTGATGAAAAAGCCTTACTCCATCGGTCTTGACATCGGTACTAACAGCGTG
GGCTGGGCCGTTATTACTGATGATTACAAGGTGCCTGCAAAGAAGATGAAGGTGCTCGGCA
ATACGGACAGATCACACATCAAGAAGAATCTCATTGGTGCGCTACTTTTTGACGCCGGGAAC
ACTGCTGAGGATAGGCGCCTGAAGAGAACGGCGCGGCGCCGCTATACAAGACGGAGGAATA
GAATACTGTATTTGCAGGAAATCTTCGCAGAAGAAATGAACAAGATTGATGAGTCCTTCTTC
CACCGGCTCGACGACAGCTTTCTCGTGCCCGAGGATAAAAGGGGCTCCAAGTATCCAATATT
TGCTACTCTGCAAGAAGAGAAGGAGTACCACAAGCAGTTCCCTACCATCTACCATCTCCGGA
AACAATTGGCTGAATCTAATGAGAAGGCAGATCTGCGATTGGTATATCTTGCTCTGGCTCAC
ATGATCAAGTACAGAGGGCATTTTCTCATTGACGATCCCAAATTCAAAGTACAAAACAATGA
TATACAGGGTCTGTTTGAGAAATTTGTTGAAGAATACGACAATGTGCAGGAAACATCTTTGT
CTAAGATAAAACTCAATGTCACAGAAATACTTACCGCCAAGATTCCGAAAAGTGAGAAGCA
AGAGCAGCTTCTCAAGAACTACCCATCGGAAAAGAAGAACACGTTATTTGGGAACTTAATC
GGCCTTGCGTTGGGCCTCACACCAAATTTCAAGACTAATTTTAGCTTGGAAAATGACGCTAA
ACTGCAAATCTCAAGCGAAAGTTATGAGGAGGACCTCGGCTCCCTCCTTGCACTAATTGGAG
AGAACTTCATTGAGCTGTTCTCCGCGGTCAAGAACTTGAGTGATGGGATTCTCCTTGCTGGG
ATAGTTTCAGATGAATCACCCCATGCTCCCCTCTCCACCAAAATGGTTATAAGGTTCAAGGA
GCACGAGGAAGACCTCGCCGCACTCAAGCATTTCATTAAGGCCAATTTGCCGGAGAAGTAC
GATGAGGTTTTTTCAGATGACTCAAAGAACGGCTACGCCGGGTACGTCGGTGTCGATAGTAA
GGTTCGCAAAAGGAACGGGAAATTAGCAACCGAGGAGGAGTTCTACAAATACCTCAAGGAT
ATTTTGAACAACGTGAAAGGCGCCGATTATTTCTTGGAGAAAATTAAACGGGAGGATTTACT
CCGCAAACAGCGCACATTCGACAATGGGACTATCCCATATCAGGTTCATTTAGAAGAGATGA
AGGCCATTTTGCAAAACCAAGGCGAGTATTACCCCTTTCTTAAAGAGAACAAGGAGAAGAT
TCAACAGATCCTGACATTCAGAATCCCGTACTACGTAGGACCTCTGGCACGCGGCAACCGCG
ACTTCGCGTGGTTGACGCGCAACTCAGATCAGGCGATTCGACCATGGAACTTTGAGGAGGTC
GTGGATAAGGCCTCCTCCGCGGAGGACTTCATCAACAAAATGACAAACTATGACCTCTATCT
ACCAGAGGAAAAGGTCCTTCCCAAGCATTCGCTCTTGTACGAGACATTCGCTGTCTACAACG
AGCTGACCAAAGTTAAGTTTATCGCTGAAGGACTGCGTGACTACCAATTCCTAGACTCCGGT
CAAAAGAAACAGATAGTTAATCAGCTGTTTAAAGAAAAGAGAAAAGTGACTGAGAAAGAT
ATTATCCATTACCTCCACAACGTGGACGGTTATGATGGGATCGAATTAAAAGGAATCGAGAA
GCAGTTTAATGCTAGTCTGTCGACGTATCATGATCTCCTAAAAATCATTAAGGACAAGGAGT
TTATGGACGATCCTAAGAACGAGGAGATCCTCGAGAACATCGTCCATACACTCACGATATTC
GAGGACCGCGAGATGATTAAGCAGAGGTTGGCTCAGTACGACTCTCTCTTTGATGAGAAAGT
CATCAAAGCTCTAACCCGTCGCCACTATACTGGCTGGGGCAAGTTATCCGCAAAGCTGATAA
ATGGCATCTGTGATAAGCAAACAAATAAAACGATCCTGGACTTTCTTATTGACGACGACAAG
ATTAACAGAAACTTCATGCAGCTTATCAACGACGACGGCCTTTCTTTCAAGGATATAATTCA
AAAGGCGCAGGTGGTCGGCAAGATCGACGATGTGAAGCAAGTTGTGCAGGAGCTTCCAGGT
TCTCCTGCGATTAAAAAGGGTATTTTGCAGTCGATAAAGATTGTTGATGAGTTGGTCAAGGT
CATGGGCCACGCCCCTGAGTCCATTGTTATCGAGATGGCCAGGGAGAATCAGACTACCGCCA
GGGGCAAGAAAAACTCCCAGCAGAGATACAAGCGCATTGAGGACGCCTTAAAAAATCTAGC
GCCGGGGTTGGATTCTAATATCCTCAAGGAAAACCCAACAGATAATATCCAGTTACAAAAC
GACCGCCTCTTCCTCTACTATCTTCAAAATGGTAAGGACATGTATACCGGTGAAGCCCTCGA
CATAAATCAACTCAGCAATTATGATATAGATCATATTGTGCCGCAGGCGTTTATAAAGGACG
ATAGTCTTGATAACAGGGTGCTTACTAGCAGCAAGGATAACAGGGGAAAGTCTGACAATGT
GCCTTCTATAGAAGTTGTTCAGAAGCGTAAGGCCTTCTGGCAGCAACTTCTGGACTCGAAAT
TGATAAGCGAGCGAAAGTTCAATAACCTCACCAAGGCAGAGCGCGGTGGACTCGACGAACG

TGACAAAGTTGGGTTCATTAAGAGGCAGTTGGTTGAAACACGGCAGATTACAAAACATGTC
GCTCAGATACTCGATGCCCGGTTCAACACGGAGGTAAACGAGAAGAACCAGAAGATAAGAA
AGGTAAAAATTATTACTCTGAAATCCAACCTCGTTTCAAATTTCAGGAAAGAGTTCGGCCTG
TACAAGGTCCGTGAGATAAATGACTACCACCACGCCCACGATGCATACCTGAATGCAGTCGT
GGCGAAAGCGATCCTGAAGAAGTACCCTAAACTTGAACCGGAGTTCGTGTATGGAGACTAC
CAGAAGTACGACCTTAAGAGGTACATCTCACGCTCCAAAGATCCAAAGGAAATAGAGAAAG
CAACAGAAAAGTACTTCTTTTATAGCAATTTGCTGAATTTTTTCAAGGAAGAAGTGCACTAC
GCTGATGGAACCATCATCAAGAGGGAGAATATAGAGTACTCTAAGGATACTGGCGAGATAG
CATGGAATAAAGAAAAGGATTTCGCCACTGTACGCAAGGTACTGAGCTGCCCTCAGGTCAA
TATCGTGAAAAAGACCGAGGTTCAAACTGGCGGCTTCTCCAAGGAGTCAATCCTCCCGAAG
AGAAATTCTGACAAGTTAATCGCCCGAAAAAAGGACTGGGACCCGAAGAAGTATGGAGGCT
TCGATTCACCGACGGTGGCATATTCCGTGCTGGTGGTTGCCAAGGTGGAGAAGGGCAAGTCA
AAAAAGCTCAAGTCAGTAAAGGAACTTGTCGGAATCACGATCATGGAGCGGTCCTCGTTCG
AGAAGGACCCGGTCGACTTCCTGGAAGCAAAAGGGTACAAGGAGGTGAGAAAAGATTTGAT
TATCAAGCTCCCCAAGTATTCTCTTTTCGAGCTGGAAAATGGACGGAAGCGCATGCTCGCGT
CGGCCGGCGAGCTTCAGAAAGGAAATGAACTGGCCCTCCCTTCAAAATATGTCAACTTCTTA
TATCTGGCAAGTCATTATGAAAAACTGAAAGGGAGTCCGGAGGATAATGAACAAAAACAAC
TGTTTGTCGAGCAGCACAAACATTACCTCGATGAGATTATTGAGCAAATCTCCGAGTTCTCC
AAGAGGGTCATTCTCGCGGACGCAAATCTCGACAAGGTATTATCTGCCTACAATAAACATAG
AGATAAGCCAATTAGGGAACAGGCGGAGAACATAATCCACCTCTTCACCCTCACGAACCTC
GGCGCCCCAGCCGCGTTTAAGTACTTTGACACCACCATCGACAGGAAACGATACACTTCTAC
AAAAGAAGTGCTCGATGCCACGCTGATCCATCAGAGCATTACAGGTCTTTATGAAACCCGTA
TAGATCTCAGCCAACTCGGAGGGGACAAGCGGCCGGCGGCCACCAAGAAAGCTGGCCAAGC
CAAGAAGAAGAAATGAGCTCAGATAATctagacccagctttcttgtacaaagtggttgataacagcgactacaaggatgacga
tgacaaggcttagagctcgaatttccccgatcgttcaaacatttggcaataaagtttcttaagattgaatcctgttgccggtcttgcgatgattatcatataatttc
tgttgaattacgttaagcatgtaataattaacatgtaatgcatgacgttatttatgagatgggtttttatgattagagtcccgcaattatacatttaatacgcgatag
aaaacaaaatatagcgcgcaaactaggataaattatcgcgcgcggtgtcatctatgttactagatcgggaattcact

Seq ID No. 2

(CaMV Poly(A) Signal-HPT-CaMV35s Promoter (enhanced)-OsU3 Promoter-crRNA cloning site-ΔtracrRNA L-PolyT-OsUbi10 Promoter-SV40 NLS-PAiD-Nucleoplasmin NLS-NOS T)

ttcggggatctggattttagtactggattttggttttaggaattagaaattttattgatagaagtattttacaaatacaaatacatactaagggtttcttatatgctc
aacacatgagcgaaaccctataggaaccctaattcccttatctgggaactactcacacattattatggagaaactcgagcttgtcgatcgacagatcccggt
cggcatctactctatttctttgccctcggacgagtgctggggcgtcggtttccactatcggcgagtacttctacacagccatcggtccagacggccgcgctt
ctgcgggcgatttgtgtacgcccgacagtcccggctccggatcggacgattgcgtcgcatcgaccctgcgcccaagctgcatcatcgaaattgccgtca
accaagctctgatagagttggtcaagaccaatgcggagcatatacgcccggagtcgtggcgatcctgcaagctccggatgcctccgctcgaagtagcg
cgtctgctgctccatacaagccaaccacggcctccagaagaagatgttggcgacctcgtattgggaatccccgaacatcgcctcgctccagtcaatgacc
gctgttatgcggccattgtccgtcaggacattgttggagccgaaatccgcgtgcacgaggtgccggacttcggggcagtcctcggcccaaagcatcagc
tcatcgagagcctgcgcgacggacgcactgacggtgtcgtccatcacagtttgccagtgatacacatggggatcagcaatcgcgcatatgaaatcacgc
catgtagtgtattgaccgattccttgcggtccgaatgggccgaacccgctcgtctggctaagatcggccgcagcgatcgcatccatagcctccgcgaccg
gttgtagaacagcgggcagttcggtttcaggcaggtcttgcaacgtgacaccctgtgaacggcgggagatgcaataggtcaggctctcgctaaactccc
caatgtcaagcacttccggaatcgggagcgcggccgatgcaaagtgccgataaacataacgatctttgtagaaaccatcggcgcagctatttacccgca
ggacatatccacgccctcctacatcgaagctgaaagcacgagattcttcgccctccgagagctgcatcaggtcggagacgctgtcgaacttttcgatcag
aaacttctcgacagacgtcgcggtgagttcaggctttttcatatctcattgccccccggatctgcgaaagctcgagagagatagatttgtagagagagactg

gtgatttcagcgtgtcctctccaaatgaaatgaacttccttatatagaggaagggtcttgcgaaggatagtgggattgtgcgtcatcccttacgtcagtggag
atatcacatcaatccacttgctttgaagacgtggttggaacgtcttctttttccacgatgctcctcgtgggtgggggtccatctttgggaccactgtcggcaga
ggcatcttgaacgatagcctttcctttatcgcaatgatggcatttgtaggtgccaccttccttttctactgtccttttgatgaagtgacagatagctgggcaatgg
aatccgaggaggtttcccgatattaccctttgttgaaaagtctcaatagccctttggtcttctgagactgtatctttgatattcttggagtagacgagagtgtcgt
gctccaccatgttcacatcaatccacttgctttgaagacgtggttggaacgtcttctttttccacgatgctcctcgtgggtgggggtccatctttgggaccact
gtcggcagaggcatcttgaacgatagcctttcctttatcgcaatgatggcatttgtaggtgccaccttccttttctactgtccttttgatgaagtgacagatagct
gggcaatggaatccgaggaggtttcccgatattaccctttgttgaaaagtctcaatagccctttggtcttctgagactgtatctttgatattcttggagtagacg
agagtgtcgtgctccaccatgttggcaagctgctctagccaatacgcaaaccgcctctccccgcgcgttggccgattcattaatgcagctggcacgacag
gtttcccgactggaaagcgggcagtgagcgcaacgcaattaatgtgagttagctcactcattaggcaccccaggctttacactttatgcttccggctcgtat
gttgtgtggaattgtgagcggataacaatttcacacaggaaacagctatgacatgattacgccaagcttaaggaatctttaaacatacgaacagatcactta
aagttcttctgaagcaacttaaagttatcaggcatgcatggatcttggaggaatcagatgtgcagtcagggaccatagcacaagacaggcgtcttctactg
gtgctaccagcaaatgctggaagccgggaacactgggtacgttggaaaccacgtgatgtgaagaagtaagataaactgtaggagaaaagcatttcgtag
tgggccatgaagcctttcaggacatgtattgcagtatgggccggcccattacgcaattggacgacaacaaagactagtattagtaccacctcggctatcca
catagatcaaagctgatttaaaagagttgtgcagatgatccgtggcaggagaccgaggtctcgGTTTTATGGCTGGAAATAGCAAG
TTAAAATAAGGctagtccgttatcaacttgaaaaagtggcACCGAGTCGGTGCTTTTTTTgcctgcaggtccacaaattcgg
gtcaaggcggaagccagcgcgccaccccacgtcagcaaatacggaggcgcggggttgacggcgtcacccggtcctaacggcgaccaacaaaccag
ccagaagaaattacagtaaaaaaaaagtaaattgcactttgatccaccttttattacctaagtctcaatttggatcacccttaaacctatcttttcaatttgggcc
gggttgtggtttggactaccatgaacaacttttcgtcatgtctaacttccctttcagcaaacatatgaaccatatatagaggagatcggccgtatactagagct
gatgtgtttaaggtcgttgattgcacgagaaaaaaaaatccaaatcgcaacaatagcaaatttatctggttcaaagtgaaaagatatgtttaaaggtagtcca
aagtaaaacttatagataataaaatgtggtccaaagcgtaattcactcaaaaaaaatcaacgagacgtgtaccaaacggagacaaacggcatcttctcgaa
atttcccaaccgctcgctcgcccgcctcgtcttcccggaaaccgcggtggtttcagcgtggcggattctccaagcagacggagacgtcacggcacggg
actcctcccaccacccaaccgccataaataccagcccctcatctcctctcctcgcatcagctccacccccgaaaaatttctccccaatctcgcgaggctct
cgtcgtcgaatcgaatcctctcgcgtcctcaaggtacgctgcttctcctctcctcgcttcgtttcgattcgatttcggacgggtgaggttgttttgttgctagatc
cgattggtggttagggttgtcgatgtgattatcgtgagatgtttaggggttgtagatctgatggttgtgatttgggcacggttggttcgataggtggaatcgtg
gttaggttttgggattggatgttggttctgatgattgggggaatttttacggttagatgaattgttggatgattcgattggggaaatcggtgtagatctgttggg
gaattgtggaactagtcatgcctgagtgattggtgcgatttgtagcgtgttccatcttgtaggccttgttgcgagcatgttcagatctactgttccgctcttgatt
gagttattggtgccatgggttggtgcaaacacaggctttaatatgttatatctgttttgtgtttgatgtagatctgtagggtagttcttcttagacatggttcaattat
gtagcttgtgcgtttcgatttgatttcatatgttcacagattagataatgatgaactcttttaattaattgtcaatggtaaataggaagtcttgtcgctatatctgtcat
aatgatctcatgttactatctgccagtaatttatgctaagaactatattagaatatcatgttacaatctgtagtaatatcatgttacaatctgtagttcatctatataat
ctattgtggtaatttctttttactatctgtgtgaagattattgccactagttcattctacttatttctgaagttcaggatacgtgtgctgttactacctatctgaatacat
gtgtgatgtgcctgttactatctttttgaatacatgtatgttctgttggaatatgtttgctgtttgatccgttgttgtgtccttaatcttgtgctagttcttaccctatctgt
ttggtgattatttcttgcagatagttatcaacaagtttgtacaaaaaagcaggcttcgaaggagatagaaccaattctctaaggaaatacttaaccatggacta
taaggaccacgacggagactacaaggatcatgatattgattacaaagacgatgacgataagatggccccaaagaagaagcggaaggtcggtatccacg
gagtcccagcagccAAAAAGCCTTACTCCATCGGTCTTGACATCGGTACTAACAGCGTGGGCTGGGC
CGTTATTACTGATGATTACAAGGTGCCTGCAAAGAAGATGAAGGTGCTCGGCAATACGGAC
AGATCACACATCAAGAAGAATCTCATTGGTGCGCTACTTTTTGACGCCGGGAACACTGCTGA
GGATAGGCGCCTGAAGAGAACGGCGCGGCGCCGCTATACAAGACGGAGGAATAGAATACT
GTATTTGCAGGAAATCTTCGCAGAAGAAATGAACAAGATTGATGAGTCCTTCTTCCACCGGC
TCGACGACAGCTTTCTCGTGCCCGAGGATAAAAGGGGCTCCAAGTATCCAATATTTGCTACT
CTGCAAGAAGAGAAGGAGTACCACAAGCAGTTCCCTACCATCTACCATCTCCGGAAACAAT
TGGCTGAATCTAATGAGAAGGCAGATCTGCGATTGGTATATCTTGCTCTGGCTCACATGATC
AAGTACAGAGGGCATTTTCTCATTGACGATCCCAAATTCAAAGTACAAAACAATGATATACA
GGGTCTGTTTGAGAAATTTGTTGAAGAATACGACAATGTGCAGGAAACATCTTTGTCTAAGA
TAAAACTCAATGTCACAGAAATACTTACCGCCAAGATTCCGAAAAGTGAGAAGCAAGAGCA
GCTTCTCAAGAACTACCCATCGGAAAAGAAGAACACGTTATTTGGGAACTTAATCGGCCTTG
CGTTGGGCCTCACACCAAATTTCAAGACTAATTTTAGCTTGGAAAATGACGCTAAACTGCAA
ATCTCAAGCGAAAGTTATGAGGAGGACCTCGGCTCCCTCCTTGCACTAATTGGAGAGAACTT
CATTGAGCTGTTCTCCGCGGTCAAGAACTTGAGTGATGGGATTCTCCTTGCTGGGATAGTTTC
AGATGAATCACCCCATGCTCCCCTCTCCACCAAAATGGTTATAAGGTTCAAGGAGCACGAGG
AAGACCTCGCCGCACTCAAGCATTTCATTAAGGCCAATTTGCCGGAGAAGTACGATGAGGTT
TTTTCAGATGACTCAAAGAACGGCTACGCCGGGTACGTCGGTGTCGATAGTAAGGTTCGCAA

AAGGAACGGGAAATTAGCAACCGAGGAGGAGTTCTACAAATACCTCAAGGATATTTTGAAC
AACGTGAAAGGCGCCGATTATTTCTTGGAGAAAATTAAACGGGAGGATTTACTCCGCAAAC
AGCGCACATTCGACAATGGGACTATCCCATATCAGGTTCATTTAGAAGAGATGAAGGCCATT
TTGCAAAACCAAGGCGAGTATTACCCCTTTCTTAAAGAGAACAAGGAGAAGATTCAACAGA
TCCTGACATTCAGAATCCCGTACTACGTAGGACCTCTGGCACGCGGCAACCGCGACTTCGCG
TGGTTGACGCGCAACTCAGATCAGGCGATTCGACCATGGAACTTTGAGGAGGTCGTGGATA
AGGCCTCCTCCGCGGAGGACTTCATCAACAAAATGACAAACTATGACCTCTATCTACCAGAG
GAAAAGGTCCTTCCCAAGCATTCGCTCTTGTACGAGACATTCGCTGTCTACAACGAGCTGAC
CAAAGTTAAGTTTATCGCTGAAGGACTGCGTGACTACCAATTCCTAGACTCCGGTCAAAAGA
AACAGATAGTTAATCAGCTGTTTAAAGAAAAGAGAAAAGTGACTGAGAAAGATATTATCCA
TTACCTCCACAACGTGGACGGTTATGATGGGATCGAATTAAAAGGAATCGAGAAGCAGTTT
AATGCTAGTCTGTCGACGTATCATGATCTCCTAAAAATCATTAAGGACAAGGAGTTTATGGA
CGATCCTAAGAACGAGGAGATCCTCGAGAACATCGTCCATACACTCACGATATTCGAGGAC
CGCGAGATGATTAAGCAGAGGTTGGCTCAGTACGACTCTCTCTTTGATGAGAAAGTCATCAA
AGCTCTAACCCGTCGCCACTATACTGGCTGGGGCAAGTTATCCGCAAAGCTGATAAATGGCA
TCTGTGATAAGCAAACAAATAAAACGATCCTGGACTTTCTTATTGACGACGACAAGATTAAC
AGAAACTTCATGCAGCTTATCAACGACGACGGCCTTTCTTTCAAGGATATAATTCAAAAGGC
GCAGGTGGTCGGCAAGATCGACGATGTGAAGCAAGTTGTGCAGGAGCTTCCAGGTTCTCCTG
CGATTAAAAAGGGTATTTTGCAGTCGATAAAGATTGTTGATGAGTTGGTCAAGGTCATGGGC
CACGCCCCTGAGTCCATTGTTATCGAGATGGCCAGGGAGAATCAGACTACCGCCAGGGGCA
AGAAAAACTCCCAGCAGAGATACAAGCGCATTGAGGACGCCTTAAAAAATCTAGCGCCGGG
GTTGGATTCTAATATCCTCAAGGAAAACCCAACAGATAATATCCAGTTACAAAACGACCGCC
TCTTCCTCTACTATCTTCAAAATGGTAAGGACATGTATACCGGTGAAGCCCTCGACATAAAT
CAACTCAGCAATTATGATATAGATCATATTGTGCCGCAGGCGTTTATAAAGGACGATAGTCT
TGATAACAGGGTGCTTACTAGCAGCAAGGATAACAGGGGAAAGTCTGACAATGTGCCTTCT
ATAGAAGTTGTTCAGAAGCGTAAGGCCTTCTGGCAGCAACTTCTGGACTCGAAATTGATAAG
CGAGCGAAAGTTCAATAACCTCACCAAGGCAGAGCGCGGTGGACTCGACGAACGTGACAAA
GTTGGGTTCATTAAGAGGCAGTTGGTTGAAACACGGCAGATTACAAAACATGTCGCTCAGAT
ACTCGATGCCCGGTTCAACACGGAGGTAAACGAGAAGAACCAGAAGATAAGAAAGGTAAA
AATTATTACTCTGAAATCCAACCTCGTTTCAAATTTCAGGAAAGAGTTCGGCCTGTACAAGG
TCCGTGAGATAAATGACTACCACCACGCCCACGATGCATACCTGAATGCAGTCGTGGCGAA
AGCGATCCTGAAGAAGTACCCTAAACTTGAACCGGAGTTCGTGTATGGAGACTACCAGAAG
TACGACCTTAAGAGGTACATCTCACGCTCCAAAGATCCAAAGGAAATAGAGAAAGCAACAG
AAAAGTACTTCTTTTATAGCAATTTGCTGAATTTTTTCAAGGAAGAAGTGCACTACGCTGAT
GGAACCATCATCAAGAGGGAGAATATAGAGTACTCTAAGGATACTGGCGAGATAGCATGGA
ATAAAGAAAAGGATTTCGCCACTGTACGCAAGGTACTGAGCTGCCCTCAGGTCAATATCGTG
AAAAAGACCGAGGTTCAAACTGGCGGCTTCTCCAAGGAGTCAATCCTCCCGAAGAGAAATT
CTGACAAGTTAATCGCCCGAAAAAAGGACTGGGACCCGAAGAAGTATGGAGGCTTCGATTC
ACCGACGGTGGCATATTCCGTGCTGGTGGTTGCCAAGGTGGAGAAGGGCAAGTCAAAAAAG
CTCAAGTCAGTAAAGGAACTTGTCGGAATCACGATCATGGAGCGGTCCTCGTTCGAGAAGG
ACCCGGTCGACTTCCTGGAAGCAAAAGGGTACAAGGAGGTGAGAAAAGATTTGATTATCAA
GCTCCCCAAGTATTCTCTTTTCGAGCTGGAAAATGGACGGAAGCGCATGCTCGCGTCGGCCG
GCGAGCTTCAGAAAGGAAATGAACTGGCCCTCCCTTCAAAATATGTCAACTTCTTATATCTG
GCAAGTCATTATGAAAAACTGAAAGGGAGTCCGGAGGATAATGAACAAAAACAACTGTTTG
TCGAGCAGCACAAACATTACCTCGATGAGATTATTGAGCAAATCTCCGAGTTCTCCAAGAGG
GTCATTCTCGCGGACGCAAATCTCGACAAGGTATTATCTGCCTACAATAAACATAGAGATAA
GCCAATTAGGGAACAGGCGGAGAACATAATCCACCTCTTCACCCTCACGAACCTCGGCGCCC
CAGCCGCGTTTAAGTACTTTGACACCACCATCGACAGGAAACGATACACTTCTACAAAAGAA
GTGCTCGATGCCACGCTGATCCATCAGAGCATTACAGGTCTTTATGAAACCCGTATAGATCT
CAGCCAACTCGGAGGGGACAAGCGGCCGGCGGCCACCAAGAAAGCTGGCCAAGCCAAGAA
GAAGAAATGAGCTCAGATAATctagacccagctttcttgtacaaagtggttgataacagcgactacaaggatgacgatgacaaggct

tagagctcgaatttccccgatcgttcaaacatttggcaataaagtttcttaagattgaatcctgttgccggtcttgcgatgattatcatataatttctgttgaattac
gttaagcatgtaataattaacatgtaatgcatgacgttatttatgagatgggtttttatgattagagtcccgcaattatacatttaatacgcgatagaaaacaaaa
tatagcgcgcaaactaggataaattatcgcgcgcggtgtcatctatgttactagatcgggaa

Seq ID No. 3

(CaMV Poly(A) Signal-HPT-CaMV35s Promoter (enhanced)-OsU3 Promoter-crRNA cloning site-ΔtracrRNA L-PolyT-OsUbi10 Promoter-SV40 NLS-PAiD-Nucleoplasmin NLS-NOS T)

ttcggggggatctggattttagtactggattttggttttaggaattagaaattttattgatagaagtattttacaaatacaaatacatactaagggtttcttatatgctc
aacacatgagcgaaaccctataggaaccctaattcccttatctgggaactactcacacattattatggagaaactcgagcttgtcgatcgacagatcccggt
cggcatctactctatttctttgccctcggacgagtgctggggcgtcggtttccactatcggcgagtacttctacacagccatcggtccagacggccgcgctt
ctgcgggcgatttgtgtacgcccgacagtcccggctccggatcggacgattgcgtcgcatcgaccctgcgcccaagctgcatcatcgaaattgccgtca
accaagctctgatagagttggtcaagaccaatgcggagcatatacgcccggagtcgtggcgatcctgcaagctccggatgcctccgctcgaagtagcg
cgtctgctgctccatacaagccaaccacggcctccagaagaagatgttggcgacctcgtattgggaatccccgaacatcgcctcgctccagtcaatgacc
gctgttatgcggccattgtccgtcaggacattgttggagccgaaatccgcgtgcacgaggtgccggacttcggggcagtcctcggcccaaagcatcagc
tcatcgagagcctgcgcgacggacgcactgacggtgtcgtccatcacagtttgccagtgatacacatggggatcagcaatcgcgcatatgaaatcacgc
catgtagtgtattgaccgattccttgcggtccgaatgggccgaacccgctcgtctggctaagatcggccgcagcgatcgcatccatagcctccgcgaccg
gttgtagaacagcgggcagttcggtttcaggcaggtcttgcaacgtgacaccctgtgaacggcgggagatgcaataggtcaggctctcgctaaactccc
caatgtcaagcacttccggaatcgggagcgcggccgatgcaaagtgccgataaacataacgatctttgtagaaaccatcggcgcagctatttacccgca
ggacatatccacgccctcctacatcgaagctgaaagcacgagattcttcgccctccgagagctgcatcaggtcggagacgctgtcgaacttttcgatcag
aaacttctcgacagacgtcgcggtgagttcaggctttttcatatctcattgccccccggatctgcgaaagctcgagagagatagatttgtagagagagactg
gtgatttcagcgtgtcctctccaaatgaaatgaacttccttatatagaggaagggtcttgcgaaggatagtgggattgtgcgtcatcccttacgtcagtggag
atatcacatcaatccacttgctttgaagacgtggttggaacgtcttctttttccacgatgctcctcgtgggtgggggtccatctttgggaccactgtcggcaga
ggcatcttgaacgatagcctttcctttatcgcaatgatggcatttgtaggtgccaccttccttttctactgtccttttgatgaagtgacagatagctgggcaatgg
aatccgaggaggtttcccgatattaccctttgttgaaaagtctcaatagccctttggtcttctgagactgtatctttgatattcttggagtagacgagagtgtcgt
gctccaccatgttcacatcaatccacttgctttgaagacgtggttggaacgtcttctttttccacgatgctcctcgtgggtgggggtccatctttgggaccact
gtcggcagaggcatcttgaacgatagcctttcctttatcgcaatgatggcatttgtaggtgccaccttccttttctactgtccttttgatgaagtgacagatagct
gggcaatggaatccgaggaggtttcccgatattaccctttgttgaaaagtctcaatagccctttggtcttctgagactgtatctttgatattcttggagtagacg
agagtgtcgtgctccaccatgttggcaagctgctctagccaatacgcaaaccgcctctccccgcgcgttggccgattcattaatgcagctggcacgacag
gtttcccgactggaaagcgggcagtgagcgcaacgcaattaatgtgagttagctcactcattaggcaccccaggctttacactttatgcttccggctcgtat
gttgtgtggaattgtgagcggataacaatttcacacaggaaacagctatgacatgattacgccaagcttaaggaatctttaaacatacgaacagatcactta
aagttcttctgaagcaacttaaagttatcaggcatgcatggatcttggaggaatcagatgtgcagtcagggaccatagcacaagacaggcgtcttctactg
gtgctaccagcaaatgctggaagccgggaacactgggtacgttggaaaccacgtgatgtgaagaagtaagataaactgtaggagaaaagcatttcgtag
tgggccatgaagcctttcaggacatgtattgcagtatgggccggcccattacgcaattggacgacaacaaagactagtattagtaccacctcggctatcca
catagatcaaagctgatttaaaagagttgtgcagatgatccgtggcaggagaccgaggtctcgGTTTTATGGCTGATAAATTTCTTT
GAATTTCTCCTTGATTATTTGTTATAAAAGTTATAAAATAATCTTGTTGGAACCATTCAAAAC
AGCATAGCAAGTTAAAATAAGGCTAGTCCgttatcaacttgaaaaagtggcACCGAGTCGGTGCTTTTTTT
gcctgcaggtccacaaattcgggtcaaggcggaagccagcgcgccaccccacgtcagcaaatacggaggcgcggggttgacggcgtcacccggtc
ctaacggcgaccaacaaaccagccagaagaaattacagtaaaaaaaaagtaaattgcactttgatccaccttttattacctaagtctcaatttggatcaccct
taaacctatcttttcaatttgggccgggttgtggtttggactaccatgaacaacttttcgtcatgtctaacttccctttcagcaaacatatgaaccatatatagag
gagatcggccgtatactagagctgatgtgtttaaggtcgttgattgcacgagaaaaaaaaatccaaatcgcaacaatagcaaatttatctggttcaaagtga
aaagatatgtttaaaggtagtccaaagtaaaacttatagataataaaatgtggtccaaagcgtaattcactcaaaaaaaatcaacgagacgtgtaccaaacg
gagacaaacggcatcttctcgaaatttcccaaccgctcgctcgcccgcctcgtcttcccggaaaccgcggtggtttcagcgtggcggattctccaagcag
acggagacgtcacggcacgggactcctcccaccacccaaccgccataaataccagcccccctcatctcctctcctcgcatcagctccacccccgaaaaat
ttctccccaatctcgcgaggctctcgtcgtcgaatcgaatcctctcgcgtcctcaaggtacgctgcttctcctctcctcgcttcgtttcgattcgatttcggacg
ggtgaggttgttttgttgctagatccgattggtggttagggttgtcgatgtgattatcgtgagatgtttaggggttgtagatctgatggttgtgatttggcacgg
ttggttcgataggtggaatcgtggttaggttttgggattggatgttggttctgatgattgggggggaatttttacggttagatgaattgttggatgattcgattggg

gaaatcggtgtagatctgttggggaattgtggaactagtcatgcctgagtgattggtgcgatttgtagcgtgttccatcttgtaggccttgttgcgagcatgttc
agatctactgttccgctcttgattgagttattggtgccatgggttggtgcaaacacaggctttaatatgttatatctgttttgtgtttgatgtagatctgtagggtag
ttcttcttagacatggttcaattatgtagcttgtgcgtttcgatttgatttcatatgttcacagattagataatgatgaactcttttaattaattgtcaatggtaaatag
gaagtcttgtcgctatatctgtcataatgatctcatgttactatctgccagtaatttatgctaagaactatattagaatatcatgttacaatctgtagtaatatcatgt
tacaatctgtagttcatctatataatctattgtggtaatttctttttactatctgtgtgaagattattgccactagttcattctacttatttctgaagttcaggatacgtgt
gctgttactacctatctgaatacatgtgtgatgtgcctgttactatctttttgaatacatgtatgttctgttggaatatgtttgctgtttgatccgttgttgtgtccttaat
cttgtgctagttcttaccctatctgtttggtgattatttcttgcagatagttatcaacaagtttgtacaaaaaagcaggcttcgaaggagatagaaccaattctcta
aggaaatacttaaccatggactataaggaccacgacggagactacaaggatcatgatattgattacaaagacgatgacgataagatggcccccaaagaag
aagcggaaggtcggtatccacggagtcccagcagccAAAAAGCCTTACTCCATCGGTCTTGACATCGGTACTAAC
AGCGTGGGCTGGGCCGTTATTACTGATGATTACAAGGTGCCTGCAAAGAAGATGAAGGTGC
TCGGCAATACGGACAGATCACACATCAAGAAGAATCTCATTGGTGCGCTACTTTTTGACGCC
GGGAACACTGCTGAGGATAGGCGCCTGAAGAGAACGGCGCGGCGCCGCTATACAAGACGG
AGGAATAGAATACTGTATTTGCAGGAAATCTTCGCAGAAGAAATGAACAAGATTGATGAGT
CCTTCTTCCACCGGCTCGACGACAGCTTTCTCGTGCCCGAGGATAAAGGGGCTCCAAGTAT
CCAATATTTGCTACTCTGCAAGAAGAGAAGGAGTACCACAAGCAGTTCCCTACCATCTACCA
TCTCCGGAAACAATTGGCTGAATCTAATGAGAAGGCAGATCTGCGATTGGTATATCTTGCTC
TGGCTCACATGATCAAGTACAGAGGGCATTTTCTCATTGACGATCCCAAATTCAAGTACAA
AACAATGATATACAGGGTCTGTTTGAGAAATTTGTTGAAGAATACGACAATGTGCAGGAAA
CATCTTTGTCTAAGATAAAACTCAATGTCACAGAAATACTTACCGCCAAGATTCCGAAAAGT
GAGAAGCAAGAGCAGCTTCTCAAGAACTACCCATCGGAAAAGAAGAACACGTTATTTGGGA
ACTTAATCGGCCTTGCGTTGGGCCTCACACCAAATTTCAAGACTAATTTTAGCTTGGAAAAT
GACGCTAAACTGCAAATCTCAAGCGAAAGTTATGAGGAGGACCTCGGCTCCCTCCTTGCACT
AATTGGAGAGAACTTCATTGAGCTGTTCTCCGCGGTCAAGAACTTGAGTGATGGGATTCTCC
TTGCTGGGATAGTTTCAGATGAATCACCCCATGCTCCCCTCTCCACCAAAATGGTTATAAGG
TTCAAGGAGCACGAGGAAGACCTCGCCGCACTCAAGCATTTCATTAAGGCCAATTTGCCGGA
GAAGTACGATGAGGTTTTTCAGATGACTCAAAGAACGGCTACGCCGGGTACGTCGGTGTCG
ATAGTAAGGTTCGCAAAAGGAACGGGAAATTAGCAACCGAGGAGGAGTTCTACAAATACCT
CAAGGATATTTTGAACAACGTGAAAGGCGCCGATTATTTCTTGGAGAAAATTAAACGGGAG
GATTTACTCCGCAAACAGCGCACATTCGACAATGGGACTATCCCATATCAGGTTCATTTAGA
AGAGATGAAGGCCATTTTGCAAAACCAAGGCGAGTATTACCCCTTTCTTAAAGAGAACAAG
GAGAAGATTCAACAGATCCTGACATTCAGAATCCCGTACTACGTAGGACCTCTGGCACGCGG
CAACCGCGACTTCGCGTGGTTGACGCGCAACTCAGATCAGGCGATTCGACCATGGAACTTTG
AGGAGGTCGTGGATAAGGCCTCCTCCGCGGAGGACTTCATCAACAAAATGACAAACTATGA
CCTCTATCTACCAGAGGAAAAGGTCCTTCCCAAGCATTCGCTCTTGTACGAGACATTCGCTG
TCTACAACGAGCTGACCAAAGTTAAGTTTATCGCTGAAGGACTGCGTGACTACCAATTCCTA
GACTCCGGTCAAAAGAAACAGATAGTTAATCAGCTGTTTAAAGAAAAGAGAAAAGTGACTG
AGAAAGATATTATCCATTACCTCCACAACGTGGACGGTTATGATGGGATCGAATTAAAAGG
AATCGAGAAGCAGTTTAATGCTAGTCTGTCGACGTATCATGATCTCCTAAAAATCATTAAGG
ACAAGGAGTTTATGGACGATCCTAAGAACGAGGAGATCCTCGAGAACATCGTCCATACACT
CACGATATTCGAGGACCGCGAGATGATTAAGCAGAGGTTGGCTCAGTACGACTCTCTCTTTG
ATGAGAAAGTCATCAAAGCTCTAACCCGTCGCCACTATACTGGCTGGGGCAAGTTATCCGCA
AAGCTGATAAATGGCATCTGTGATAAGCAAACAATAAAACGATCCTGGACTTTCTTATTGA
CGACGACAAGATTAACAGAAACTTCATGCAGCTTATCAACGACGACGGCCTTTCTTTCAAGG
ATATAATTCAAAAGGCGCAGGTGGTCGGCAAGATCGACGATGTGAAGCAAGTTGTGCAGGA
GCTTCCAGGTTCTCCTGCGATTAAAAAGGGTATTTTGCAGTCGATAAAGATTGTTGATGAGT
TGGTCAAGGTCATGGGCCACGCCCCTGAGTCCATTGTTATCGAGATGGCCAGGGAGAATCAG
ACTACCGCCAGGGGCAAGAAAAACTCCCAGCAGAGATACAAGCGCATTGAGGACGCCTTAA
AAAATCTAGCGCCGGGGTTGGATTCTAATATCCTCAAGGAAAACCCAACAGATAATATCCA
GTTACAAAACGACCGCCTCTTCCTCTACTATCTTCAAAATGGTAAGGACATGTATACCGGTG
AAGCCCTCGACATAAATCAACTCAGCAATTATGATATAGATCATATTGTGCCGCAGGCGTTT
ATAAAGGACGATAGTCTTGATAACAGGGTGCTTACTAGCAGCAAGGATAACAGGGGAAAGT

CTGACAATGTGCCTTCTATAGAAGTTGTTCAGAAGCGTAAGGCCTTCTGGCAGCAACTTCTG
GACTCGAAATTGATAAGCGAGCGAAAGTTCAATAACCTCACCAAGGCAGAGCGCGGTGGAC
TCGACGAACGTGACAAAGTTGGGTTCATTAAGAGGCAGTTGGTTGAAACACGGCAGATTAC
AAAACATGTCGCTCAGATACTCGATGCCCGGTTCAACACGGAGGTAAACGAGAAGAACCAG
AAGATAAGAAAGGTAAAAATTATTACTCTGAAATCCAACCTCGTTTCAAATTTCAGGAAAGA
GTTCGGCCTGTACAAGGTCCGTGAGATAAATGACTACCACCACGCCCACGATGCATACCTGA
ATGCAGTCGTGGCGAAAGCGATCCTGAAGAAGTACCCTAAACTTGAACCGGAGTTCGTGTAT
GGAGACTACCAGAAGTACGACCTTAAGAGGTACATCTCACGCTCCAAAGATCCAAAGGAAA
TAGAGAAAGCAACAGAAAAGTACTTCTTTTATAGCAATTTGCTGAATTTTTTCAAGGAAGAA
GTGCACTACGCTGATGGAACCATCATCAAGAGGGAGAATATAGAGTACTCTAAGGATACTG
GCGAGATAGCATGGAATAAAGAAAAGGATTTCGCCACTGTACGCAAGGTACTGAGCTGCCC
TCAGGTCAATATCGTGAAAAAGACCGAGGTTCAAACTGGCGGCTTCTCCAAGGAGTCAATCC
TCCCGAAGAGAAATTCTGACAAGTTAATCGCCCGAAAAAAGGACTGGGACCCGAAGAAGTA
TGGAGGCTTCGATTCACCGACGGTGGCATATTCCGTGCTGGTGGTTGCCAAGGTGGAGAAGG
GCAAGTCAAAAAAGCTCAAGTCAGTAAAGGAACTTGTCGGAATCACGATCATGGAGCGGTC
CTCGTTCGAGAAGGACCCGGTCGACTTCCTGGAAGCAAAAGGGTACAAGGAGGTGAGAAAA
GATTTGATTATCAAGCTCCCCAAGTATTCTCTTTTCGAGCTGGAAAATGGACGGAAGCGCAT
GCTCGCGTCGGCCGGCGAGCTTCAGAAAGGAAATGAACTGGCCCTCCCTTCAAAATATGTCA
ACTTCTTATATCTGGCAAGTCATTATGAAAAACTGAAAGGGAGTCCGGAGGATAATGAACA
AAAACAACTGTTTGTCGAGCAGCACAAACATTACCTCGATGAGATTATTGAGCAAATCTCCG
AGTTCTCCAAGAGGGTCATTCTCGCGGACGCAAATCTCGACAAGGTATTATCTGCCTACAAT
AAACATAGAGATAAGCCAATTAGGGAACAGGCGGAGAACATAATCCACCTCTTCACCCTCA
CGAACCTCGGCGCCCCAGCCGCGTTTAAGTACTTTGACACCACCATCGACAGGAAACGATAC
ACTTCTACAAAAGAAGTGCTCGATGCCACGCTGATCCATCAGAGCATTACAGGTCTTTATGA
AACCCGTATAGATCTCAGCCAACTCGGAGGGGACAAGCGGCCGGCGGCCACCAAGAAAGCT
GGCCAAGCCAAGAAGAAGAAATGAGCTCAGATAATctagacccagctttcttgtacaaagtggttgataacagcgacta
caaggatgacgatgacaaggcttagagctcgaatttccccgatcgttcaaacatttggcaataaagtttcttaagattgaatcctgttgccggtcttgcgatga
ttatcatataatttctgttgaattacgttaagcatgtaataattaacatgtaatgcatgacgttatttatgagatgggtttttatgattagagtcccgcaattatacatt
taatacgcgatagaaaacaaaatatagcgcgcaaactaggataaattatcgcgcgcggtgtcatctatgttactagatcgggaa

Seq ID No. 4

(OsU3 Promoter-crRNA cloning site-tracrRNA-PolyT-ZmUbi Promoter-SV40 NLS (Bipartite)-Adenine Deaminase-Linker-nPAiD-Nucleoplsmin NLS-E9 terminator-CaMV35S Promoter (enhanced)-HPT-CaMV Poly(A) Signal)

aacgacggccagtgccaagcttaaggaatctttaaacatacgaacagatcacttaaagttcttctgaagcaacttaaagttatcaggcatgcatggatcttg
gaggaatcagatgtgcagtcagggaccatagcacaagacaggcgtcttctactggtgctaccagcaaatgctggaagccgggaacactgggtacgttg
gaaaccacgtgatgtgaagaagtaagataaactgtaggagaaaagcatttcgtagtgggccatgaagcctttcaggacatgtattgcagtatgggccggc
ccattacgcaattggacgacaacaaagactagtattagtaccacctcggctatccacatagatcaaagctgatttaaaagagttgtgcagatgatccgtggc
aggagaccgaggtctcggtttaagagctatgctggaaacagcatagcaagtttaaataaggctagtccgttatcaacttgaaaaagtggcaccgagtcgg
tgctttttgttttagagctagaaatagcaagttaaaataaggctagtgctagtgcagcgtgacccggtcgtgcccctcactagagataatgagcattgcatgt
ctaagttataaaaaattaccacatatttttttgtcacacttgtttgaagtgcagtttatctatctttatacatatatttaaactttactctacgaataatataatctatagt
actacaataatatcagtgttttagagaatcatataaatgaacagttagacatggtctaaaggacaattgagtattttgacaacaggactctacagttttatcttttt
agtgtgcatgtgttctcctttttttttgcaaatagcttcacctatataatacttcatccattttattagtacatccatttagggtttagggttaatggtttttatagactaa
tttttttagtacatctattttattctattttagcctctaaattaagaaaactaaaactctattttagtttttttatttaataatttagatataaaatagaataaaataaagtga
ctaaaaattaaacaaataccctttaagaaattaaaaaaactaaggaaacatttttcttgtttcgagtagataatgccagcctgttaaacgccgtcgacgagtct
aacggacaccaaccagcgaaccagcagcgtcgcgtcgggccaagcgaagcagacggcacggcatctctgtcgctgcctctggacccctctcgagag
ttccgctccaccgttggacttgctccgctgtcggcatccagaaattgcgtggcggagcggcagacgtgagccggcacggcaggcggcctcctcctcctc
tcacggcaccggcagctacgggggattcctttcccaccgctccttcgctttcccttcctcgcccgccgtaataaatagacacccctccacaccctctttcc

ccaacctcgtgttgttcggagcgcacacacacacaaccagatctcccccaaatccacccgtcggcacctccgcttcaaggtacgccgctcgtcctccccc
ccccccccctctctaccttctctagatcggcgttccggtccatgcttagggcccggtagttctacttctgttcatgtttgtgttagatccgtgtttgtgttagatccg
tgctgctagcgttcgtacacggatgcgacctgtacgtcagacacgttctgattgctaacttgccagtgtttctctttggggaatcctgggatggctctagccgt
tccgcagacgggatcgatttcatgattttttttgtttcgttgcatagggtttggtttgcccttttcctttatttcaatatatgccgtgcacttgtttgtcgggtcatctttt
catgcttttttttgtcttggttgtgatgatgtggtctggttgggcggtcgttctagatcggagtacaattctgtttcaaactacctggtggatttattaattttggatct
gtatgtgtgtgccatacatattcatagttacgaattgaagatgatggatggaaatatcgatctaggataggtatacatgttgatgcgggttttactgatgcatat
acagagatgcttttttgttcgcttggttgtgatgatgtggtgtggttgggcggtcgttcattcgttctagatcggagtagaatactgtttcaaactacctggtgtatt
tattaattttggaactgtatgtgtgtgtcatacatcttcatagttacgagtttaagatggatggaaatatcgatctaggataggtatacatgttgatgtgggttttac
tgatgcatatacatgatggcatatgcagcatctattcatatgctctaaccttgagtacctatctattataataaacaagtatgttttataattattttgatcttgatata
cttggatgatggcatatgcagcagctatatgtggatttttttagccctgccttcatacgctatttatttgcttggtactgtttcttttgtcgatgctcaccctgttgttt
ggtgttacttctgcaggtacctaggaagccaccatggattataaagatcacgacggcgactacaaagatcatgacatcgattacaaggacgacgacgac
aagatgaaacgtacagctgatggcagcgaatttgaatcgccgaagaagaagcggaaggtgagtgaggtggagttctcgcacgagtactggatgaggc
acgccctgacactcgcaaagcgagcacgcgatgaaagggaggtcccggtgggtgccgtcctcgttcttaataatcgggtaataggggaaggctggaat
agggccatcggactccacgatcctacagctcatgctgagatcatggcgctgcgccagggcggcctggtcatgcagaattataggctaattgacgcgacg
ctatacgtcacgttcgagccgtgcgttatgtgcgcaggcgccatgatccacagcaggattgggagagtcgtctttggcgtgaggaactccaaacggggg
gcggccggctccctcatgaacgttcttaattaccctggcatgaatcatcgtgtggagataacggaggggattcttgccgatgagtgcgccgccctgctgtg
tgatttctaccgtatgcctagacaggtcttcaacgcgcaaaaaaaagcacagtcctccatcaactccggcggcagtagcggtgggagcagtggatctga
gactcctggcacctccgagagcgctacaccagaatcatcgggggggtcaagcgggggtctatgaaaaagccttactccatcggtcttgccatcggtac
taacagcgtgggctgggccgttattactgatgattacaaggtgcctgcaaagaagatgaaggtgctcggcaatacggacagatcacacatcaagaagaa
tctcattggtgcgctactttttgacgccgggaacactgctgaggataggcgcctgaagagaacggcgcggcgccgctatacaagacggaggaatagaa
tactgtatttgcaggaaatcttcgcagaagaaatgaacaagattgatgagtccttcttccaccggctcgacgacagctttctcgtgcccgaggataaaaggg
gctccaagtatccaatatttgctactctgcaagaagagaaggagtaccacaagcagttccctaccatctaccatctccggaaacAATTGGCTGA
ATCTAATGAGAAGGCAGATCTGCGATTGGTATATCTTGCTCTGGCTCACATGATCAAGTACA
GAGGGCATTTTCTCATTGACGATCCCAAATTCAAAGTACAAAACAATGATATACAGGGTCTG
TTTGAGAAATTTGTTGAAGAATACGACAATGTGCAGGAAACATCTTTGTCTAAGATAAAACT
CAATGTCACAGAAATACTTACCGCCAAGATTCCGAAAAGTGAGAAGCAAGAGCAGCTTCTC
AAGAACTACCCATCGGAAAAGAAGAACACGTTATTTGGGAACTTAATCGGCCTTGCGTTGG
GCCTCACACCAAATTTCAAGACTAATTTTAGCTTGGAAAATGACGCTAAACTGCAAATCTCA
AGCGAAAGTTATGAGGAGGACCTCGGCTCCCTCCTTGCACTAATTGGAGAGAACTTCATTGA
GCTGTTCTCCGCGGTCAAGAACTTGAGTGATGGGATTCTCCTTGCTGGGATAGTTTCAGATG
AATCACCCCATGCTCCCCTCTCCACCAAAATGGTTATAAGGTTCAAGGAGCACGAGGAAGAC
CTCGCCGCACTCAAGCATTTCATTAAGGCCAATTTGCCGGAGAAGTACGATGAGGTTTTTTC
AGATGACTCAAAGAACGGCTACGCCGGGTACGTCGGTGTCGATAGTAAGGTTCGCAAAAGG
AACGGGAAATTAGCAACCGAGGAGGAGTTCTACAAATACCTCAAGGATATTTTGAACAACG
TGAAAGGCGCCGATTATTTCTTGGAGAAAATTAAACGGGAGGATTTACTCCGCAAACAGCG
CACATTCGACAATGGGACTATCCCATATCAGGTTCATTTAGAAGAGATGAAGGCCATTTTGC
AAAACCAAGGCGAGTATTACCCCTTTCTTAAAGAGAACAAGGAGAAGATTCAACAGATCCT
GACATTCAGAATCCCGTACTACGTAGGACCTCTGGCACGCGGCAACCGCGACTTCGCGTGGT
TGACGCGCAACTCAGATCAGGCGATTCGACCATGGAACTTTGAGGAGGTCGTGGATAAGGC
CTCCTCCGCGGAGGACTTCATCAACAAAATGACAAACTATGACCTCTATCTACCAGAGGAAA
AGGTCCTTCCCAAGCATTCGCTCTTGTACGAGACATTCGCTGTCTACAACGAGCTGACCAAA
GTTAAGTTTATCGCTGAAGGACTGCGTGACTACCAATTCCTAGACTCCGGTCAAAAGAAACA
GATAGTTAATCAGCTGTTTAAAGAAAAGAGAAAAGTGACTGAGAAAGATATTATCCATTAC
CTCCACAACGTGGACGGTTATGATGGGATCGAATTAAAAGGAATCGAGAAGCAGTTTAATG
CTAGTCTGTCGACGTATCATGATCTCCTAAAAATCATTAAGGACAAGGAGTTTATGGACGAT
CCTAAGAACGAGGAGATCCTCGAGAACATCGTCCATACACTCACGATATTCGAGGACCGCG
AGATGATTAAGCAGAGGTTGGCTCAGTACGACTCTCTCTTTGATGAGAAAGTCATCAAAGCT
CTAACCCGTCGCCACTATACTGGCTGGGGCAAGTTATCCGCAAAGCTGATAAATGGCATCTG
TGATAAGCAAACAAATAAAACGATCCTGGACTTTCTTATTGACGACGACAAGATTAACAGA
AACTTCATGCAGCTTATCAACGACGACGGCCTTTCTTTCAAGGATATAATTCAAAAGGCGCA
GGTGGTCGGCAAGATCGACGATGTGAAGCAAGTTGTGCAGGAGCTTCCAGGTTCTCCTGCGA

TTAAAAAGGGTATTTTGCAGTCGATAAAGATTGTTGATGAGTTGGTCAAGGTCATGGGCCAC
GCCCCTGAGTCCATTGTTATCGAGATGGCCAGGGAGAATCAGACTACCGCCAGGGGCAAGA
AAAACTCCCAGCAGAGATACAAGCGCATTGAGGACGCCTTAAAAAATCTAGCGCCGGGGTT
GGATTCTAATATCCTCAAGGAAAACCCAACAGATAATATCCAGTTACAAAACGACCGCCTCT
TCCTCTACTATCTTCAAAATGGTAAGGACATGTATACCGGTGAAGCCCTCGACATAAATCAA
CTCAGCAATTATGATATAGATCATATTGTGCCGCAGGCGTTTATAAAGGACGATAGTCTTGA
TAACAGGGTGCTTACTAGCAGCAAGGATAACAGGGGAAAGTCTGACAATGTGCCTTCTATA
GAAGTTGTTCAGAAGCGTAAGGCCTTCTGGCAGCAACTTCTGGACTCGAAATTGATAAGCGA
GCGAAAGTTCAATAACCTCACCAAGGCAGAGCGCGGTGGACTCGACGAACGTGACAAAGTT
GGGTTCATTAAGAGGCAGTTGGTTGAAACACGGCAGATTACAAAACATGTCGCTCAGATACT
CGATGCCCGGTTCAACACGGAGGTAAACGAGAAGAACCAGAAGATAAGAAAGGTAAAAAT
TATTACTCTGAAATCCAACCTCGTTTCAAATTTCAGGAAAGAGTTCGGCCTGTACAAGGTCC
GTGAGATAAATGACTACCACCACGCCCACGATGCATACCTGAATGCAGTCGTGGCGAAAGC
GATCCTGAAGAAGTACCCTAAACTTGAACCGGAGTTCGTGTATGGAGACTACCAGAAGTAC
GACCTTAAGAGGTACATCTCACGCTCCAAAGATCCAAAGGAAATAGAGAAAGCAACAGAAA
AGTACTTCTTTTATAGCAATTTGCTGAATTTTTTCAAGGAAGAAGTGCACTACGCTGATGGA
ACCATCATCAAGAGGGAGAATATAGAGTACTCTAAGGATACTGGCGAGATAGCATGGAATA
AAGAAAAGGATTTCGCCACTGTACGCAAGGTACTGAGCTGCCCTCAGGTCAATATCGTGAA
AAAGACCGAGGTTCAAACTGGCGGCTTCTCCAAGGAGTCAATCCTCCCGAAGAGAAATTCT
GACAAGTTAATCGCCCGAAAAAAGGACTGGGACCCGAAGAAGTATGGAGGCTTCGATTCAC
CGACGGTGGCATATTCCGTGCTGGTGGTTGCCAAGGTGGAGAAGGGCAAGTCAAAAAAGCT
CAAGTCAGTAAAGGAACTTGTCGGAATCACGATCATGGAGCGGTCCTCGTTCGAGAAGGAC
CCGGTCGACTTCCTGGAAGCAAAAGGGTACAAGGAGGTGAGAAAAGATTTGATTATCAAGC
TCCCCAAGTATTCTCTTTTCGAGCTGGAAAATGGACGGAAGCGCATGCTCGCGTCGGCCGGC
GAGCTTCAGAAAGGAAATGAACTGGCCCTCCCTTCAAAATATGTCAACTTCTTATATCTGGC
AAGTCATTATGAAAAACTGAAAGGGAGTCCGGAGGATAATGAACAAAAACAACTGTTTGTC
GAGCAGCACAAACATTACCTCGATGAGATTATTGAGCAAATCTCCGAGTTCTCCAAGAGGGT
CATTCTCGCGGACGCAAATCTCGACAAGGTATTATCTGCCTACAATAAACATAGAGATAAGC
CAATTAGGGAACAGGCGGAGAACATAATCCACCTCTTCACCCTCACGAACCTCGGCGCCCCA
GCCGCGTTTAAGTACTTTGACACCACCATCGACAGGAAACGATACACTTCTACAAAAGAAGT
GCTCGATGCCACGCTGATCCATCAGAGCATTACAGGTCTTTATGAAACCCGTATAGATCTCA
GCCAACTCGGAGGGGACAAGCGGCCGGCGGCCACCAAGAAAGCTGGCCAAGCCAAGAAGA
AGAAATGAGCTcagagctttcgttcgtatcatcggtttcgacaacgttcgtcaagttcaatgcatcagtttcattgcgcacacaccagaatcctact
gagtttgagtattatggcattgggaaaactgtttttcttgtaccatttgttgtgcttgtaatttactgtgttttttattcggttttcgctatcgaactgtgaaatggaaat
ggatggagaagagttaatgaatgatatggtccttttgttcattctcaaattaatattatttgttttttctcttatttgttgtgtgttgaatttgaaattataagagatatgc
aaacattttgttttgagtaaaaatgtgtcaaatcgtggcctctaatgaccgaagttaatatgaggagtaaaacacttgtagttgtaccattatgcttattcactag
gcaacaaatatattttcagacctagaaaagctgcaaatgttactgaatacaagtatgtcctcttgtgttttagacatttatgaactttcctttatgtaattttccaga
atccttgtcagattctaatcattgctttataattatagttatactcatggatttgtagttgagtatgaaaatatttttaatgcattttatgacttgccaattgattgacaa
cgaattcgtaatcatgtcatagctgtttcctgtgtgaaattgttatccgctcacaattccacacaacatacgagccggaagcataaagtgtaaagcctggggt
gcctaatgagtgagctaactcacattaattgcgttgcgctcactgcccgctttccagtcgggaaacctgtcgtgccagctgcattaatgaatcggccaacgc
gcggggagaggcggtttgcgtattggctagagcagcttgccaacatggtggagcacgacactctcgtctactccaagaatatcaaagatacagtctcaga
agaccaaagggctattgagacttttcaacaaagggtaatatcgggaaacctcctcggattccattgcccagctatctgtcacttcatcaaaaggacagtaga
aaaggaaggtggcacctacaaatgccatcattgcgataaaggaaaggctatcgttcaagatgcctctgccgacagtggtcccaaagatggacccccacc
cacgaggagcatcgtggaaaaagaagacgttccaaccacgtcttcaaagcaagtggattgatgtgataacatggtggagcacgacactctcgtctactcc
aagaatatcaaagatacagtctcagaagaccaaagggctattgagacttttcaacaaagggtaatatcgggaaacctcctcggattccattgcccagctat
ctgtcacttcatcaaaaggacagtagaaaaggaaggtggcacctacaaatgccatcattgcgataaaggaaaggctatcgttcaagatgcctctgccgac
agtggtcccaaagatggacccccacccacgaggagcatcgtggaaaaagaagacgttccaaccacgtcttcaaagcaagtggattgatgtgatatctcc
actgacgtaagggatgacgcacaatcccactatccttcgcaagaccttcctctatataaggaagttcatttcatttggagaggacacgctgaaatcaccagt
ctctctctacaaatctatctctctcgagctttcgcagatcccggggggcaatgagatatgaaaaagcctgaactcaccgcgacgtctgtcgagaagtttctg
atcgaaaagttcgacagcgtctccgacctgatgcagctctcggagggcgaagaatctcgtgctttcagcttcgatgtaggagggcgtggatatgtcctgc
gggtaaatagctgcgccgatggtttctacaaagatcgttatgtttatcggcactttgcatcggccgcgctcccgattccggaagtgcttgacattggggagtt

tagcgagagcctgacctattgcatctcccgccgtgcacagggtgtcacgttgcaagacctgcctgaaaccgaactgcccgctgttctacaaccggtcgcg
gaggctatggatgcgatcgctgcggccgatcttagccagacgagcgggttcggcccattcggaccgcaaggaatcggtcaatacactacatggcgtga
tttcatatgcgcgattgctgatccccatgtgtatcactggcaaactgtgatggacgacaccgtcagtgcgtccgtcgcgcaggctctcgatgagctgatgct
ttgggccgaggactgccccgaagtccggcacctcgtgcacgcggatttcggctccaacaatgtcctgacggacaatggccgcataacagcggtcattg
actggagcgaggcgatgttcggggattcccaatacgaggtcgccaacatcttcttctggaggccgtggttggcttgtatggagcagcagacgcgctactt
cgagcggaggcatccggagcttgcaggatcgccacgactccgggcgtatatgctccgcattggtcttgaccaactctatcagagcttggttgacggcaat
ttcgatgatgcagcttgggcgcagggtcgatgcgacgcaatcgtccgatccggagccgggactgtcgggcgtacacaaatcgcccgcagaagcgcgg
ccgtctggaccgatggctgtgtagaagtactcgccgatagtggaaaccgacgccccagcactcgtccgagggcaaagaaatagagtagatgccgacc
ggatctgtcgatcgacaagctcgagtttctccataataatgtgtgagtagttcccagataagggaattagggttcctatagggtttcgctcatgtgttgagcat
ataagaaacccttagtatgtatttgtatttgtaaaatacttctatcaataaaatttctaattcctaaaaccaaaatccagtactaaaatccagatccccgaat

Seq ID No. 5

(OsU3 Promoter-crRNA cloning site-tracrRNA-PolyT-ZmUbi Promoter-Cytidine deaminase-SV40 NLS-nPAiD-Nucleoplasmin NLS-Uridine glycosylase inhibitor-SV40 NLS-E9 terminator-CaMV35S Promoter (enhanced)-HPT-CaMV Poly(A) Signal)

gacggccagtgccaagcttaaggaatctttaaacatacgaacagatcacttaaagttcttctgaagcaacttaaagttatcaggcatgcatggatcttggag
gaatcagatgtgcagtcagggaccatagcacaagacaggcgtcttctactggtgctaccagcaaatgctggaagccgggaacactgggtacgttggaa
accacgtgatgtgaagaagtaagataaactgtaggagaaaagcatttcgtagtgggccatgaagcctttcaggacatgtattgcagtatgggccggccca
ttacgcaattggacgacaacaaagactagtattagtaccacctcggctatccacatagatcaaagctgatttaaaagagttgtgcagatgatccgtggcag
gagaccgaggtctcggtttaagagctatgctggaaacagcatagcaagtttaaataaggctagtccgttatcaacttgaaaaagtggcaccgagtcggtg
cttttttgttttagagctagaaatagcaagttaaaataaggctagtgctagtgcagcgtgacccggtcgtgcccctcactagagataatgagcattgcatgtct
aagttataaaaaattaccacatattttttttgtcacacttgtttgaagtgcagtttatctatctttatacatatatttaaactttactctacgaataatataatctatagta
ctacaataatatcagtgttttagagaatcatataaatgaacagttagacatggtctaaaggacaattgagtattttgacaacaggactctacagttttatctttttta
gtgtgcatgtgttctccttttttttttgcaaatagcttcacctatataatacttcatccattttattagtacatccatttagggtttagggttaatggtttttatagactaatt
tttttagtacatctattttattctattttagcctctaaattaagaaaactaaaactctattttagtttttttatttaataatttagatataaaatagaataaaataaagtgac
taaaaattaaacaaataccctttaagaaattaaaaaaactaaggaaacatttttcttgtttcgagtagataatgccagcctgttaaacgccgtcgacgagtcta
acggacaccaaccagcgaaccagcagcgtcgcgtcgggccaagcgaagcagacggcacggcatctctgtcgctgcctctggacccctctcgagagtt
ccgctccaccgttggacttgctccgctgtcggcatccagaaattgcgtggcggagcggcagacgtgagccggcacggcaggcggcctcctcctcctct
cacggcaccggcagctacgggggattcctttcccaccgctccttcgctttcccttcctcgcccgccgtaataaatagacaccccctccacaccctctttccc
caacctcgtgttgttcggagcgcacacacacacaaccagatctcccccaaatccacccgtcggcacctccgcttcaaggtacgccgctcgtcctcccccc
ccccccctctctaccttctctagatcggcgttccggtccatgcttagggcccggtagttctacttctgttcatgtttgtgttagatccgtgtttgtgttagatccgt
gctgctagcgttcgtacacggatgcgacctgtacgtcagacacgttctgattgctaacttgccagtgtttctctttggggaatcctgggatggctctagccgtt
ccgcagacgggatcgatttcatgattttttttgtttcgttgcatagggtttggtttgcccttttcctttatttcaatatatgccgtgcacttgtttgtcgggtcatctttt
catgcttttttttgtcttggttgtgatgatgtggtctggttgggcggtcgttctagatcggagtacaattctgtttcaaactacctggtggatttattaattttggatct
gtatgtgtgtgccatacatattcatagttacgaattgaagatgatggatggaaatatcgatctaggataggtatacatgttgatgcgggttttactgatgcatat
acagagatgcttttttgttcgcttggttgtgatgatgtggtgtggttgggcggtcgttcattcgttctagatcggagtagaatactgtttcaaactacctggtgtatt
tattaattttggaactgtatgtgtgtgtcatacatcttcatagttacgagtttaagatggatggaaatatcgatctaggataggtatacatgttgatgtgggttttac
tgatgcatatacatgatggcatatgcagcatctattcatatgctctaaccttgagtacctatctattataataaacaagtatgttttataattattttgatcttgatata
cttggatgatggcatatgcagcagctatatgtggatttttttagccctgccttcatacgctatttatttgcttggtactgtttcttttgtcgatgctcaccctgttgttt
ggtgttacttctgcaggtacctaggaagccaccatggaggcctctccggcttcaggtcctcgccatctcatggacccccatatctttacctccaatttcaata
acggcattggcaggcacaagacatacctctgttatgaggtcgagcggctcgataacggaacctcagtgaagatggaccaacacaggggtttccttcata
accaagcgaaaaatctcctttgcggtttctatgggcggcacgcggagctgagattcttggatttggtcccgagcctccagctcgatcccgcgcaaatatac
cgcgtcacttggttcatatcatggagcccgtgctttagctgggggtgtgccggtgaggttcgcgcgttcctccaggaaaacacccatgttcgcctccggatt
ttcgcagcgaggattttcgactatgatccactttacaaagaggccctccaaatgcttagagatgctggggctcaagtttcaattatgacatacgacgagttta
agcattgttgggacacttttgtcgatcatcagggttgtcccttccagccgtgggatggtcttgatgagcattcgcaggctttgagtggcaggcttcgcgcgat
acttcagaaccagggtaactcaggttctgagacacctggcacaagtgagtcagcaacacccgagtccaccatggattacaaggaccacgacggggatt
acaaggaccacgacattgattacaaggatgatgatgacaagatggctccgaagaagaagaggaaggttggcatccacggggtgccagctgctatgaaa

aagccttactccatcggtcttgccatcggtactaacagcgtgggctgggccgttattactgatgattacaaggtgcctgcaaagaagatgaaggtgctcgg
caatacggacagatcacacatcaagaagaatctcattggtgcgctactttttgacgccgggaacactgctgaggataggcgcctgaagagaacggcgcg
gcgccgctatacaagacggaggaatagaatactgtatttgcaggaaatcttcgcagaagaaatgaacaagattgatgagtccttcttccaccggctcgac
gacagctttctcgtgcccgaggataaaaggggctccaagtatccaatatttgctactctgcaagaagagaaggagtaccacaagcagttccctaccatcta
ccatctccggaaacAATTGGCTGAATCTAATGAGAAGGCAGATCTGCGATTGGTATATCTTGCTCTG
GCTCACATGATCAAGTACAGAGGGCATTTTCTCATTGACGATCCCAAATTCAAAGTACAAAA
CAATGATATACAGGGTCTGTTTGAGAAATTTGTTGAAGAATACGACAATGTGCAGGAAACAT
CTTTGTCTAAGATAAAACTCAATGTCACAGAAATACTTACCGCCAAGATTCCGAAAAGTGAG
AAGCAAGAGCAGCTTCTCAAGAACTACCCATCGGAAAAGAAGAACACGTTATTTGGGAACT
TAATCGGCCTTGCGTTGGGCCTCACACCAAATTTCAAGACTAATTTTAGCTTGGAAAATGAC
GCTAAACTGCAAATCTCAAGCGAAAGTTATGAGGAGGACCTCGGCTCCCTCCTTGCACTAAT
TGGAGAGAACTTCATTGAGCTGTTCTCCGCGGTCAAGAACTTGAGTGATGGGATTCTCCTTG
CTGGGATAGTTTCAGATGAATCACCCCATGCTCCCCTCTCCACCAAAATGGTTATAAGGTTC
AAGGAGCACGAGGAAGACCTCGCCGCACTCAAGCATTTCATTAAGGCCAATTTGCCGGAGA
AGTACGATGAGGTTTTTTCAGATGACTCAAAGAACGGCTACGCCGGGTACGTCGGTGTCGAT
AGTAAGGTTCGCAAAAGGAACGGGAAATTAGCAACCGAGGAGGAGTTCTACAAATACCTCA
AGGATATTTTGAACAACGTGAAAGGCGCCGATTATTTCTTGGAGAAAATTAAACGGGAGGA
TTTACTCCGCAAACAGCGCACATTCGACAATGGGACTATCCCATATCAGGTTCATTTAGAAG
AGATGAAGGCCATTTTGCAAAACCAAGGCGAGTATTACCCCTTTCTTAAAGAGAACAAGGA
GAAGATTCAACAGATCCTGACATTCAGAATCCCGTACTACGTAGGACCTCTGGCACGCGGCA
ACCGCGACTTCGCGTGGTTGACGCGCAACTCAGATCAGGCGATTCGACCATGGAACTTTGAG
GAGGTCGTGGATAAGGCCTCCTCCGCGGAGGACTTCATCAACAAAATGACAAACTATGACC
TCTATCTACCAGAGGAAAAGGTCCTTCCCAAGCATTCGCTCTTGTACGAGACATTCGCTGTCT
ACAACGAGCTGACCAAAGTTAAGTTTATCGCTGAAGGACTGCGTGACTACCAATTCCTAGAC
TCCGGTCAAAAGAAACAGATAGTTAATCAGCTGTTTAAAGAAAAGAGAAAAGTGACTGAGA
AAGATATTATCCATTACCTCCACAACGTGGACGGTTATGATGGGATCGAATTAAAAGGAATC
GAGAAGCAGTTTAATGCTAGTCTGTCGACGTATCATGATCTCCTAAAAATCATTAAGGACAA
GGAGTTTATGGACGATCCTAAGAACGAGGAGATCCTCGAGAACATCGTCCATACACTCACG
ATATTCGAGGACCGCGAGATGATTAAGCAGAGGTTGGCTCAGTACGACTCTCTCTTTGATGA
GAAAGTCATCAAAGCTCTAACCCGTCGCCACTATACTGGCTGGGGCAAGTTATCCGCAAAGC
TGATAAATGGCATCTGTGATAAGCAAACAAATAAAACGATCCTGGACTTTCTTATTGACGAC
GACAAGATTAACAGAAACTTCATGCAGCTTATCAACGACGACGGCCTTTCTTTCAAGGATAT
AATTCAAAAGGCGCAGGTGGTCGGCAAGATCGACGATGTGAAGCAAGTTGTGCAGGAGCTT
CCAGGTTCTCCTGCGATTAAAAAGGGTATTTTGCAGTCGATAAAGATTGTTGATGAGTTGGT
CAAGGTCATGGGCCACGCCCCTGAGTCCATTGTTATCGAGATGGCCAGGGAGAATCAGACT
ACCGCCAGGGGCAAGAAAAACTCCCAGCAGAGATACAAGCGCATTGAGGACGCCTTAAAAA
ATCTAGCGCCGGGGTTGGATTCTAATATCCTCAAGGAAAACCCAACAGATAATATCCAGTTA
CAAAACGACCGCCTCTTCCTCTACTATCTTCAAAATGGTAAGGACATGTATACCGGTGAAGC
CCTCGACATAAATCAACTCAGCAATTATGATATAGATCATATTGTGCCGCAGGCGTTTATAA
AGGACGATAGTCTTGATAACAGGGTGCTTACTAGCAGCAAGGATAACAGGGGAAAGTCTGA
CAATGTGCCTTCTATAGAAGTTGTTCAGAAGCGTAAGGCCTTCTGGCAGCAACTTCTGGACT
CGAAATTGATAAGCGAGCGAAAGTTCAATAACCTCACCAAGGCAGAGCGCGGTGGACTCGA
CGAACGTGACAAAGTTGGGTTCATTAAGAGGCAGTTGGTTGAAACACGGCAGATTACAAAA
CATGTCGCTCAGATACTCGATGCCCGGTTCAACACGGAGGTAAACGAGAAGAACCAGAAGA
TAAGAAAGGTAAAAATTATTACTCTGAAATCCAACCTCGTTTCAAATTTCAGGAAAGAGTTC
GGCCTGTACAAGGTCCGTGAGATAAATGACTACCACCACGCCCACGATGCATACCTGAATGC
AGTCGTGGCGAAAGCGATCCTGAAGAAGTACCCTAAACTTGAACCGGAGTTCGTGTATGGA
GACTACCAGAAGTACGACCTTAAGAGGTACATCTCACGCTCCAAAGATCCAAAGGAAATAG
AGAAAGCAACAGAAAAGTACTTCTTTTATAGCAATTTGCTGAATTTTTCAAGGAAGAAGTG
CACTACGCTGATGGAACCATCATCAAGAGGGAGAATATAGAGTACTCTAAGGATACTGGCG
AGATAGCATGGAATAAAGAAAAGGATTTCGCCACTGTACGCAAGGTACTGAGCTGCCCTCA

GGTCAATATCGTGAAAAAGACCGAGGTTCAAACTGGCGGCTTCTCCAAGGAGTCAATCCTCC
CGAAGAGAAATTCTGACAAGTTAATCGCCCGAAAAAAGGACTGGGACCCGAAGAAGTATGG
AGGCTTCGATTCACCGACGGTGGCATATTCCGTGCTGGTGGTTGCCAAGGTGGAGAAGGGCA
AGTCAAAAAAGCTCAAGTCAGTAAAGGAACTTGTCGGAATCACGATCATGGAGCGGTCCTC
GTTCGAGAAGGACCCGGTCGACTTCCTGGAAGCAAAAGGGTACAAGGAGGTGAGAAAAGAT
TTGATTATCAAGCTCCCCAAGTATTCTCTTTTCGAGCTGGAAAATGGACGGAAGCGCATGctcg
cgtcggccggcgagcttcagaaaggaaatgaactggccctcccttcaaaatatgtcaacttcttatatctggcaagtcattatgaaaaactgaaagggagt
ccggaggataatgaacaaaaacaactgtttgtcgagcagcacaaacattacctcgatgagattattgagcaaatctccgagttctccaagagggtcattctc
gcggacgcaaatctcgacaaggtattatctgcctacaataaacatagagataagccaattagggaacaggcggagaacataatccacctcttcaccctca
cgaacctcggcgccccagccgcgtttaagtactttgacaccaccatcgacaggaaacgatacacttctacaaaagaagtgctcgatgccacgctgatcca
tcagagcattacaggtctttatgaaacccgtatagatctcagccaactcggaggggacaagcggccggcggccaccaagaaagctggccaagccaag
aagaagaaaggagacggatcaggcgggtcaacaaacctcagtgacatcatagagaaggaaactggtaagcaactggttattcaagagagcattctgat
gctccctgaggaagtcgaggaagttataggaaacaagcctgagagcgatatactcgtgcatacggcgtatgacgagagcacggatgaaaatgtgatgct
cttgaccagcgatgcgccagaatataaaccctgggcactggttattcaggactctaacggggaaaataaaatcaagatgctctcaggtggctccccaaag
aagaaacgcaaggtttaggctcagataatctagacccagctttcttgtacaaagtggttgataacagcgactacaaggatgacgatgacaaggcttagag
ctcagagctttcgttcgtatcatcggtttcgacaacgttcgtcaagttcaatgcatcagtttcattgcgcacacaccagaatcctactgagtttgagtattatgg
cattgggaaaactgtttttcttgtaccatttgttgtgcttgtaatttactgtgttttttattcggttttcgctatcgaactgtgaaatggaaatggatggagaagagtt
aatgaatgatatggtccttttgttcattctcaaattaatattatttgttttttctcttatttgttgtgtgttgaatttgaaattataagagatatgcaaacattttgttttgag
taaaaatgtgtcaaatcgtggcctctaatgaccgaagttaatatgaggagtaaaacacttgtagttgtaccattatgcttattcactaggcaacaaatatattttc
agacctagaaaagctgcaaatgttactgaatacaagtatgtcctcttgtgttttagacatttatgaactttcctttatgtaattttccagaatccttgtcagattctaa
tcattgctttataattatagttatactcatggatttgtagttgagtatgaaaatatttttaatgcattttatgacttgccaattgattgacaacgaattcgtaatcatgt
catagctgtttcctgtgtgaaattgttatccgctcacaattccacacaacatacgagccggaagcataaagtgtaaagcctggggtgcctaatgagtgagct
aactcacattaattgcgttgcgctcactgcccgctttccagtcgggaaacctgtcgtgccagctgcattaatgaatcggccaacgcgcggggagaggcgg
tttgcgtattggctagagcagcttgccaacatggtggagcacgacactctcgtctactccaagaatatcaaagatacagtctcagaagaccaaagggctat
tgagacttttcaacaaagggtaatatcgggaaacctcctcggattccattgcccagctatctgtcacttcatcaaaaggacagtagaaaaggaaggtggca
cctacaaatgccatcattgcgataaaggaaaggctatcgttcaagatgcctctgccgacagtggtcccaaagatggacccccacccacgaggagcatcg
tggaaaaagaagacgttccaaccacgtcttcaaagcaagtggattgatgtgataacatggtggagcacgacactctcgtctactccaagaatatcaaagat
acagtctcagaagaccaaagggctattgagacttttcaacaaagggtaatatcgggaaacctcctcggattccattgcccagctatctgtcacttcatcaaa
aggacagtagaaaaggaaggtggcacctacaaatgccatcattgcgataaaggaaaggctatcgttcaagatgcctctgccgacagtggtcccaaagat
ggacccccacccacgaggagcatcgtggaaaaagaagacgttccaaccacgtcttcaaagcaagtggattgatgtgatatctccactgacgtaagggat
gacgcacaatcccactatccttcgcaagaccttcctctatataaggaagttcatttcatttggagaggacacgctgaaatcaccagtctctctctacaaatcta
tctctctcgagctttcgcagatcccggggggcaatgagatatgaaaaagcctgaactcaccgcgacgtctgtcgagaagtttctgatcgaaaagttcgaca
gcgtctccgacctgatgcagctctcggagggcgaagaatctcgtgctttcagcttcgatgtaggagggcgtggatatgtcctgcgggtaaatagctgcgc
cgatggtttctacaaagatcgttatgtttatcggcactttgcatcggccgcgctcccgattccggaagtgcttgacattggggagtttagcgagagcctgacc
tattgcatctcccgccgtgcacagggtgtcacgttgcaagacctgcctgaaaccgaactgcccgctgttctacaaccggtcgcggaggctatggatgcga
tcgctgcggccgatcttagccagacgagcgggttcggcccattcggaccgcaaggaatcggtcaatacactacatggcgtgatttcatatgcgcgattgc
tgatccccatgtgtatcactggcaaactgtgatggacgacaccgtcagtgcgtccgtcgcgcaggctctcgatgagctgatgctttgggccgaggactgc
cccgaagtccggcacctcgtgcacgcggatttcggctccaacaatgtcctgacggacaatggccgcataacagcggtcattgactggagcgaggcgat
gttcggggattcccaatacgaggtcgccaacatcttcttctggaggccgtggttggcttgtatggagcagcagacgcgctacttcgagcggaggcatccg
gagcttgcaggatcgccacgactccgggcgtatatgctccgcattggtcttgaccaactctatcagagcttggttgacggcaatttcgatgatgcagcttgg
gcgcagggtcgatgcgacgcaatcgtccgatccggagccgggactgtcgggcgtacacaaatcgcccgcagaagcgcggccgtctggaccgatgg
ctgtgtagaagtactcgccgatagtggaaaccgacgccccagcactcgtccgagggcaaagaaatagagtagatgccgaccggatctgtcgatcgaca
agctcgagtttctccataataatgtgtgagtagttcccagataagggaattagggttcctatagggtttcgctcatgtgttgagcatataagaaacccttagtat
gtatttgtatttgtaaaatacttctatcaataaaatttctaattcctaaaaccaaaatccagtactaaaatccagatccccgaattaattcggcg

Seq ID No. 6

(Composit Promoter-tGly-AmpR Promoter-AmpR-sgRNA2m-HDV ribozyme-PolyT HSPt-ZmUbi Promoter-SV40 NLS-nPAiD-Linker-M MLV RT-NLSc Myc-E9 terminator-CaMV35S Promoter (enhanced)-HPT-CaMV Poly(A) Signal)

ccaagcttatggagtcaaagattcaaatagaggacctaacagaactcgccgtaaagactggcgaacagttcatacagagtctcttacgactcaatgacaa
gaagaaaatcttcgtcaacatggtggagcacgacacacttgtctactccaaaaatatcaaagatacagtctcagaagaccaaagggcaattgagacttttc
aacaaagggtaatatccggaaacctcctcggattccattgcccagctatctgtcactttattgtgaagatagtggaaaaggaaggtggctcctacaaatgcc
atcattgcgataaaggaaaggccatcgttgaagatgcctctgccgacagtggtcccaaagatggacccccacccacgaggagcatcgtggaaaaagaa
gacgttccaaccacgtcttcaaagcaagtggattgatgtgattggcagacatactgtcccacaaatgaagatggaatctgtaaaagaaaacgcgtgaaata
atgcgtctgacaaaggttaggtcggctgcctttaatcaataccaaagtggtccctaccacgatggaaaaactgtgcagtcggtttggctttttctgacgaaca
aataagattcgtggccgacaggtgggggtccaccatgtgaaggcatcttcagactccaataatggagcaatgacgtaagggcttacgaaataagtaagg
gtagtttgggaaatgtccactcacccgtcagtctataaatacttagcccctccctcattgttaagggagcaaaatctcagagagatagtcctagagagagaa
agagagcaagtagcctagaagtagtcaaggcggcgaagtattcaggcacgtggccaggaagaagaaaagccaagacgacgaaaacaggtaagagc
taagcatctaggtaagttgaaaacaatcttcaaaagtcccacatcgcttagataagaaaacgaagctgagtttatatacagctagagtcgaagtagtgattg
aacaaagcaccagtggtctagtggtagaatagtaccctgccacggtacagacccgggttcgattcccggctggtgcatgagaccttcggggaaatgtgc
gcggaacccctatttgtttatttttctaaatacattcaaatatgtatccgctcatgggacaataaccctgataaatgcttcaataatattgaaaaaggaagagtat
gagtattcaacatttccgtgtcgcccttattcccttttttgcggcattttgccttcctgtttttgctcacccagaaacgctggtgaaagtaaaagatgctgaagatc
agttgggtgcacgagtgggttacatcgaactggatctcaacagcggtaagatccttgagagttttcgccccgaagaacgttttccaatgatgagcactttta
aagttctgctatgtggcgcggtattatcccgtattgacgccgggcaagagcaactcggtcgccgcatacactattctcagaatgacttggttgagtactcac
cagtcacagaaaagcatcttacggatggcatgacagtaagagaattatgcagtgctgccataaccatgagtgataacactgcggccaacttacttctgaca
acgatcggaggaccgaaggagctaaccgcttttttgcacaacatgggggatcatgtaactcgccttgatcgttgggaaccggagctgaatgaagccatac
caaacgacgagcgtgacaccacgatgcctgtagcaatggcaacaacgttgcgcaaactattaactggcgaactacttactctagcttcccggcaacaatt
aatagactggatggaggcggataaagttgcaggaccacttctgcgctcggcccttccggctggctggtttattgctgataaatctggagccggtgagcgt
ggctctcgcggtatcattgcagcactggggccagatggtaagccctcccgtatcgtagttatctacacgacggggagtcaggcaactatggatgaacgaa
atagacagatcgctgagataggtgcctcactgattaagcattggtaactgtcagaccaagtttactcatatatactttagattgatttaaaacttcatttttaattta
aaaggatctaggtgaagatcctttttgataatcccatgaccaaaatcccttaacgtgagttttcgttccactgagcgtcaggtctcagtttcagagctatgctg
gaaacagcatagcaagttgaaataaggctagtccgttatcaacttgaaaaagtggcaccgagtcggtgcttttttttggccggcatggtcccagcctcctcg
ctggcgccggctgggcaacatgcttcggcatggcgaatgggacttttttttgatatctccggggctaattgaatatgaagatgaagatgaaatatttggtgtg
tcaaataaaaagctggtgtgcttaagtttgtgttttttttcttggcttgttgtgttatgaatttgtggctttttctaatattaaatgaatgtaagatctcattataatgaata
aacaaatgtttctataatccattgtgaatgttttgttggatctcttctgcagcatataactactgtatgtgctatggtatggactatggaatatgattaaagataact
agtgcagcgtgacccggtcgtgcccctcactagagataatgagcattgcatgtctaagttataaaaaattaccacatattttttttgtcacacttgtttgaagtg
cagtttatctatctttatacatatatttaaactttactctacgaataatataatctatagtactacaataatatcagtgttttagagaatcatataaatgaacagttaga
catggtctaaaggacaattgagtattttgacaacaggactctacagttttatctttttagtgtgcatgtgttctcctttttttttgcaaatagcttcacctatataatac
ttcatccattttattagtacatccatttagggtttagggttaatggtttttatagactaatttttttagtacatctattttattctattttagcctctaaattaagaaaactaa
aactctattttagtttttttatttaataatttagatataaaatagaataaaataaagtgactaaaaattaaacaaataccctttaagaaattaaaaaaactaaggaaa
catttttcttgtttcgagtagataatgccagcctgttaaacgccgtcgacgagtctaacggacaccaaccagcgaaccagcagcgtcgcgtcgggccaag
cgaagcagacggcacggcatctctgtcgctgcctctggacccctctcgagagttccgctccaccgttggacttgctccgctgtcggcatccagaaattgc
gtggcggagcggcagacgtgagccggcacggcaggcggcctcctcctcctctcacggcaccggcagctacgggggattcctttcccaccgctccttc
gctttcccttcctcgcccgccgtaataaatagacacccccctccacaccctctttccccaacctcgtgttgttcggagcgcacacacacacaaccagatctcc
cccaaatccacccgtcggcacctccgcttcaaggtacgccgctcgtcctcccccccccccccctctctaccttctctagatcggcgttccggtccatgcttag
ggcccggtagttctacttctgttcatgtttgtgttagatccgtgtttgtgttagatccgtgctgctagcgttcgtacacggatgcgacctgtacgtcagacacgtt
ctgattgctaacttgccagtgtttctctttggggaatcctgggatggctctagccgttccgcagacgggatcgatttcatgattttttttgtttcgttgcatagggt
ttggtttgcccttttcctttatttcaatatatgccgtgcacttgtttgtcgggtcatcttttcatgcttttttttgtcttggttgtgatgatgtggtctggttgggcggtcg
ttctagatcggagtacaattctgtttcaaactacctggtggatttattaattttggatctgtatgtgtgtgccatacatattcatagttacgaattgaagatgatgga
tggaaatatcgatctaggataggtatacatgttgatgcgggttttactgatgcatatacagagatgctttttgttcgcttggttgtgatgatgtggtgtggttggg
cggtcgttcattcgttctagatcggagtagaatactgtttcaaactacctggtgtatttattaattttggaactgtatgtgtgtgtcatacatcttcatagttacgagt
ttaagatggatggaaatatcgatctaggataggtatacatgttgatgtgggttttactgatgcatatacatgatggcatatgcagcatctattcatatgctctaac
cttgagtacctatctattataataaacaagtatgttttataattattttgatcttgatatacttggatgatggcatatgcagcagctatatgtggatttttttagccctg
ccttcatacgctatttatttgcttggtactgtttcttttgtcgatgctcaccctgttgtttggtgttacttctgcaggtacCTAGGAAGCCACCATGG
ATTATAAAGATCACGACGGCGACTACAAAGATCATGACATCGATTACAAGGACGACGACGA
CAAGATGAAACGTACAGCTGATGGCAGCGAATTTGAATCGCCGAAGAAGAAGCGGAAGGTG
ATGAAAAAGCCTTACTCCATCGGTCTTGACATCGGTACTAACAGCGTGGGCTGGGCCGTTAT

TACTGATGATTACAAGGTGCCTGCAAAGAAGATGAAGGTGCTCGGCAATACGGACAGATCA
CACATCAAGAAGAATCTCATTGGTGCGCTACTTTTTGACGCCGGGAACACTGCTGAGGATAG
GCGCCTGAAGAGAACGGCGCGGCGCCGCTATACAAGACGGAGGAATAGAATACTGTATTTG
CAGGAAATCTTCGCAGAAGAAATGAACAAGATTGATGAGTCCTTCTTCCACCGGCTCGACGA
CAGCTTTCTCGTGCCCGAGGATAAAAGGGGCTCCAAGTATCCAATATTTGCTACTCTGCAAG
AAGAGAAGGAGTACCACAAGCAGTTCCCTACCATCTACCATCTCCGGAAACAATTGGCTGA
ATCTAATGAGAAGGCAGATCTGCGATTGGTATATCTTGCTCTGGCTCACATGATCAAGTACA
GAGGGCATTTTCTCATTGACGATCCCAAATTCAAAGTACAAAACAATGATATACAGGGTCTG
TTTGAGAAATTTGTTGAAGAATACGACAATGTGCAGGAAACATCTTTGTCTAAGATAAAACT
CAATGTCACAGAAATACTTACCGCCAAGATTCCGAAAAGTGAGAAGCAAGAGCAGCTTCTC
AAGAACTACCCATCGGAAAAGAAGAACACGTTATTTGGGAACTTAATCGGCCTTGCGTTGG
GCCTCACACCAAATTTCAAGACTAATTTTAGCTTGGAAAATGACGCTAAACTGCAAATCTCA
AGCGAAAGTTATGAGGAGGACCTCGGCTCCCTCCTTGCACTAATTGGAGAGAACTTCATTGA
GCTGTTCTCCGCGGTCAAGAACTTGAGTGATGGGATTCTCCTTGCTGGGATAGTTTCAGATG
AATCACCCCATGCTCCCCTCTCCACCAAAATGGTTATAAGGTTCAAGGAGCACGAGGAAGAC
CTCGCCGCACTCAAGCATTTCATTAAGGCCAATTTGCCGGAGAAGTACGATGAGGTTTTTTC
AGATGACTCAAAGAACGGCTACGCCGGGTACGTCGGTGTCGATAGTAAGGTTCGCAAAAGG
AACGGGAAATTAGCAACCGAGGAGGAGTTCTACAAATACCTCAAGGATATTTTGAACAACG
TGAAAGGCGCCGATTATTTCTTGGAGAAAATTAAACGGGAGGATTTACTCCGCAAACAGCG
CACATTCGACAATGGGACTATCCCATATCAGGTTCATTTAGAAGAGATGAAGGCCATTTTGC
AAAACCAAGGCGAGTATTACCCCTTTCTTAAAGAGAACAAGGAGAAGATTCAACAGATCCT
GACATTCAGAATCCCGTACTACGTAGGACCTCTGGCACGCGGCAACCGCGACTTCGCGTGGT
TGACGCGCAACTCAGATCAGGCGATTCGACCATGGAACTTTGAGGAGGTCGTGGATAAGGC
CTCCTCCGCGGAGGACTTCATCAACAAAATGACAAACTATGACCTCTATCTACCAGAGGAAA
AGGTCCTTCCCAAGCATTCGCTCTTGTACGAGACATTCGCTGTCTACAACGAGCTGACCAAA
GTTAAGTTTATCGCTGAAGGACTGCGTGACTACCAATTCCTAGACTCCGGTCAAAAGAAACA
GATAGTTAATCAGCTGTTTAAAGAAAAGAGAAAAGTGACTGAGAAAGATATTATCCATTAC
CTCCACAACGTGGACGGTTATGATGGGATCGAATTAAAAGGAATCGAGAAGCAGTTTAATG
CTAGTCTGTCGACGTATCATGATCTCCTAAAAATCATTAAGGACAAGGAGTTTATGGACGAT
CCTAAGAACGAGGAGATCCTCGAGAACATCGTCCATACACTCACGATATTCGAGGACCGCG
AGATGATTAAGCAGAGGTTGGCTCAGTACGACTCTCTCTTTGATGAGAAAGTCATCAAAGCT
CTAACCCGTCGCCACTATACTGGCTGGGGCAAGTTATCCGCAAAGCTGATAAATGGCATCTG
TGATAAGCAAACAAATAAAACGATCCTGGACTTTCTTATTGACGACGACAAGATTAACAGA
AACTTCATGCAGCTTATCAACGACGACGGCCTTTCTTTCAAGGATATAATTCAAAAGGCGCA
GGTGGTCGGCAAGATCGACGATGTGAAGCAAGTTGTGCAGGAGCTTCCAGGTTCTCCTGCGA
TTAAAAAGGGTATTTTGCAGTCGATAAAGATTGTTGATGAGTTGGTCAAGGTCATGGGCCAC
GCCCCTGAGTCCATTGTTATCGAGATGGCCAGGGAGAATCAGACTACCGCCAGGGGCAAGA
AAAACTCCCAGCAGAGATACAAGCGCATTGAGGACGCCTTAAAAAATCTAGCGCCGGGGTT
GGATTCTAATATCCTCAAGGAAAACCCAACAGATAATATCCAGTTACAAAACGACCGCCTCT
TCCTCTACTATCTTCAAAATGGTAAGGACATGTATAccggtgaagccctcgacataaatcaactcagcaattatgatata
gatgccattgtgccgcaggcgtttataaaggacgatagtcttgataacagggtgcttactagcagcaaggataacaggggaaagtctgacaatgtgccttc
tatagaagttgttcagaagcgtaaggccttctggcagcaacttctggactcgaaattgataagcgagcgaaagttcaataacctcaccaaggcagagcgc
ggtggactcgacgaacgtgacaaagttgggttcattaagaggcagttggttgaaacacggcagattacaaaacatgtcgctcagatactcgatgcccggt
tcaacacggaggtaaacgagaagaaccagaagataagaaaggtaaaaattattactctgaaatccaacctcgtttcaaatttcaggaaagagttcggcct
GTACAAGGTCCGTGAGATAAATGACTACCACCACGCCCACGATGCATACCTGAATGCAGTC
GTGGCGAAAGCGATCCTGAAGAAGTACCCTAAACTTGAACCGGAGTTCGTGTATGGAGACT
ACCAGAAGTACGACCTTAAGAGGTACATCTCACGCTCCAAAGATCCAAAGGAAATAGAGAA
AGCAACAGAAAAGTACTTCTTTTATAGCAATTTGCTGAATTTTTTCAAGGAAGAAGTGCACT
ACGCTGATGGAACCATCATCAAGAGGGAGAATATAGAGTACTCTAAGGATACTGGCGAGAT
AGCATGGAATAAAGAAAAGGATTTCGCCACTGTACGCAAGGTACTGAGCTGCCCTCAGGTC
AATATCGTGAAAAAGACCGAGGTTCAAACTGGCGGCTTCTCCAAGGAGTCAATCCTCCCGA

AGAGAAATTCTGACAAGTTAATCGCCCGAAAAAAGGACTGGGACCCGAAGAAGTATGGAGG
CTTCGATTCACCGACGGTGGCATATTCCGTGCTGGTGGTTGCCAAGGTGGAGAAGGGCAAGT
CAAAAAAGCTCAAGTCAGTAAAGGAACTTGTCGGAATCACGATCATGGAGCGGTCCTCGTT
CGAGAAGGACCCGGTCGACTTCCTGGAAGCAAAAGGGTACAAGGAGGTGAGAAAAGATTTG
ATTATCAAGCTCCCCAAGTATTCTCTTTTCGAGCTGGAAAATGGACGGAAGCGCATGctcgcgtc
ggccggcgagcttcagaaaggaaatgaactggccctcccttcaaaatatgtcaacttcttatatctggcaagtcattatgaaaaactgaaagggagtccgg
aggataatgaacaaaaacaactgtttgtcgagcagcacaaacattacctcgatgagattattgagcaaatctccgagttctccaagagggtcattctcgcgg
acgcaaatctcgacaaggtattatctgcctacaataaacatagagataagccaattagggaacaggcggagaacataatccacctcttcaccctcacgaa
cctcggcgccccagccgcgtttaagtactttgacaccaccatcgacaggaaacgatacacttctacaaaagaagtgctcgatgccacgctgatccatcag
agcattacaggtctttatgaaacccgtatagatctcagccaactcggaggggactcaggcggctcatctggcgggtcaaagcgcacagccgacggctct
gagttcgagagccctaagaagaagcgcaaggtgtcaggcggctcttcaggcggcagcaccctgaacattgaggacgagtaccggctgcacgagacg
agcaaggagccagacgtttcgctcggcagcacttggctctctgacttcccacaggcttgggccgagactggcggcatgggcctggccgtgcgccagg
ctccactgatcatccctctgaaggcgacctccaccccggtttctattaagcagtacccgatgagccaggaggccaggctggggatcaagccacacattca
gcggctgctggaccagggcatcctggtgccatgccagtccccgtggaatactccgctcctgccggtgaagaagcctgggacaaacgactacaggccg
gttcaggatctcagggaggtgaacaagcgcgtggaggacatccatccgacagtgccgaacccgtacaatctgctgtcgggcctgcctccgagccacca
gtggtacaccgtcctggacctcaaggacgctttcttctgcctgcggctgcacccgacgtctcagccgctgttcgcgttcgagtggcgcgacccagagatg
ggcatttccggccagctgacctggacacgcctaccccagggcttcaagaactccccgactctcttcaacgaggctctccaccgggatctcgcggacttca
ggattcagcatcccgatctgatcctgctccagtatgttgacgacctcctcctggccgcgacgtcggagctggactgccagcagggcacccgggcgctgc
tgcagacactgggcaatctggggtaccgcgcctctgcgaagaaggcgcagatctgccagaagcaagtgaagtacctgggctacctcctgaaggaggg
ccagcgctggctcactgaggcgaggaaggagactgttatgggccagcccactccaaagactccgaggcagctcagggagttcctcggcaaggctgg
gttctgccgcctgttcatccctgggttcgctgagatggctgcgccgctctacccgctgactaagccggggacactgttcaactgggggccagaccagca
gaaggcgtaccaggagattaagcaggcgctgctgacggccccagcgctcggcctaccagacctgacgaagccgttcgagctgttcgttgacgagaag
caggggtacgcgaagggcgtgctgacacagaagctggggccttggcgccgcccggtcgcgtacctgtcgaagaagctggacccagtcgctgctggg
tggcctccatgcctccggatggtcgctgctattgcggttctgaccaaggatgcggggaagctcacaatggggcagcctctcgtgatcctggctccacatg
cggtggaggcgctggtgaagcagccaccggaccggtggctgtcgaacgctcggatgacacactaccaggcgctcctcctcgatacagaccgggttca
gttcgggcctgtggttgctctgaacccagccacactgctgccactccctgaggagggcctccagcacaattgcctcgacatcctggctgaggcgcacgg
cacccgccctgatctcaccgaccagcctctgccagatgctgaccacacctggtacacggatgggtcctcgctgctgcaggagggccagaggaaggcg
ggcgccgccgtcaccacagagacagaggttatttgggccaaggccctaccggctggcaccagcgcccagcgcgctgagctgatcgcgctgactcag
gcgctgaagatggccgaggggaagaagctcaatgtttacaccgactcgcggtacgcgttcgctacagctcacattcatggggagatctaccgccggcg
cgggtggctgacttcggagggcaaggagattaagaataaggacgagatcctggccctgctcaaggcgctgttcctgccgaagcgcctctcaatcattca
ctgcccgggccaccagaagggccattcggccgaggctaggggcaatcggatggctgaccaggcggcgcggaaggcggctatcaccgagactcccg
atacatctaccctcctgatcgagaactcgagcccatctggcggctctaagcggactgcggatgggtctgagttcgagtcaccaaagaagaagaggaag
gtgggctctggccctgctgctaagcgcgtgaagctcgattgagctcagagctttcgttcgtatcatcggtttcgacaacgttcgtcaagttcaatgcatcagt
ttcattgcgcacacaccagaatcctactgagtttgagtattatggcattgggaaaactgtttttcttgtaccatttgttgtgcttgtaatttactgtgtttttattcggt
tttcgctatcgaactgtgaaatggaaatggatggagaagagttaatgaatgatatggtccttttgttcattctcaaattaatattatttgtttttctcttatttgttgtg
tgttgaatttgaaattataagagatatgcaaacattttgttttgagtaaaaatgtgtcaaatcgtggcctctaatgaccgaagttaatatgaggagtaaaacactt
gtagttgtaccattatgcttattcactaggcaacaaatatattttcagacctagaaaagctgcaaatgttactgaatacaagtatgtcctcttgtgttttagacattt
atgaactttcctttatgtaattttccagaatccttgtcagattctaatcattgctttataattatagttatactcatggatttgtagttgagtatgaaaatattttttaatgc
attttatgacttgccaattgattgacaacgaattcgtaatcatgtcatagctgtttcctgtgtgaaattgttatccgctcacaattccacacaacatacgagccgg
aagcataaagtgtaaagcctggggtgcctaatgagtgagctaactcacattaattgcgttgcgctcactgcccgctttccagtcgggaaacctgtcgtgcc
agctgcattaatgaatcggccaacgcgcggggagaggcggtttgcgtattggctagagcagcttgccaacatggtggagcacgacactctcgtctactc
caagaatatcaaagatacagtctcagaagaccaaagggctattgagacttttcaacaaagggtaatatcgggaaacctcctcggattccattgcccagcta
tctgtcacttcatcaaaaggacagtagaaaaggaaggtggcacctacaaatgccatcattgcgataaaggaaaggctatcgttcaagatgcctctgccga
cagtggtcccaaagatggacccccacccacgaggagcatcgtggaaaaagaagacgttccaaccacgtcttcaaagcaagtggattgatgtgataacat
ggtggagcacgacactctcgtctactccaagaatatcaaagatacagtctcagaagaccaaagggctattgagacttttcaacaaagggtaatatcggga
aacctcctcggattccattgcccagctatctgtcacttcatcaaaaggacagtagaaaaggaaggtggcacctacaaatgccatcattgcgataaaggaaa
ggctatcgttcaagatgcctctgccgacagtggtcccaaagatggacccccacccacgaggagcatcgtggaaaaagaagacgttccaaccacgtcttc
aaagcaagtggattgatgtgatatctccactgacgtaagggatgacgcacaatcccactatccttcgcaagacccttcctctatataaggaagttcatttcattt
ggagaggacacgctgaaatcaccagtctctctctacaaatctatctctctcgagctttcgcagatcccggggggcaatgagatatgaaaaagcctgaactc
accgcgacgtctgtcgagaagtttctgatcgaaaagttcgacagcgtctccgacctgatgcagctctcggagggcgaagaatctcgtgctttcagcttcga
tgtaggagggcgtggatatgtcctgcgggtaaatagctgcgccgatggtttctacaaagatcgttatgtttatcggcactttgcatcggccgcgctcccgatt

ccggaagtgcttgacattggggagtttagcgagagcctgacctattgcatctcccgccgtgcacagggtgtcacgttgcaagacctgcctgaaaccgaac
tgcccgctgttctacaaccggtcgcggaggctatggatgcgatcgctgcggccgatcttagccagacgagcgggttcggcccattcggaccgcaagga
atcggtcaatacactacatggcgtgatttcatatgcgcgattgctgatccccatgtgtatcactggcaaactgtgatggacgacaccgtcagtgcgtccgtc
gcgcaggctctcgatgagctgatgctttgggccgaggactgccccgaagtccggcacctcgtgcacgcggatttcggctccaacaatgtcctgacggac
aatggccgcataacagcggtcattgactggagcgaggcgatgttcggggattcccaatacgaggtcgccaacatcttcttctggaggccgtggttggctt
gtatggagcagcagacgcgctacttcgagcggaggcatccggagcttgcaggatcgccacgactccgggcgtatatgctccgcattggtcttgaccaac
tctatcagagcttggttgacggcaatttcgatgatgcagcttgggcgcagggtcgatgcgacgcaatcgtccgatccggagccgggactgtcgggcgta
cacaaatcgcccgcagaagcgcggccgtctggaccgatggctgtgtagaagtactcgccgatagtggaaaccgacgccccagcactcgtccgaggg
caaagaaatagagtagatgccgaccggatctgtcgatcgacaagctcgagtttctccataataatgtgtgagtagttcccagataagggaattagggttcct
atagggtttcgctcatgtgttgagcatataagaaacccttagtatgtatttgtatttgtaaaatacttctatcaataaaatttctaattcctaaaaccaaaatccagt
actaaaatccagatccccccgaattaatt

Seq ID No. 7

(CaMV Poly(A) Signal-HPT-CaMV35s Promoter (enhanced)-OsU3 Promoter-tRNA-crRNA1-tracrRNA-tRNA-crRNA2-tracrRNA-PolyT-OsUbi10 Promoter-SV40 NLS-PAiD-Nucleoplasmin NLS-NOS T)

aattcggggggatctggattttagtactggattttggttttaggaattagaaattttattgatagaagtattttacaaatacaaatacatactaagggtttcttatatgc
tcaacacatgagcgaaaccctataggaaccctaattcccttatctgggaactactcacacattattatggagaaactcgagcttgtcgatcgacagatcccg
gtcggcatctactctatttctttgccctcggacgagtgctggggcgtcggtttccactatcggcgagtacttctacacagccatcggtccagacggccgcgc
ttctgcgggcgatttgtgtacgcccgacagtcccggctccggatcggacgattgcgtcgcatcgaccctgcgcccaagctgcatcatcgaaattgccgtc
aaccaagctctgatagagttggtcaagaccaatgcggagcatatacgcccggagtcgtggcgatcctgcaagctccggatgcctccgctcgaagtagc
gcgtctgctgctccatacaagccaaccacggcctccagaagaagatgttggcgacctcgtattgggaatccccgaacatcgcctcgctccagtcaatgac
cgctgttatgcggccattgtccgtcaggacattgttggagccgaaatccgcgtgcacgaggtgccggacttcggggcagtcctcggcccaaagcatcag
ctcatcgagagcctgcgcgacggacgcactgacggtgtcgtccatcacagtttgccagtgatacacatggggatcagcaatcgcgcatatgaaatcacg
ccatgtagtgtattgaccgattccttgcggtccgaatgggccgaacccgctcgtctggctaagatcggccgcagcgatcgcatccatagcctccgcgacc
ggttgtagaacagcgggcagttcggtttcaggcaggtcttgcaacgtgacaccctgtgaacggcgggagatgcaataggtcaggctctcgctaaactcc
ccaatgtcaagcacttccggaatcgggagcgcggccgatgcaaagtgccgataaacataacgatctttgtagaaaccatcggcgcagctatttacccgc
aggacatatccacgccctcctacatcgaagctgaaagcacgagattcttcgccctccgagagctgcatcaggtcggagacgctgtcgaacttttcgatca
gaaacttctcgacagacgtcgcggtgagttcaggctttttcatatctcattgccccccggatctgcgaaagctcgagagagatagatttgtagagagagact
ggtgatttcagcgtgtcctctccaaatgaaatgaacttccttatatagaggaagggtcttgcgaaggatagtgggattgtgcgtcatcccttacgtcagtgga
gatatcacatcaatccacttgctttgaagacgtggttggaacgtcttctttttccacgatgctcctcgtgggtgggggtccatctttgggaccactgtcggcag
aggcatcttgaacgatagcctttcctttatcgcaatgatggcatttgtaggtgccaccttccttttctactgtccttttgatgaagtgacagatagctgggcaatg
gaatccgaggaggtttcccgatattaccctttgttgaaaagtctcaatagccctttggtcttctgagactgtatctttgatattcttggagtagacgagagtgtcg
tgctccaccatgttcacatcaatccacttgctttgaagacgtggttggaacgtcttctttttccacgatgctcctcgtgggtgggggtccatctttgggaccact
gtcggcagaggcatcttgaacgatagcctttcctttatcgcaatgatggcatttgtaggtgccaccttccttttctactgtccttttgatgaagtgacagatagct
gggcaatggaatccgaggaggtttcccgatattaccctttgttgaaaagtctcaatagccctttggtcttctgagactgtatctttgatattcttggagtagacg
agagtgtcgtgctccaccatgttggcaagctgctctagccaatacgcaaaccgcctctccccgcgcgttggccgattcattaatgcagctggcacgacag
gtttcccgactggaaagcgggcagtgagcgcaacgcaattaatgtgagttagctcactcattaggcaccccaggctttacactttatgcttccggctcgtat
gttgtgtggaattgtgagcggataacaatttcacacaggaaacagctatgacatgattacgccaagcttaaggaatctttaaacatacgaacagatcactta
aagttcttctgaagcaacttaaagttatcaggcatgcatggatcttggaggaatcagatgtgcagtcagggaccatagcacaagacaggcgtcttctactg
gtgctaccagcaaatgctggaagccgggaacactgggtacgttggaaaccacgtgatgtgaagaagtaagataaactgtaggagaaaagcatttcgtag
tgggccatgaagcctttcaggacatgtattgcagtatgggccggcccattacgcaattggacgacaacaaagactagtattagtaccacctcggctatcca
catagatcaaagctgatttaaaagagttgtgcagatgatccgtGGCAGGATGGGCAGTCTGGGCAACAAAGCACCAGT
GGtctagtggtagaatagtaccctgccacggtacagacccgggttcgattcccggctggtgcaNNNNNNNNNNNNNNNNNNNNgt
tttagagctagaaatagcaagttaaaataaggctagtccgttatcaacttgaaaaagtggcaccgagtcggtgcaacaaagcaccagtggtctagtggtag
aatagtaccctgccacggtacagacccgggttcgattcccggctggtgcaNNNNNNNNNNNNNNNNNNNNgttttagagctagaaat
agcaagttaaaataaggctagtccgttatcaacttgaaaaagtggcacCGAGTCGGTGCTTTTTTTTTTGTTTGCTGTCGCT
CATCCgttttagagctagaaatagcaagttaaaataaggctagtccgttatcaacttgaaaaagtggcaccgagtcggtgctttttgttttagagctaga

aatagcaagttaaaataaggctagtccgtttttagcgcgtgcatgcctgcaggtccacaaattcgggtcaaggcggaagccagcgcgccaccccacgtc
agcaaatacggaggcgcggggttgacggcgtcacccggtcctaacggcgaccaacaaaccagccagaagaaattacagtaaaaaaaaagtaaattgc
actttgatccaccttttattacctaagtctcaatttggatcacccttaaacctatcttttcaatttgggccgggttgtggtttggactaccatgaacaacttttcgtca
tgtctaacttccctttcagcaaacatatgaaccatatatagaggagatcggccgtatactagagctgatgtgtttaaggtcgttgattgcacgagaaaaaaa
atccaaatcgcaacaatagcaaatttatctggttcaaagtgaaaagatatgtttaaaggtagtccaaagtaaaacttatagataataaaatgtggtccaaagc
gtaattcactcaaaaaaaatcaacgagacgtgtaccaaacggagacaaacggcatcttctcgaaatttcccaaccgctcgctcgcccgcctcgtcttcccg
gaaaccgcggtggtttcagcgtggcggattctccaagcagacggagacgtcacggcacgggactcctcccaccacccaaccgccataaataccagcc
ccctcatctcctctcctcgcatcagctccacccccgaaaaatttctccccaatctcgcgaggctctcgtcgtcgaatcgaatcctctcgcgtcctcaaggtac
gctgcttctcctctcctcgcttcgtttcgattcgatttcggacgggtgaggttgttttgttgctagatccgattggtggttagggttgtcgatgtgattatcgtgag
atgtttaggggttgtagatctgatggttgtgatttgggcacggttggttcgataggtggaatcgtggttaggttttgggattggatgttggttctgatgattggg
gggaatttttacggttagatgaattgttggatgattcgattggggaaatcggtgtagatctgttggggaattgtggaactagtcatgcctgagtgattggtgcg
atttgtagcgtgttccatcttgtaggccttgttgcgagcatgttcagatctactgttccgctcttgattgagttattggtgccatgggttggtgcaaacacaggct
ttaatatgttatatctgttttgtgtttgatgtagatctgtagggtagttcttcttagacatggttcaattatgtagcttgtgcgtttcgatttgatttcatatgttcacaga
ttagataatgatgaactcttttaattaattgtcaatggtaaataggaagtcttgtcgctatatctgtcataatgatctcatgttactatctgccagtaatttatgctaa
gaactatattagaatatcatgttacaatctgtagtaatatcatgttacaatctgtagttcatctatataatctattgtggtaatttctttttactatctgtgtgaagattat
tgccactagttcattctacttatttctgaagttcaggatacgtgtgctgttactacctatctgaatacatgtgtgatgtgcctgttactatctttttgaatacatgtatg
ttctgttggaatatgtttgctgtttgatccgttgttgtgtccttaatcttgtgctagttcttaccctatctgtttggtgattatttcttgcagatagttatcaacaagtttgt
acaaaaaagcaggcttcgaaggagatagaaccaattctctaaggaaatacttaaccatggactataaggaccacgacggagactacaaggatcatgata
ttgattacaaagacgatgacgataagatggccccaaagaagaagcggaaggtcggtatccacggagtcccagcagccAAAAAGCCTTACT
CCATCGGTCTTGACATCGGTACTAACAGCGTGGGCTGGGCCGTTATTACTGATGATTACAAG
GTGCCTGCAAAGAAGATGAAGGTGCTCGGCAATACGGACAGATCACACATCAAGAAGAATC
TCATTGGTGCGCTACTTTTTGACGCCGGGAACACTGCTGAGGATAGGCGCCTGAAGAGAACG
GCGCGGCGCCGCTATACAAGACGGAGGAATAGAATACTGTATTTGCAGGAAATCTTCGCAG
AAGAAATGAACAAGATTGATGAGTCCTTCTTCCACCGGCTCGACGACAGCTTTCTCGTGCCC
GAGGATAAAAGGGGCTCCAAGTATCCAATATTTGCTACTCTGCAAGAAGAGAAGGAGTACC
ACAAGCAGTTCCCTACCATCTACCATCTCCGGAAACAATTGGCTGAATCTAATGAGAAGGCA
GATCTGCGATTGGTATATCTTGCTCTGGCTCACATGATCAAGTACAGAGGGCATTTTCTCATT
GACGATCCCAAATTCAAAGTACAAAACAATGATATACAGGGTCTGTTTGAGAAATTTGTTGA
AGAATACGACAATGTGCAGGAAACATCTTTGTCTAAGATAAAACTCAATGTCACAGAAATA
CTTACCGCCAAGATTCCGAAAAGTGAGAAGCAAGAGCAGCTTCTCAAGAACTACCCATCGG
AAAAGAAGAACACGTTATTTGGGAACTTAATCGGCCTTGCGTTGGGCCTCACACCAAATTTC
AAGACTAATTTTAGCTTGGAAAATGACGCTAAACTGCAAATCTCAAGCGAAAGTTATGAGG
AGGACCTCGGCTCCCTCCTTGCACTAATTGGAGAGAACTTCATTGAGCTGTTCTCCGCGGTC
AAGAACTTGAGTGATGGGATTCTCCTTGCTGGGATAGTTTCAGATGAATCACCCCATGCTCC
CCTCTCCACCAAAATGGTTATAAGGTTCAAGGAGCACGAGGAAGACCTCGCCGCACTCAAG
CATTTCATTAAGGCCAATTTGCCGGAGAAGTACGATGAGGTTTTTTCAGATGACTCAAAGAA
CGGCTACGCCGGGTACGTCGGTGTCGATAGTAAGGTTCGCAAAAGGAACGGGAAATTAGCA
ACCGAGGAGGAGTTCTACAAATACCTCAAGGATATTTTGAACAACGTGAAAGGCGCCGATT
ATTTCTTGGAGAAAATTAAACGGGAGGATTTACTCCGCAAACAGCGCACATTCGACAATGG
GACTATCCCATATCAGGTTCATTTAGAAGAGATGAAGGCCATTTTGCAAAACCAAGGCGAGT
ATTACCCCTTTCTTAAAGAGAACAAGGAGAAGATTCAACAGATCCTGACATTCAGAATCCCG
TACTACGTAGGACCTCTGGCACGCGGCAACCGCGACTTCGCGTGGTTGACGCGCAACTCAGA
TCAGGCGATTCGACCATGGAACTTTGAGGAGGTCGTGGATAAGGCCTCCTCCGCGGAGGACT
TCATCAACAAAATGACAAACTATGACCTCTATCTACCAGAGGAAAAGGTCCTTCCCAAGCAT
TCGCTCTTGTACGAGACATTCGCTGTCTACAACGAGCTGACCAAAGTTAAGTTTATCGCTGA
AGGACTGCGTGACTACCAATTCCTAGACTCCGGTCAAAAGAAACAGATAGTTAATCAGCTGT
TTAAAGAAAAGAGAAAAGTGACTGAGAAAGATATTATCCATTACCTCCACAACGTGGACGG
TTATGATGGGATCGAATTAAAAGGAATCGAGAAGCAGTTTAATGCTAGTCTGTCGACGTATC
ATGATCTCCTAAAAATCATTAAGGACAAGGAGTTTATGGACGATCCTAAGAACGAGGAGAT
CCTCGAGAACATCGTCCATACACTCACGATATTCGAGGACCGCGAGATGATTAAGCAGAGG
TTGGCTCAGTACGACTCTCTCTTTGATGAGAAAGTCATCAAAGCTCTAACCCGTCGCCACTAT

ACTGGCTGGGGCAAGTTATCCGCAAAGCTGATAAATGGCATCTGTGATAAGCAAACAAATA
AAACGATCCTGGACTTTCTTATTGACGACGACAAGATTAACAGAAACTTCATGCAGCTTATC
AACGACGACGGCCTTTCTTTCAAGGATATAATTCAAAAGGCGCAGGTGGTCGGCAAGATCG
ACGATGTGAAGCAAGTTGTGCAGGAGCTTCCAGGTTCTCCTGCGATTAAAAAGGGTATTTTG
CAGTCGATAAAGATTGTTGATGAGTTGGTCAAGGTCATGGGCCACGCCCCTGAGTCCATTGT
TATCGAGATGGCCAGGGAGAATCAGACTACCGCCAGGGGCAAGAAAAACTCCCAGCAGAG
ATACAAGCGCATTGAGGACGCCTTAAAAAATCTAGCGCCGGGGTTGGATTCTAATATCCTCA
AGGAAAACCCAACAGATAATATCCAGTTACAAAACGACCGCCTCTTCCTCTACTATCTTCAA
AATGGTAAGGACATGTATACCGGTGAAGCCCTCGACATAAATCAACTCAGCAATTATGATAT
AGATCATATTGTGCCGCAGGCGTTTATAAAGGACGATAGTCTTGATAACAGGGTGCTTACTA
GCAGCAAGGATAACAGGGGAAAGTCTGACAATGTGCCTTCTATAGAAGTTGTTCAGAAGCG
TAAGGCCTTCTGGCAGCAACTTCTGGACTCGAAATTGATAAGCGAGCGAAAGTTCAATAACC
TCACCAAGGCAGAGCGCGGTGGACTCGACGAACGTGACAAAGTTGGGTTCATTAAGAGGCA
GTTGGTTGAAACACGGCAGATTACAAAACATGTCGCTCAGATACTCGATGCCCGGTTCAACA
CGGAGGTAAACGAGAAGAACCAGAAGATAAGAAAGGTAAAAATTATTACTCTGAAATCCAA
CCTCGTTTCAAATTTCAGGAAAGAGTTCGGCCTGTACAAGGTCCGTGAGATAAATGACTACC
ACCACGCCCACGATGCATACCTGAATGCAGTCGTGGCGAAAGCGATCCTGAAGAAGTACCC
TAAACTTGAACCGGAGTTCGTGTATGGAGACTACCAGAAGTACGACCTTAAGAGGTACATCT
CACGCTCCAAAGATCCAAAGGAAATAGAGAAAGCAACAGAAAAGTACTTCTTTTATAGCAA
TTTGCTGAATTTTTTCAAGGAAGAAGTGCACTACGCTGATGGAACCATCATCAAGAGGGAGA
ATATAGAGTACTCTAAGGATACTGGCGAGATAGCATGGAATAAAGAAAAGGATTTCGCCAC
TGTACGCAAGGTACTGAGCTGCCCTCAGGTCAATATCGTGAAAAAGACCGAGGTTCAAACT
GGCGGCTTCTCCAAGGAGTCAATCCTCCCGAAGAGAAATTCTGACAAGTTAATCGCCCGAAA
AAAGGACTGGGACCCGAAGAAGTATGGAGGCTTCGATTCACCGACGGTGGCATATTCCGTG
CTGGTGGTTGCCAAGGTGGAGAAGGGCAAGTCAAAAAAGCTCAAGTCAGTAAAGGAACTTG
TCGGAATCACGATCATGGAGCGGTCCTCGTTCGAGAAGGACCCGGTCGACTTCCTGGAAGCA
AAAGGGTACAAGGAGGTGAGAAAAGATTTGATTATCAAGCTCCCCAAGTATTCTCTTTTCGA
GCTGGAAAATGGACGGAAGCGCATGCTCGCGTCGGCCGGCGAGCTTCAGAAAGGAAATGAA
CTGGCCCTCCCTTCAAAATATGTCAACTTCTTATATCTGGCAAGTCATTATGAAAAACTGAA
AGGGAGTCCGGAGGATAATGAACAAAAACAACTGTTTGTCGAGCAGCACAAACATTACCTC
GATGAGATTATTGAGCAAATCTCCGAGTTCTCCAAGAGGGTCATTCTCGCGGACGCAAATCT
CGACAAGGTATTATCTGCCTACAATAAACATAGAGATAAGCCAATTAGGGAACAGGCGGAG
AACATAATCCACCTCTTCACCCTCACGAACCTCGGCGCCCCAGCCGCGTTTAAGTACTTTGA
CACCACCATCGACAGGAAACGATACACTTCTACAAAAGAAGTGCTCGATGCCACGCTGATC
CATCAGAGCATTACAGGTCTTTATGAAACCCGTATAGATCTCAGCCAACTCGGAGGGGACaaa
aggccggcggccacgaaaaaggccggccaggcaaaaaagaaaaagtaagaattcgcggccgcactcgagatatctagacccagctttcttgtacaaa
gtggttgataacagcgactacaaggatgacgatgacaaggcttagagctcgaatttccccgatcgttcaaacatttggcaataaagtttcttaagattgaatc
ctgttgccggtcttgcgatgattatcatataatttctgttgaattacgttaagcatgtaataattaacatgtaatgcatgacgttatttatgagatgggtttttatgatt
agagtcccgcaattatacatttaatacgcgatagaaaacaaaatatagcgcgcaaactaggataaattatcgcgcgcggtgtcatctatgttactagatcgg
gaattc

Seq ID No. 8

AAAAAGCCTTACTCCATCGGTCTTGACATCGGTACTAACAGCGTGGGCTGGGCCGTTATTAC
TGATGATTACAAGGTGCCTGCAAAGAAGATGAAGGTGCTCGGCAATACGGACAGATCACAC
ATCAAGAAGAATCTCATTGGTGCGCTACTTTTTGACGCCGGGAACACTGCTGAGGATAGGCG
CCTGAAGAGAACGGCGCGGCGCCGCTATACAAGACGGAGGAATAGAATACTGTATTTGCAG
GAAATCTTCGCAGAAGAAATGAACAAGATTGATGAGTCCTTCTTCCACCGGCTCGACGACAG
CTTTCTCGTGCCCGAGGATAAAAGGGGCTCCAAGTATCCAATATTTGCTACTCTGCAAGAAG
AGAAGGAGTACCACAAGCAGTTCCCTACCATCTACCATCTCCGGAAACAATTGGCTGAATCT
AATGAGAAGGCAGATCTGCGATTGGTATATCTTGCTCTGGCTCACATGATCAAGTACAGAGG

GCATTTTCTCATTGACGATCCCAAATTCAAAGTACAAAACAATGATATACAGGGTCTGTTTG
AGAAATTTGTTGAAGAATACGACAATGTGCAGGAAACATCTTTGTCTAAGATAAAACTCAAT
GTCACAGAAATACTTACCGCCAAGATTCCGAAAAGTGAGAAGCAAGAGCAGCTTCTCAAGA
ACTACCCATCGGAAAAGAAGAACACGTTATTTGGGAACTTAATCGGCCTTGCGTTGGGCCTC
ACACCAAATTTCAAGACTAATTTTAGCTTGGAAAATGACGCTAAACTGCAAATCTCAAGCGA
AAGTTATGAGGAGGACCTCGGCTCCCTCCTTGCACTAATTGGAGAGAACTTCATTGAGCTGT
TCTCCGCGGTCAAGAACTTGAGTGATGGGATTCTCCTTGCTGGGATAGTTTCAGATGAATCA
CCCCATGCTCCCCTCTCCACCAAAATGGTTATAAGGTTCAAGGAGCACGAGGAAGACCTCGC
CGCACTCAAGCATTTCATTAAGGCCAATTTGCCGGAGAAGTACGATGAGGTTTTTCAGATG
ACTCAAAGAACGGCTACGCCGGGTACGTCGGTGTCGATAGTAAGGTTCGCAAAAGGAACGG
GAAATTAGCAACCGAGGAGGAGTTCTACAAATACCTCAAGGATATTTTGAACAACGTGAAA
GGCGCCGATTATTTCTTGGAGAAAATTAAACGGGAGGATTTACTCCGCAAACAGCGCACATT
CGACAATGGGACTATCCCATATCAGGTTCATTTAGAAGAGATGAAGGCCATTTTGCAAAACC
AAGGCGAGTATTACCCCTTTCTTAAAGAGAACAAGGAGAAGATTCAACAGATCCTGACATTC
AGAATCCCGTACTACGTAGGACCTCTGGCACGCGGCAACCGCGACTTCGCGTGGTTGACGCG
CAACTCAGATCAGGCGATTCGACCATGGAACTTTGAGGAGGTCGTGGATAAGGCCTCCTCCG
CGGAGGACTTCATCAACAAAATGACAAACTATGACCTCTATCTACCAGAGGAAAAGGTCCTT
CCCAAGCATTCGCTCTTGTACGAGACATTCGCTGTCTACAACGAGCTGACCAAAGTTAAGTT
TATCGCTGAAGGACTGCGTGACTACCAATTCCTAGACTCCGGTCAAAAGAAACAGATAGTTA
ATCAGCTGTTTAAAGAAAAGAGAAAAGTGACTGAGAAAGATATTATCCATTACCTCCACAA
CGTGGACGGTTATGATGGGATCGAATTAAAAGGAATCGAGAAGCAGTTTAATGCTAGTCTGT
CGACGTATCATGATCTCCTAAAAATCATTAAGGACAAGGAGTTTATGGACGATCCTAAGAAC
GAGGAGATCCTCGAGAACATCGTCCATACACTCACGATATTCGAGGACCGCGAGATGATTA
AGCAGAGGTTGGCTCAGTACGACTCTCTCTTTGATGAGAAAGTCATCAAAGCTCTAACCCGT
CGCCACTATACTGGCTGGGGCAAGTTATCCGCAAAGCTGATAAATGGCATCTGTGATAAGCA
AACAAATAAAACGATCCTGGACTTTCTTATTGACGACGACAAGATTAACAGAAACTTCATGC
AGCTTATCAACGACGACGGCCTTTCTTTCAAGGATATAATTCAAAAGGCGCAGGTGGTCGGC
AAGATCGACGATGTGAAGCAAGTTGTGCAGGAGCTTCCAGGTTCTCCTGCGATTAAAAAGG
GTATTTTGCAGTCGATAAAGATTGTTGATGAGTTGGTCAAGGTCATGGGCCACGCCCCTGAG
TCCATTGTTATCGAGATGGCCAGGGAGAATCAGACTACCGCCAGGGGCAAGAAAAACTCCC
AGCAGAGATACAAGCGCATTGAGGACGCCTTAAAAAATCTAGCGCCGGGGTTGGATTCTAA
TATCCTCAAGGAAAACCCAACAGATAATATCCAGTTACAAAACGACCGCCTCTTCCTCTACT
ATCTTCAAAATGGTAAGGACATGTATACCGGTGAAGCCCTCGACATAAATCAACTCAGCAAT
TATGATATAGATCATATTGTGCCGCAGGCGTTTATAAAGGACGATAGTCTTGATAACAGGGT
GCTTACTAGCAGCAAGGATAACAGGGGAAAGTCTGACAATGTGCCTTCTATAGAAGTTGTTC
AGAAGCGTAAGGCCTTCTGGCAGCAACTTCTGGACTCGAAATTGATAAGCGAGCGAAAGTT
CAATAACCTCACCAAGGCAGAGCGCGGTGGACTCGACGAACGTGACAAAGTTGGGTTCATT
AAGAGGCAGTTGGTTGAAACACGGCAGATTACAAAACATGTCGCTCAGATACTCGATGCCC
GGTTCAACACGGAGGTAAACGAGAAGAACCAGAAGATAAGAAAGGTAAAAATTATTACTCT
GAAATCCAACCTCGTTTCAAATTTCAGGAAAGAGTTCGGCCTGTACAAGGTCCGTGAGATAA
ATGACTACCACCACGCCCACGATGCATACCTGAATGCAGTCGTGGCGAAAGCGATCCTGAA
GAAGTACCCTAAACTTGAACCGGAGTTCGTGTATGGAGACTACCAGAAGTACGACCTTAAG
AGGTACATCTCACGCTCCAAAGATCCAAAGGAAATAGAGAAAGCAACAGAAAAGTACTTCT
TTTATAGCAATTTGCTGAATTTTTTCAAGGAAGAAGTGCACTACGCTGATGGAACCATCATC
AAGAGGGAGAATATAGAGTACTCTAAGGATACTGGCGAGATAGCATGGAATAAAGAAAAG
GATTTCGCCACTGTACGCAAGGTACTGAGCTGCCCTCAGGTCAATATCGTGAAAAAGACCGA
GGTTCAAACTGGCGGCTTCTCCAAGGAGTCAATCCTCCCGAAGAGAAATTCTGACAAGTTAA
TCGCCCGAAAAAAGGACTGGGACCCGAAGAAGTATGGAGGCTTCGATTCACCGACGGTGGC
ATATTCCGTGCTGGTGGTTGCCAAGGTGGAGAAGGGCAAGTCAAAAAAGCTCAAGTCAGTA
AAGGAACTTGTCGGAATCACGATCATGGAGCGGTCCTCGTTCGAGAAGGACCCGGTCGACTT
CCTGGAAGCAAAAGGGTACAAGGAGGTGAGAAAAGATTTGATTATCAAGCTCCCCAAGTAT

TCTCTTTTCGAGCTGGAAAATGGACGGAAGCGCATGCTCGCGTCGGCCGGCGAGCTTCAGAA
AGGAAATGAACTGGCCCTCCCTTCAAAATATGTCAACTTCTTATATCTGGCAAGTCATTATG
AAAAACTGAAAGGGAGTCCGGAGGATAATGAACAAAAACAACTGTTTGTCGAGCAGCACAA
ACATTACCTCGATGAGATTATTGAGCAAATCTCCGAGTTCTCCAAGAGGGTCATTCTCGCGG
ACGCAAATCTCGACAAGGTATTATCTGCCTACAATAAACATAGAGATAAGCCAATTAGGGA
ACAGGCGGAGAACATAATCCACCTCTTCACCCTCACGAACCTCGGCGCCCCAGCCGCGTTTA
AGTACTTTGACACCACCATCGACAGGAAACGATACACTTCTACAAAAGAAGTGCTCGATGCC
ACGCTGATCCATCAGAGCATTACAGGTCTTTATGAAACCCGTATAGATCTCAGCCAACTCGG
AGGGGAC

Seq ID No. 9

AGTGAGGTGGAGTTCTCGCACGAGTACTGGATGAGGCACGCCCTGACACTCGCAAAGCGAG
CACGCGATGAAAGGGAGGTCCCGGTGGGTGCCGTCCTCGTTCTTAATAATCGGGTAATAGGG
GAAGGCTGGAATAGGGCCATCGGACTCCACGATCCTACAGCTCATGCTGAGATCATGGCGCT
GCGCCAGGGCGGCCTGGTCATGCAGAATTATAGGCTAATTGACGCGACGCTATACGTCACGT
TCGAGCCGTGCGTTATGTGCGCAGGCGCCATGATCCACAGCAGGATTGGGAGAGTCGTCTTT
GGCGTGAGGAACTCCAAACGGGGGGCGGCCGGCTCCCTCATGAACGTTCTTAATTACCCTGG
CATGAATCATCGTGTGGAGATAACGGAGGGGATTCTTGCCGATGAGTGCGCCGCCCTGCTGT
GTGATTTCTACCGTATGCCTAGACAGGTCTTCAACGCGCAAAAAAAAGCACAGTCCTCCATC
AAC

Seq ID No. 10

atggaggcctctccggcttcaggtcctcgccatctcatggacccccatatctttacctccaatttcaataacggcattggcaggcacaagacatacctctgtt
atgaggtcgagcggctcgataacggaacctcagtgaagatggaccaacacaggggtttccttcataaccaagcgaaaaatctcctttgcggtttctatggg
cggcacgcggagctgagattcttggatttggtcccgagcctccagctggatcccgcgcaaatataccgcgtcacttggttcatatcatggagcccgtgcttt
agctgggggtgtgccggtgaggttcgcgcgttcctccaggaaaacacccatgttcgcctccggattttcgcagcgaggattttcgactatgatccactttac
aaagaggccctccaaatgcttagagatgctggggctcaagtttcaattatgacatacgacgagtttaagcattgttgggacacttttgtcgatcatcagggtt
gtcccttccagccgtgggatggtcttgatgagcattcgcaagctttgagtggcaggcttcgcgcgatacttcagaaccagggtaactcaggttctgagacc
cctggcacaagtgagtcagcaacacccgagtccaccatg

Seq ID No. 11

acaaacctcagtgacatcatagagaaggaaactggtaagcaactggttattcaagagagcattctgatgctccctgaggaagtcgaggaagttataggaa
acaagcctgagagcgatatactcgtgcatacggcgtatgacgagagcacggatgaaaatgtgatgctcttgaccagcgatgcgccagaatataaaccct
gggcactggttattcaggactctaacggggaaaataaaatcaagatgctc

Seq ID No. 12

CCGAAGAAGAAGCGGAAGGTG

Seq ID No. 13

aaaaggccggcggccacgaaaaaggccggccaggcaaaaaagaaaaag

Seq ID No. 14

aaaaagccttactccatcggtcttgccatcggtactaacagcgtgggctgggccgttattactgatgattacaaggtgcctgcaaagaagatgaaggtgct
cggcaatacggacagatcacacatcaagaagaatctcattggtgcgctactttttgacgccgggaacactgctgaggataggcgcctgaagagaacggc

gcggcgccgctatacaagacggaggaatagaatactgtatttgcaggaaatcttcgcagaagaaatgaacaagattgatgagtccttcttccaccggctc
gacgacagctttctcgtgcccgaggataaaaggggctccaagtatccaatatttgctactctgcaagaagagaaggagtaccacaagcagttccctacca
tctaccatctccggaaacAATTGGCTGAATCTAATGAGAAGGCAGATCTGCGATTGGTATATCTTGCTC
TGGCTCACATGATCAAGTACAGAGGGCATTTTCTCATTGACGATCCCAAATTCAAAGTACAA
AACAATGATATACAGGGTCTGTTTGAGAAATTTGTTGAAGAATACGACAATGTGCAGGAAA
CATCTTTGTCTAAGATAAAACTCAATGTCACAGAAATACTTACCGCCAAGATTCCGAAAAGT
GAGAAGCAAGAGCAGCTTCTCAAGAACTACCCATCGGAAAAGAAGAACACGTTATTTGGGA
ACTTAATCGGCCTTGCGTTGGGCCTCACACCAAATTTCAAGACTAATTTTAGCTTGGAAAAT
GACGCTAAACTGCAAATCTCAAGCGAAAGTTATGAGGAGGACCTCGGCTCCCTCCTTGCACT
AATTGGAGAGAACTTCATTGAGCTGTTCTCCGCGGTCAAGAACTTGAGTGATGGGATTCTCC
TTGCTGGGATAGTTTCAGATGAATCACCCCATGCTCCCCTCTCCACCAAAATGGTTATAAGG
TTCAAGGAGCACGAGGAAGACCTCGCCGCACTCAAGCATTTCATTAAGGCCAATTTGCCGGA
GAAGTACGATGAGGTTTTTCAGATGACTCAAAGAACGGCTACGCCGGGTACGTCGGTGTCG
ATAGTAAGGTTCGCAAAAGGAACGGGAAATTAGCAACCGAGGAGGAGTTCTACAAATACCT
CAAGGATATTTTGAACAACGTGAAAGGCGCCGATTATTTCTTGGAGAAAATTAAACGGGAG
GATTTACTCCGCAAACAGCGCACATTCGACAATGGGACTATCCCATATCAGGTTCATTTAGA
AGAGATGAAGGCCATTTTGCAAAACCAAGGCGAGTATTACCCCTTTCTTAAAGAGAACAAG
GAGAAGATTCAACAGATCCTGACATTCAGAATCCCGTACTACGTAGGACCTCTGGCACGCGG
CAACCGCGACTTCGCGTGGTTGACGCGCAACTCAGATCAGGCGATTCGACCATGGAACTTTG
AGGAGGTCGTGGATAAGGCCTCCTCCGCGGAGGACTTCATCAACAAAATGACAAACTATGA
CCTCTATCTACCAGAGGAAAAGGTCCTTCCCAAGCATTCGCTCTTGTACGAGACATTCGCTG
TCTACAACGAGCTGACCAAAGTTAAGTTTATCGCTGAAGGACTGCGTGACTACCAATTCCTA
GACTCCGGTCAAAAGAAACAGATAGTTAATCAGCTGTTTAAAGAAAAGAGAAAAGTGACTG
AGAAAGATATTATCCATTACCTCCACAACGTGGACGGTTATGATGGGATCGAATTAAAAGG
AATCGAGAAGCAGTTTAATGCTAGTCTGTCGACGTATCATGATCTCCTAAAAATCATTAAGG
ACAAGGAGTTTATGGACGATCCTAAGAACGAGGAGATCCTCGAGAACATCGTCCATACACT
CACGATATTCGAGGACCGCGAGATGATTAAGCAGAGGTTGGCTCAGTACGACTCTCTCTTTG
ATGAGAAAGTCATCAAAGCTCTAACCCGTCGCCACTATACTGGCTGGGGCAAGTTATCCGCA
AAGCTGATAAATGGCATCTGTGATAAGCAAACAAATAAAACGATCCTGGACTTTCTTATTGA
CGACGACAAGATTAACAGAAACTTCATGCAGCTTATCAACGACGACGGCCTTTCTTTCAAGG
ATATAATTCAAAAGGCGCAGGTGGTCGGCAAGATCGACGATGTGAAGCAAGTTGTGCAGGA
GCTTCCAGGTTCTCCTGCGATTAAAAAGGGTATTTTGCAGTCGATAAAGATTGTTGATGAGT
TGGTCAAGGTCATGGGCCACGCCCCTGAGTCCATTGTTATCGAGATGGCCAGGGAGAATCAG
ACTACCGCCAGGGGCAAGAAAAACTCCCAGCAGAGATACAAGCGCATTGAGGACGCCTTAA
AAAATCTAGCGCCGGGGTTGGATTCTAATATCCTCAAGGAAAACCCAACAGATAATATCCA
GTTACAAAACGACCGCCTCTTCCTCTACTATCTTCAAAATGGTAAGGACATGTATACCGGTG
AAGCCCTCGACATAAATCAACTCAGCAATTATGATATAGATCATATTGTGCCGCAGGCGTTT
ATAAAGGACGATAGTCTTGATAACAGGGTGCTTACTAGCAGCAAGGATAACAGGGGAAAGT
CTGACAATGTGCCTTCTATAGAAGTTGTTCAGAAGCGTAAGGCCTTCTGGCAGCAACTTCTG
GACTCGAAATTGATAAGCGAGCGAAAGTTCAATAACCTCACCAAGGCAGAGCGCGGTGGAC
TCGACGAACGTGACAAAGTTGGGTTCATTAAGAGGCAGTTGGTTGAAACACGGCAGATTAC
AAAACATGTCGCTCAGATACTCGATGCCCGGTTCAACACGGAGGTAAACGAGAAGAACCAG
AAGATAAGAAAGGTAAAAATTATTACTCTGAAATCCAACCTCGTTTCAAATTTCAGGAAAGA
GTTCGGCCTGTACAAGGTCCGTGAGATAAATGACTACCACCACGCCCACGATGCATACCTGA
ATGCAGTCGTGGCGAAAGCGATCCTGAAGAAGTACCCTAAACTTGAACCGGAGTTCGTGTAT
GGAGACTACCAGAAGTACGACCTTAAGAGGTACATCTCACGCTCCAAAGATCCAAAGGAAA
TAGAGAAAGCAACAGAAAAGTACTTCTTTTATAGCAATTTGCTGAATTTTTTCAAGGAAGAA
GTGCACTACGCTGATGGAACCATCATCAAGAGGGAGAATATAGAGTACTCTAAGGATACTG
GCGAGATAGCATGGAATAAAGAAAAGGATTTCGCCACTGTACGCAAGGTACTGAGCTGCCC
TCAGGTCAATATCGTGAAAAAGACCGAGGTTCAAACTGGCGGCTTCTCCAAGGAGTCAATCC
TCCCGAAGAGAAATTCTGACAAGTTAATCGCCCGAAAAAAGGACTGGGACCCGAAGAAGTA

TGGAGGCTTCGATTCACCGACGGTGGCATATTCCGTGCTGGTGGTTGCCAAGGTGGAGAAGG
GCAAGTCAAAAAAGCTCAAGTCAGTAAAGGAACTTGTCGGAATCACGATCATGGAGCGGTC
CTCGTTCGAGAAGGACCCGGTCGACTTCCTGGAAGCAAAAGGGTACAAGGAGGTGAGAAAA
GATTTGATTATCAAGCTCCCCAAGTATTCTCTTTTCGAGCTGGAAAATGGACGGAAGCGCAT
GCTCGCGTCGGCCGGCGAGCTTCAGAAAGGAAATGAACTGGCCCTCCCTTCAAAATATGTCA
ACTTCTTATATCTGGCAAGTCATTATGAAAAACTGAAAGGGAGTCCGGAGGATAATGAACA
AAAACAACTGTTTGTCGAGCAGCACAAACATTACCTCGATGAGATTATTGAGCAAATCTCCG
AGTTCTCCAAGAGGGTCATTCTCGCGGACGCAAATCTCGACAAGGTATTATCTGCCTACAAT
AAACATAGAGATAAGCCAATTAGGGAACAGGCGGAGAACATAATCCACCTCTTCACCCTCA
CGAACCTCGGCGCCCCAGCCGCGTTTAAGTACTTTGACACCACCATCGACAGGAAACGATAC
ACTTCTACAAAAGAAGTGCTCGATGCCACGCTGATCCATCAGAGCATTACAGGTCTTTATGA
AACCCGTATAGATCTCAGCCAACTCGGAGGGGAC

Seq ID No. 15

AAAAAGCCTTACTCCATCGGTCTTGACATCGGTACTAACAGCGTGGGCTGGGCCGTTATTAC
TGATGATTACAAGGTGCCTGCAAAGAAGATGAAGGTGCTCGGCAATACGGACAGATCACAC
ATCAAGAAGAATCTCATTGGTGCGCTACTTTTTGACGCCGGGAACACTGCTGAGGATAGGCG
CCTGAAGAGAACGGCGCGGCGCCGCTATACAAGACGGAGGAATAGAATACTGTATTTGCAG
GAAATCTTCGCAGAAGAAATGAACAAGATTGATGAGTCCTTCTTCCACCGGCTCGACGACAG
CTTTCTCGTGCCCGAGGATAAAAGGGGCTCCAAGTATCCAATATTTGCTACTCTGCAAGAAG
AGAAGGAGTACCACAAGCAGTTCCCTACCATCTACCATCTCCGGAAACAATTGGCTGAATCT
AATGAGAAGGCAGATCTGCGATTGGTATATCTTGCTCTGGCTCACATGATCAAGTACAGAGG
GCATTTTCTCATTGACGATCCCAAATTCAAAGTACAAAACAATGATATACAGGGTCTGTTTG
AGAAATTTGTTGAAGAATACGACAATGTGCAGGAAACATCTTTGTCTAAGATAAAACTCAAT
GTCACAGAAATACTTACCGCCAAGATTCCGAAAAGTGAGAAGCAAGAGCAGCTTCTCAAGA
ACTACCCATCGGAAAAGAAGAACACGTTATTTGGGAACTTAATCGGCCTTGCGTTGGGCCTC
ACACCAAATTTCAAGACTAATTTTAGCTTGGAAAATGACGCTAAACTGCAAATCTCAAGCGA
AAGTTATGAGGAGGACCTCGGCTCCCTCCTTGCACTAATTGGAGAGAACTTCATTGAGCTGT
TCTCCGCGGTCAAGAACTTGAGTGATGGGATTCTCCTTGCTGGGATAGTTTCAGATGAATCA
CCCCATGCTCCCCTCTCCACCAAAATGGTTATAAGGTTCAAGGAGCACGAGGAAGACCTCGC
CGCACTCAAGCATTTCATTAAGGCCAATTTGCCGGAGAAGTACGATGAGGTTTTTTCAGATG
ACTCAAAGAACGGCTACGCCGGGTACGTCGGTGTCGATAGTAAGGTTCGCAAAAGGAACGG
GAAATTAGCAACCGAGGAGGAGTTCTACAAATACCTCAAGGATATTTTGAACAACGTGAAA
GGCGCCGATTATTTCTTGGAGAAAATTAAACGGGAGGATTTACTCCGCAAACAGCGCACATT
CGACAATGGGACTATCCCATATCAGGTTCATTTAGAAGAGATGAAGGCCATTTTGCAAAACC
AAGGCGAGTATTACCCCTTTCTTAAAGAGAACAAGGAGAAGATTCAACAGATCCTGACATTC
AGAATCCCGTACTACGTAGGACCTCTGGCACGCGGCAACCGCGACTTCGCGTGGTTGACGCG
CAACTCAGATCAGGCGATTCGACCATGGAACTTTGAGGAGGTCGTGGATAAGGCCTCCTCCG
CGGAGGACTTCATCAACAAAATGACAAACTATGACCTCTATCTACCAGAGGAAAAGGTCCTT
CCCAAGCATTCGCTCTTGTACGAGACATTCGCTGTCTACAACGAGCTGACCAAAGTTAAGTT
TATCGCTGAAGGACTGCGTGACTACCAATTCCTAGACTCCGGTCAAAAGAAACAGATAGTTA
ATCAGCTGTTTAAAGAAAAGAGAAAAGTGACTGAGAAAGATATTATCCATTACCTCCACAA
CGTGGACGGTTATGATGGGATCGAATTAAAAGGAATCGAGAAGCAGTTTAATGCTAGTCTGT
CGACGTATCATGATCTCCTAAAAATCATTAAGGACAAGGAGTTTATGGACGATCCTAAGAAC
GAGGAGATCCTCGAGAACATCGTCCATACACTCACGATATTCGAGGACCGCGAGATGATTA
AGCAGAGGTTGGCTCAGTACGACTCTCTCTTTGATGAGAAAGTCATCAAAGCTCTAACCCGT
CGCCACTATACTGGCTGGGGCAAGTTATCCGCAAAGCTGATAAATGGCATCTGTGATAAGCA
AACAAATAAAACGATCCTGGACTTTCTTATTGACGACGACAAGATTAACAGAAACTTCATGC
AGCTTATCAACGACGACGGCCTTTCTTTCAAGGATATAATTCAAAAGGCGCAGGTGGTCGGC
AAGATCGACGATGTGAAGCAAGTTGTGCAGGAGCTTCCAGGTTCTCCTGCGATTAAAAAGG

GTATTTTGCAGTCGATAAAGATTGTTGATGAGTTGGTCAAGGTCATGGGCCACGCCCCTGAG
TCCATTGTTATCGAGATGGCCAGGGAGAATCAGACTACCGCCAGGGGCAAGAAAAACTCCC
AGCAGAGATACAAGCGCATTGAGGACGCCTTAAAAAATCTAGCGCCGGGGTTGGATTCTAA
TATCCTCAAGGAAAACCCAACAGATAATATCCAGTTACAAAACGACCGCCTCTTCCTCTACT
ATCTTCAAAATGGTAAGGACATGTATAccggtgaagccctcgacataaatcaactcagcaattatgatatagatgccattgtgcc
gcaggcgtttataaaggacgatagtcttgataacagggtgcttactagcagcaaggataacaggggaaagtctgacaatgtgccttctatagaagttgttca
gaagcgtaaggccttctggcagcaacttctggactcgaaattgataagcgagcgaaagttcaataacctcaccaaggcagagcgcggtggactcgacg
aacgtgacaaagttgggttcattaagaggcagttggttgaaacacggcagattacaaaacatgtcgctcagatactcgatgcccggttcaacacggaggt
aaacgagaagaaccagaagataagaaaggtaaaaattattactctgaaatccaacctcgtttcaaatttcaggaaagagttcggcctGTACAAGG
TCCGTGAGATAAATGACTACCACCACGCCCACGATGCATACCTGAATGCAGTCGTGGCGAA
AGCGATCCTGAAGAAGTACCCTAAACTTGAACCGGAGTTCGTGTATGGAGACTACCAGAAG
TACGACCTTAAGAGGTACATCTCACGCTCCAAAGATCCAAAGGAAATAGAGAAAGCAACAG
AAAAGTACTTCTTTTATAGCAATTTGCTGAATTTTTTCAAGGAAGAAGTGCACTACGCTGAT
GGAACCATCATCAAGAGGGAGAATATAGAGTACTCTAAGGATACTGGCGAGATAGCATGGA
ATAAAGAAAAGGATTTCGCCACTGTACGCAAGGTACTGAGCTGCCCTCAGGTCAATATCGTG
AAAAAGACCGAGGTTCAAACTGGCGGCTTCTCCAAGGAGTCAATCCTCCCGAAGAGAAATT
CTGACAAGTTAATCGCCCGAAAAAAGGACTGGGACCCGAAGAAGTATGGAGGCTTCGATTC
ACCGACGGTGGCATATTCCGTGCTGGTGGTTGCCAAGGTGGAGAAGGGCAAGTCAAAAAAG
CTCAAGTCAGTAAAGGAACTTGTCGGAATCACGATCATGGAGCGGTCCTCGTTCGAGAAGG
ACCCGGTCGACTTCCTGGAAGCAAAAGGGTACAAGGAGGTGAGAAAAGATTTGATTATCAA
GCTCCCCAAGTATTCTCTTTTCGAGCTGGAAAATGGACGGAAGCGCATGctcgcgtcggccggcgagctt
cagaaaggaaatgaactggccctcccttcaaaatatgtcaacttcttatatctggcaagtcattatgaaaaactgaaagggagtccggaggataatgaaca
aaaacaactgtttgtcgagcagcacaaacattacctcgatgagattattgagcaaatctccgagttctccaagagggtcattctcgcggacgcaaatctcga
caaggtattatctgcctacaataaacatagagataagccaattagggaacaggcggagaacataatccacctcttcaccctcacgaacctcggcgcccca
gccgcgtttaagtactttgacaccaccatcgacaggaaacgatacacttctacaaaagaagtgctcgatgccacgctgatccatcagagcattacaggtct
ttatgaaacccgtatagatctcagccaactcggaggggac

Seq ID No. 16

tcaggcggctcatctggcgggtcaaagcgcacagccgacggctctgagttcgagagccctaagaagaagcgcaaggtgtcaggcggctcttcaggcg
gcagc

Seq ID No. 17

Accctgaacattgaggacgagtaccggctgcacgagacgagcaaggagccagacgtttcgctcggcagcacttggctctctgacttcccacaggcttg
ggccgagactggcggcatgggcctggccgtgcgccaggctccactgatcatccctctgaaggcgacctccaccccggtttctattaagcagtacccgat
gagccaggaggccaggctggggatcaagccacacattcagcggctgctggaccagggcatcctggtgccatgccagtccccgtggaatactccgctc
ctgccggtgaagaagcctgggacaaacgactacaggccggttcaggatctcagggaggtgaacaagcgcgtggaggacatccatccgacagtgccg
aacccgtacaatctgctgtcgggcctgcctccgagccaccagtggtacaccgtcctggacctcaaggacgctttcttctgcctgcggctgcacccgacgt
ctcagccgctgttcgcgttcgagtggcgcgacccagagatgggcatttccggccagctgacctggacacgcctaccccagggcttcaagaactccccg
actctcttcaacgaggctctccaccgggatctcgcggacttcaggattcagcatcccgatctgatcctgctccagtatgttgacgacctcctcctggccgcg
acgtcggagctggactgccagcagggcacccgggcgctgctgcagacactgggcaatctggggtaccgcgcctctgcgaagaaggcgcagatctgc
cagaagcaagtgaagtacctgggctacctcctgaaggagggccagcgctggctcactgaggcgaggaaggagactgttatgggccagcccactccaa
agactccgaggcagctcagggagttcctcggcaaggctgggttctgccgcctgttcatccctgggttcgctgagatggctgcgccgctctacccgctgac
taagccggggacactgttcaactgggggccagaccagcagaaggcgtaccaggagattaagcaggcgctgctgacggccccagcgctcggcctacc
agacctgacgaagccgttcgagctgttcgttgacgagaagcaggggtacgcgaagggcgtgctgacacagaagctggggccttggcgccgcccggt
cgcgtacctgtcgaagaagctggacccagtcgctgctgggtggcctccatgcctccggatggtcgctgctattgcggttctgaccaaggatgcggggaa
gctcacaatggggcagcctctcgtgatcctggctccacatgcggtggaggcgctggtgaagcagccaccggaccggtggctgtcgaacgctcggatg
acacactaccaggcgctcctcctcgatacagaccgggttcagttcgggcctgtggttgctctgaacccagccacactgctgccactccctgaggagggc
ctccagcacaattgcctcgacatcctggctgaggcgcacggcacccgccctgatctcaccgaccagcctctgccagatgctgaccacacctggtacacg

gatgggtcctcgctgctgcaggagggccagaggaaggcgggcgccgccgtcaccacagagacagaggttatttgggccaaggccctaccggctggc
accagcgcccagcgcgctgagctgatcgcgctgactcaggcgctgaagatggccgaggggaagaagctcaatgtttacaccgactcgcggtacgcgt
tcgctacagctcacattcatggggagatctaccgccggcgcgggtggctgacttcggagggcaaggagattaagaataaggacgagatcctggccctg
ctcaaggcgctgttcctgccgaagcgcctctcaatcattcactgcccgggccaccagaagggccattcggccgaggctaggggcaatcggatggctga
ccaggcggcgcggaaggcggctatcaccgagactcccgatacatctaccctcctgatcgagaactcgagccca

Seq ID No. 18

GTTTTATGGCTGGAAATAGCAAGTTAAAATAAGGctagtccgttatcaacttgaaaaagtggcACCGAGTCGG
TGC

Seq ID No. 19

GTTTTATGGCTGATAAATTTCTTTGAATTTCTCCTTGATTATTTGTTATAAAAGTTATAAAATA
ATCTTGTTGGAACCATTCAAAACAGCATAGCAAGTTAAAATAAGGCTAGTCCgttatcaacttgaaaaa
gtggcACCGAGTCGGTGCTTTTTTT

#### Polypeptides

##### Rice codon optimized PAiD

KKPYSIGLDIGTNSVGWAVITDDYKVPAKKMKVLGNTDRSHIKKNLIGALLFDAGNTAEDRRLK
RTARRRYTRRRNRILYLQEIFAEEMNKIDESFFHRLDDSFLVPEDKRGSKYPIFATLQEEKEYHKQ
FPTIYHLRKQLAESNEKADLRLVYLALAHMIKYRGHFLIDDPKFKVQNNDIQGLFEKFVEEYDNV
QETSLSKIKLNVTEILTAKIPKSEKQEQLLKNYPSEKKNTLFGNLIGLALGLTPNFKTNFSLENDAK
LQISSESYEEDLGSLLALIGENFIELFSAVKNLSDGILLAGIVSDESPHAPLSTKMVIRFKEHEEDLA
ALKHFIKANLPEKYDEVFSDDSKNGYAGYVGVDSKVRKRNGKLATEEEFYKYLKDILNNVKGA
DYFLEKIKREDLLRKQRTFDNGTIPYQVHLEEMKAILQNQGEYYPFLKENKEKIQQILTFRIPYYV
GPLARGNRDFAWLTRNSDQAIRPWNFEEVVDKASSAEDFINKMTNYDLYLPEEKVLPKHSLLYE
TFAVYNELTKVKFIAEGLRDYQFLDSGQKKQIVNQLFKEKRKVTEKDIIHYLHNVDGYDGIELKG
IEKQFNASLSTYHDLLKIIKDKEFMDDPKNEEILENIVHTLTIFEDREMIKQRLAQYDSLFDEKVIK
ALTRRHYTGWGKLSAKLINGICDKQTNKTILDFLIDDDKINRNFMQLINDDGLSFKDIIQKAQVV
GKIDDVKQVVQELPGSPAIKKGILQSIKIVDELVKVMGHAPESIVIEMARENQTTARGKKNSQQR
YKRIEDALKNLAPGLDSNILKENPTDNIQLQNDRLFLYYLQNGKDMYTGEALDINQLSNYDIDHI
VPQAFIKDDSLDNRVLTSSKDNRGKSDNVPSIEVVQKRKAFWQQLLDSKLISERKFNNLTKAERG
GLDERDKVGFIKRQLVETRQITKHVAQILDARFNTEVNEKNQKIRKVKIITLKSNLVSNFRKEFGL
YKVREINDYHHAHDAYLNAVVAKAILKKYPKLEPEFVYGDYQKYDLKRYISRSKDPKEIEKATE
KYFFYSNLLNFFKEEVHYADGTIIKRENIEYSKDTGEIAWNKEKDFATVRKVLSCPQVNIVKKTE
VQTGGFSKESILPKRNSDKLIARKKDWDPKKYGGFDSPTVAYSVLVVAKVEKGKSKKLKSVKEL
VGITIMERSSFEKDPVDFLEAKGYKEVRKDLIIKLPKYSLFELENGRKRMLASAGELQKGNELAL
PSKYVNFLYLASHYEKLKGSPEDNEQKQLFVEQHKHYLDEIIEQISEFSKRVILADANLDKVLSA
YNKHRDKPIREQAENIIHLFTLTNLGAPAAFKYFDTTIDRKRYTSTKEVLDATLIHQSITGLYETRI
DLSQLGGD

##### Rice codon optimized PAiD-D10A

KKPYSIGLAIGTNSVGWAVITDDYKVPAKKMKVLGNTDRSHIKKNLIGALLFDAGNTAEDRRLK
RTARRRYTRRRNRILYLQEIFAEEMNKIDESFFHRLDDSFLVPEDKRGSKYPIFATLQEEKEYHKQ
FPTIYHLRKQLAESNEKADLRLVYLALAHMIKYRGHFLIDDPKFKVQNNDIQGLFEKFVEEYDNV
QETSLSKIKLNVTEILTAKIPKSEKQEQLLKNYPSEKKNTLFGNLIGLALGLTPNFKTNFSLENDAK
LQISSESYEEDLGSLLALIGENFIELFSAVKNLSDGILLAGIVSDESPHAPLSTKMVIRFKEHEEDLA
ALKHFIKANLPEKYDEVFSDDSKNGYAGYVGVDSKVRKRNGKLATEEEFYKYLKDILNNVKGA
DYFLEKIKREDLLRKQRTFDNGTIPYQVHLEEMKAILQNQGEYYPFLKENKEKIQQILTFRIPYYV
GPLARGNRDFAWLTRNSDQAIRPWNFEEVVDKASSAEDFINKMTNYDLYLPEEKVLPKHSLLYE
TFAVYNELTKVKFIAEGLRDYQFLDSGQKKQIVNQLFKEKRKVTEKDIIHYLHNVDGYDGIELKG
IEKQFNASLSTYHDLLKIIKDKEFMDDPKNEEILENIVHTLTIFEDREMIKQRLAQYDSLFDEKVIK
ALTRRHYTGWGKLSAKLINGICDKQTNKTILDFLIDDDKINRNFMQLINDDGLSFKDIIQKAQVV
GKIDDVKQVVQELPGSPAIKKGILQSIKIVDELVKVMGHAPESIVIEMARENQTTARGKKNSQQR
YKRIEDALKNLAPGLDSNILKENPTDNIQLQNDRLFLYYLQNGKDMYTGEALDINQLSNYDIDHI
VPQAFIKDDSLDNRVLTSSKDNRGKSDNVPSIEVVQKRKAFWQQLLDSKLISERKFNNLTKAERG
GLDERDKVGFIKRQLVETRQITKHVAQILDARFNTEVNEKNQKIRKVKIITLKSNLVSNFRKEFGL
YKVREINDYHHAHDAYLNAVVAKAILKKYPKLEPEFVYGDYQKYDLKRYISRSKDPKEIEKATE
KYFFYSNLLNFFKEEVHYADGTIIKRENIEYSKDTGEIAWNKEKDFATVRKVLSCPQVNIVKKTE
VQTGGFSKESILPKRNSDKLIARKKDWDPKKYGGFDSPTVAYSVLVVAKVEKGKSKKLKSVKEL
VGITIMERSSFEKDPVDFLEAKGYKEVRKDLIIKLPKYSLFELENGRKRMLASAGELQKGNELAL
PSKYVNFLYLASHYEKLKGSPEDNEQKQLFVEQHKHYLDEIIEQISEFSKRVILADANLDKVLSA
YNKHRDKPIREQAENIIHLFTLTNLGAPAAFKYFDTTIDRKRYTSTKEVLDATLIHQSITGLYETRI
DLSQLGGD

**Rice codon optimized PAiD-H850A**

KKPYSIGLDIGTNSVGWAVITDDYKVPAKKMKVLGNTDRSHIKKNLIGALLFDAGNTAEDRRLK
RTARRRYTRRRNRILYLQEIFAEEMNKIDESFFHRLDDSFLVPEDKRGSKYPIFATLQEEKEYHKQ
FPTIYHLRKQLAESNEKADLRLVYLALAHMIKYRGHFLIDDPKFKVQNNDIQGLFEKFVEEYDNV
QETSLSKIKLNVTEILTAKIPKSEKQEQLLKNYPSEKKNTLFGNLIGLALGLTPNFKTNFSLENDAK
LQISSESYEEDLGSLLALIGENFIELFSAVKNLSDGILLAGIVSDESPHAPLSTKMVIRFKEHEEDLA
ALKHFIKANLPEKYDEVFSDDSKNGYAGYVGVDSKVRKRNGKLATEEEFYKYLKDILNNVKGA
DYFLEKIKREDLLRKQRTFDNGTIPYQVHLEEMKAILQNQGEYYPFLKENKEKIQQILTFRIPYYV
GPLARGNRDFAWLTRNSDQAIRPWNFEEVVDKASSAEDFINKMTNYDLYLPEEKVLPKHSLLYE
TFAVYNELTKVKFIAEGLRDYQFLDSGQKKQIVNQLFKEKRKVTEKDIIHYLHNVDGYDGIELKG
IEKQFNASLSTYHDLLKIIKDKEFMDDPKNEEILENIVHTLTIFEDREMIKQRLAQYDSLFDEKVIK
ALTRRHYTGWGKLSAKLINGICDKQTNKTILDFLIDDDKINRNFMQLINDDGLSFKDIIQKAQVV
GKIDDVKQVVQELPGSPAIKKGILQSIKIVDELVKVMGHAPESIVIEMARENQTTARGKKNSQQR
YKRIEDALKNLAPGLDSNILKENPTDNIQLQNDRLFLYYLQNGKDMYTGEALDINQLSNYDIDAI
VPQAFIKDDSLDNRVLTSSKDNRGKSDNVPSIEVVQKRKAFWQQLLDSKLISERKFNNLTKAERG
GLDERDKVGFIKRQLVETRQITKHVAQILDARFNTEVNEKNQKIRKVKIITLKSNLVSNFRKEFGL
YKVREINDYHHAHDAYLNAVVAKAILKKYPKLEPEFVYGDYQKYDLKRYISRSKDPKEIEKATE
KYFFYSNLLNFFKEEVHYADGTIIKRENIEYSKDTGEIAWNKEKDFATVRKVLSCPQVNIVKKTE
VQTGGFSKESILPKRNSDKLIARKKDWDPKKYGGFDSPTVAYSVLVVAKVEKGKSKKLKSVKEL
VGITIMERSSFEKDPVDFLEAKGYKEVRKDLIIKLPKYSLFELENGRKRMLASAGELQKGNELAL
PSKYVNFLYLASHYEKLKGSPEDNEQKQLFVEQHKHYLDEIIEQISEFSKRVILADANLDKVLSA
YNKHRDKPIREQAENIIHLFTLTNLGAPAAFKYFDTTIDRKRYTSTKEVLDATLIHQSITGLYETRI
DLSQLGGD

**TadA8e**

SEVEFSHEYWMRHALTLAKRARDEREVPVGAVLVLNNRVIGEGWNRAIGLHDPTAHAEIMALR
QGGLVMQNYRLIDATLYVTFEPCVMCAGAMIHSRIGRVVFGVRNSKRGAAGSLMNVLNYPGM
NHRVEITEGILADECAALLCDFYRMPRQVFNAQKKAQSSIN

**A3A_Y130F**

MEASPASGPRHLMDPHIFTSNFNNGIGRHKTYLCYEVERLDNGTSVKMDQHRGFLHNQAKNLL
CGFYGRHAELRFLDLVPSLQLDPAQIYRVTWFISWSPCFSWGCAGEVRAFLQENTHVRLRIFAAR
IFDYDPLYKEALQMLRDAGAQVSIMTYDEFKHCWDTFVDHQGCPFQPWDGLDEHSQALSGRLR
AILQNQGNSGSETPGTSESATPESTM

**UGI**

TNLSDIIEKETGKQLVIQESILMLPEEVEEVIGNKPESDILVHTAYDESTDENVMLLTSDAPEYKP
WALVIQDSNGENKIKML

**Linker for prime editing**

SGGSSGGSKRTADGSEFESPKKKRKVSGGSSGGS

**M-MLV-RT**

TLNIEDEYRLHETSKEPDVSLGSTWLSDFPQAWAETGGMGLAVRQAPLIIPLKATSTPVSIKQYP
MSQEARLGIKPHIQRLLDQGILVPCQSPWNTPLLPVKKPGTNDYRPVQDLREVNKRVEDIHPTVP

NPYNLLSGLPPSHQWYTVLDLKDAFFCLRLHPTSQPLFAFEWRDPEMGISGQLTWTRLPQGFKN
SPTLFNEALHRDLADFRIQHPDLILLQYVDDLLLAATSELDCQQGTRALLQTLGNLGYRASAKKA
QICQKQVKYLGYLLKEGQRWLTEARKETVMGQPTPKTPRQLREFLGKAGFCRLFIPGFAEMAAP
LYPLTKPGTLFNWGPDQQKAYQEIKQALLTAPALGLPDLTKPFELFVDEKQGYAKGVLTQKLGP
WRRPVAYLSKKLDPVAAGWPPCLRMVAAIAVLTKDAGKLTMGQPLVILAPHAVEALVKQPPDR
WLSNARMTHYQALLLDTDRVQFGPVVALNPATLLPLPEEGLQHNCLDILAEAHGTRPDLTDQPL
PDADHTWYTDGSSLLQEGQRKAGAAVTTETEVIWAKALPAGTSAQRAELIALTQALKMAEGKK
LNVYTDSRYAFATAHIHGEIYRRRGWLTSEGKEIKNKDEILALLKALFLPKRLSIIHCPGHQKGHS
AEARGNRMADQAARKAAITETPDTSTLLIENSSP

**Bipartite SV40 NLS**

KRTADGSEFESPKKKRKV

**Nucleoplasmin NLS**

KRPAATKKAGQAKKKK

#### Vector maps

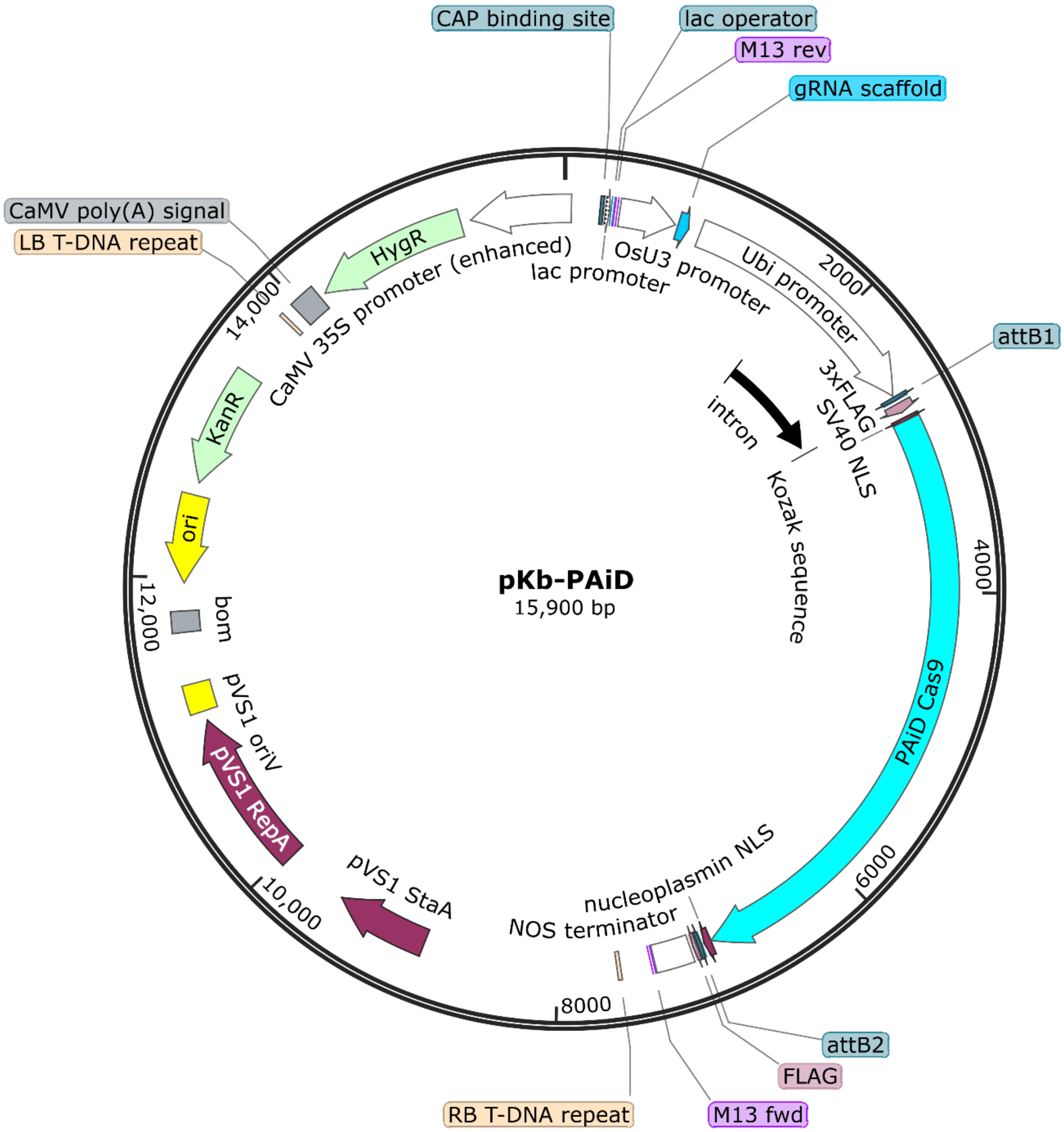


**Supplementary figure 1:** Schematic diagram showing different components of the pKb-PAiD mother vector, used for knock-out experiments in rice. The sgRNA component is driven under OsU3 promoter (An RNA PolIII promoter, terminated by PolyT) and the endonuclease is driven under OsUbi10 promoter (An RNA PolII promoter, terminated by Nos terminator).

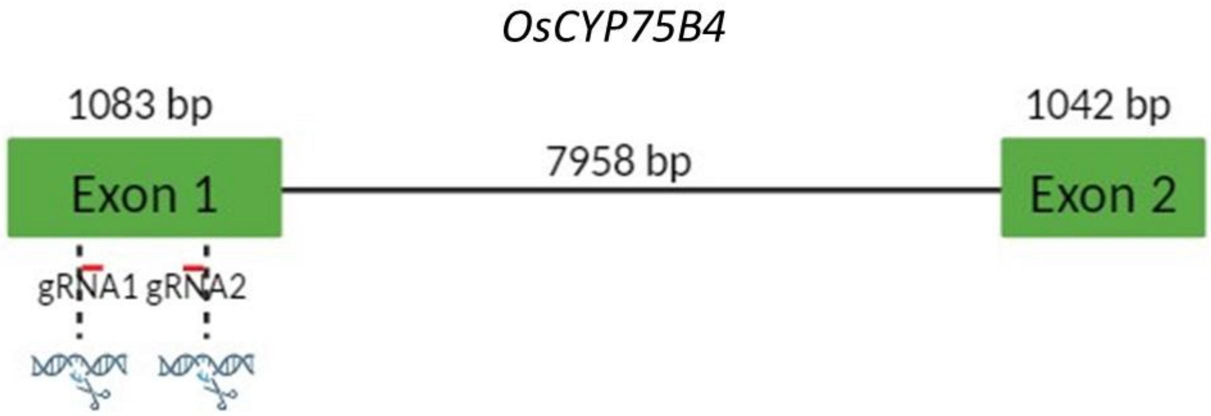


**Supplementary figure 2:** Schematic diagram showing the architecture of *OsCYP75B4* gene comprising two exons, intervened by one intron. Two guide RNAs used in the study, had been chosen from exon1 flanking 573bp region. This target gene was also used to generate mutant plant lines for this study.

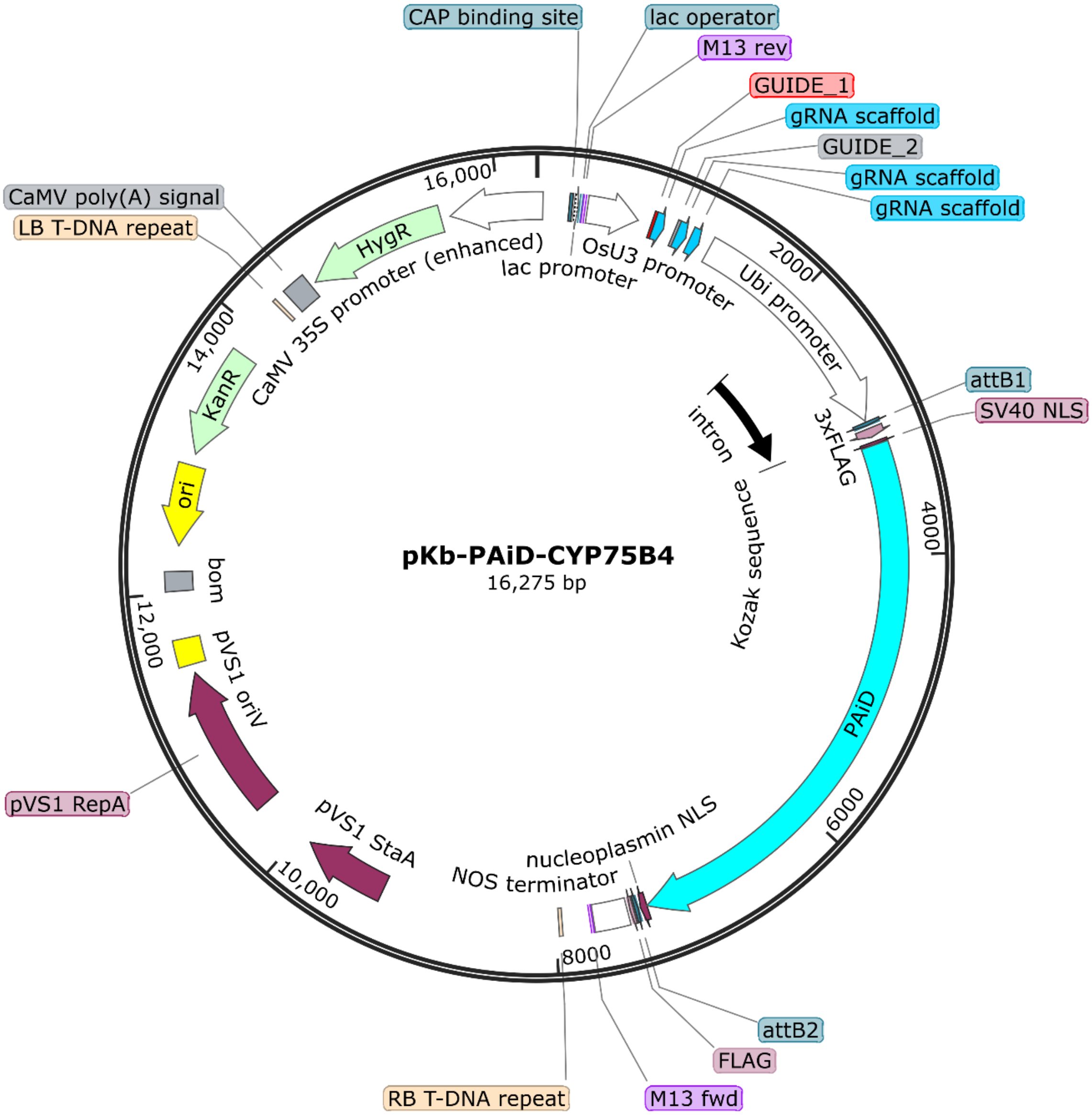


**Supplementary figure 3:** Schematic diagram showing different components of the pKb-PAiD-CYP75B4, used for multiplexed knock-out of *OsCYP75B4* gene in rice. The PTG (multiplexed sgRNA cassette) component is driven under OsU3 promoter (an RNA Pol III promoter, terminated by PolyT) and the endonuclease is driven under OsUbi10 promoter (an RNA Pol II promoter, terminated by Nos terminator).

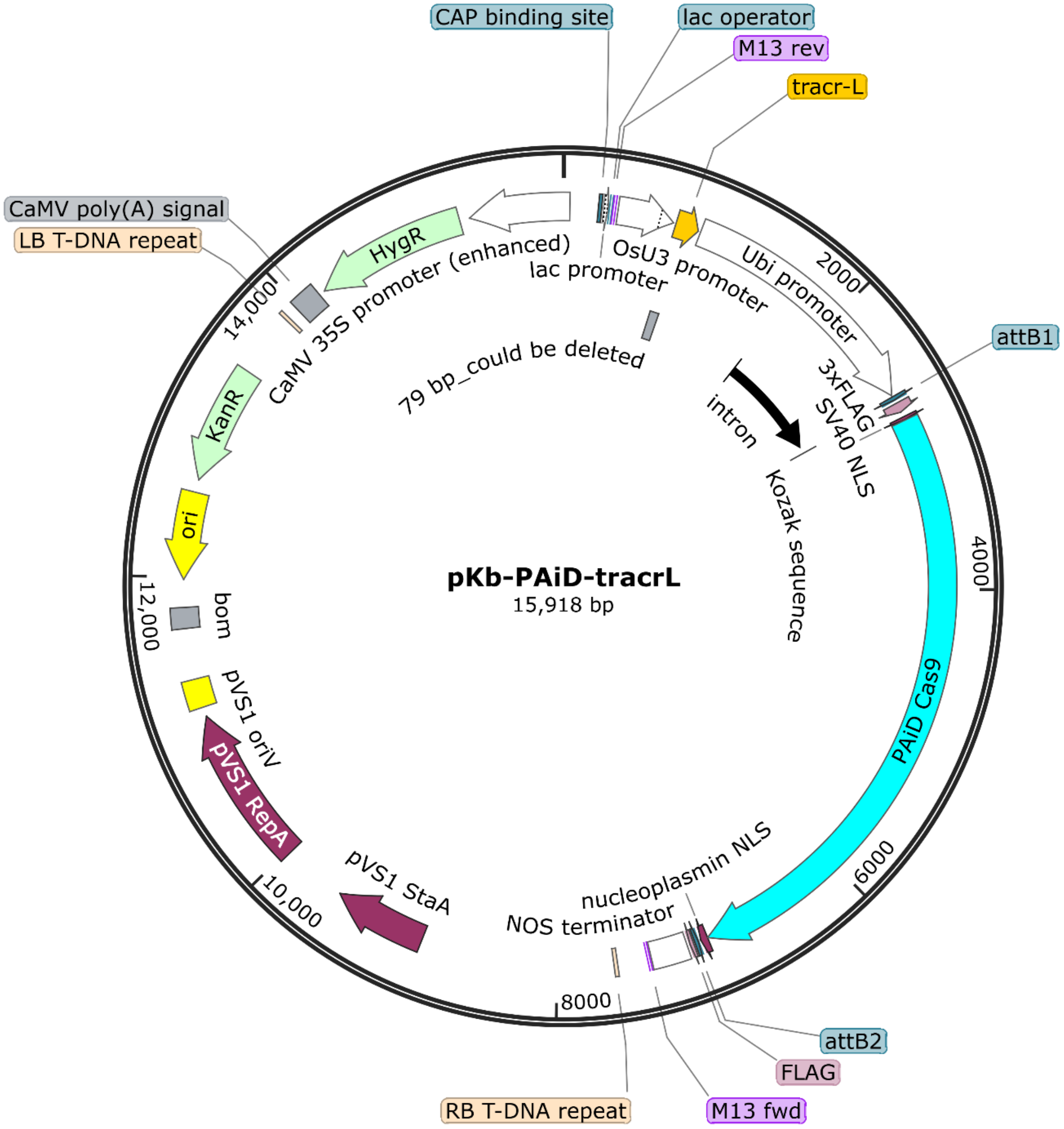


**Supplementary figure 4:** Schematic diagram showing different components of the pKb-PAiD-tracrL mother vector, used for alternative tracrRNA experiments in rice. The sgRNA component is driven under OsU3 promoter (An RNA PolIII promoter, terminated by PolyT) and the endonuclease is driven under OsUbi10 promoter (An RNA PolII promoter, terminated by Nos terminator). The normal tracrRNA is replaced by long tracrRNA.

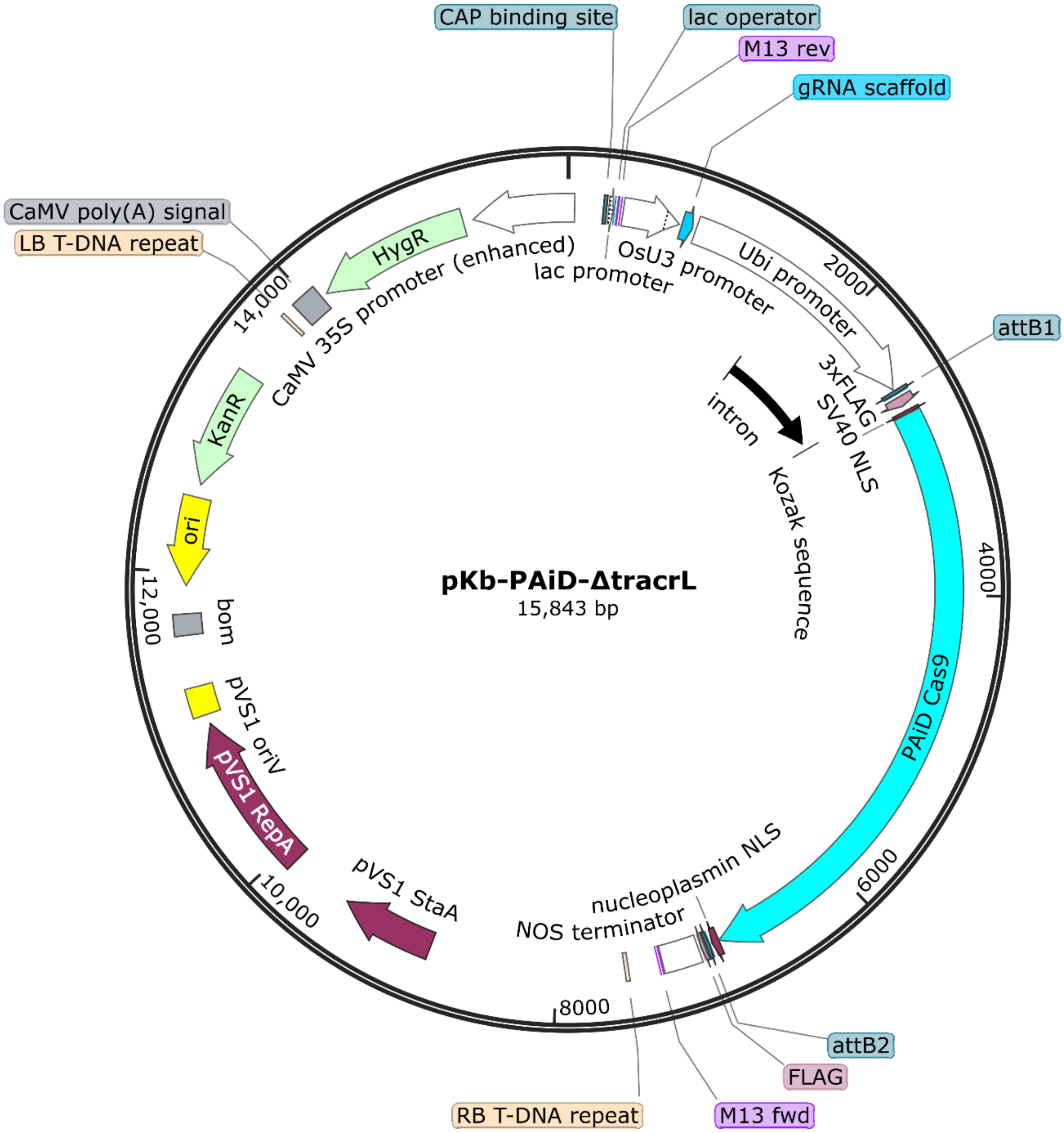


**Supplementary figure 5:** Schematic diagram showing different components of the pKb-PAiD-ΔtracrL mother vector, used for alternative tracrRNA experiments in rice. The sgRNA component is driven under OsU3 promoter (An RNA PolIII promoter, terminated by PolyT) and the endonuclease is driven under OsUbi10 promoter (An RNA PolII promoter, terminated by Nos terminator). The normal tracrRNA is replaced by a truncated long tracrRNA.

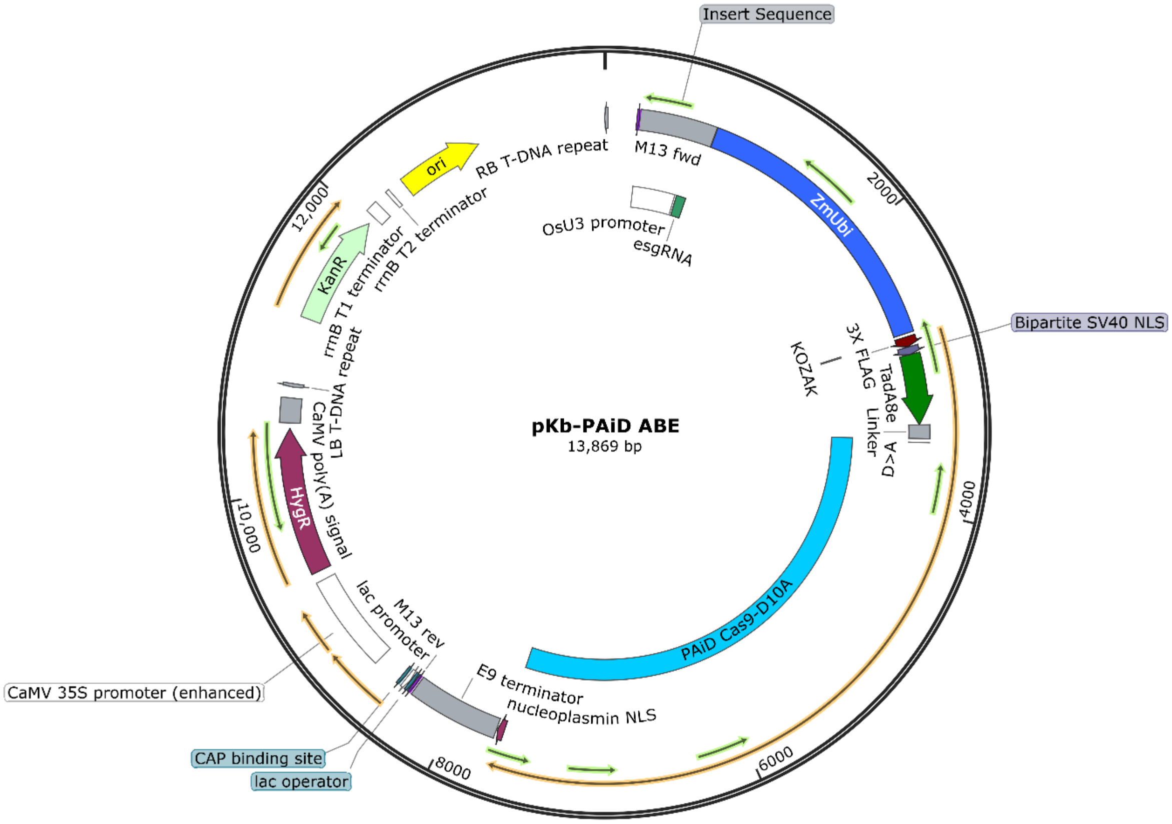


**Supplementary figure 6:** Schematic diagram showing different components of the pKb-PAiD-ABE mother vector, used for Adenine Base Editing experiments in rice. The sgRNA component is driven under OsU3 promoter (An RNA PolIII promoter, terminated by PolyT) while the endonuclease-nickase is driven by ZmUbi promoter (An RNA PolII promoter, terminated by E9 terminator). The Adenine deaminase has been fused to the N-terminal of the nickase PAiD using a short linker sequence.

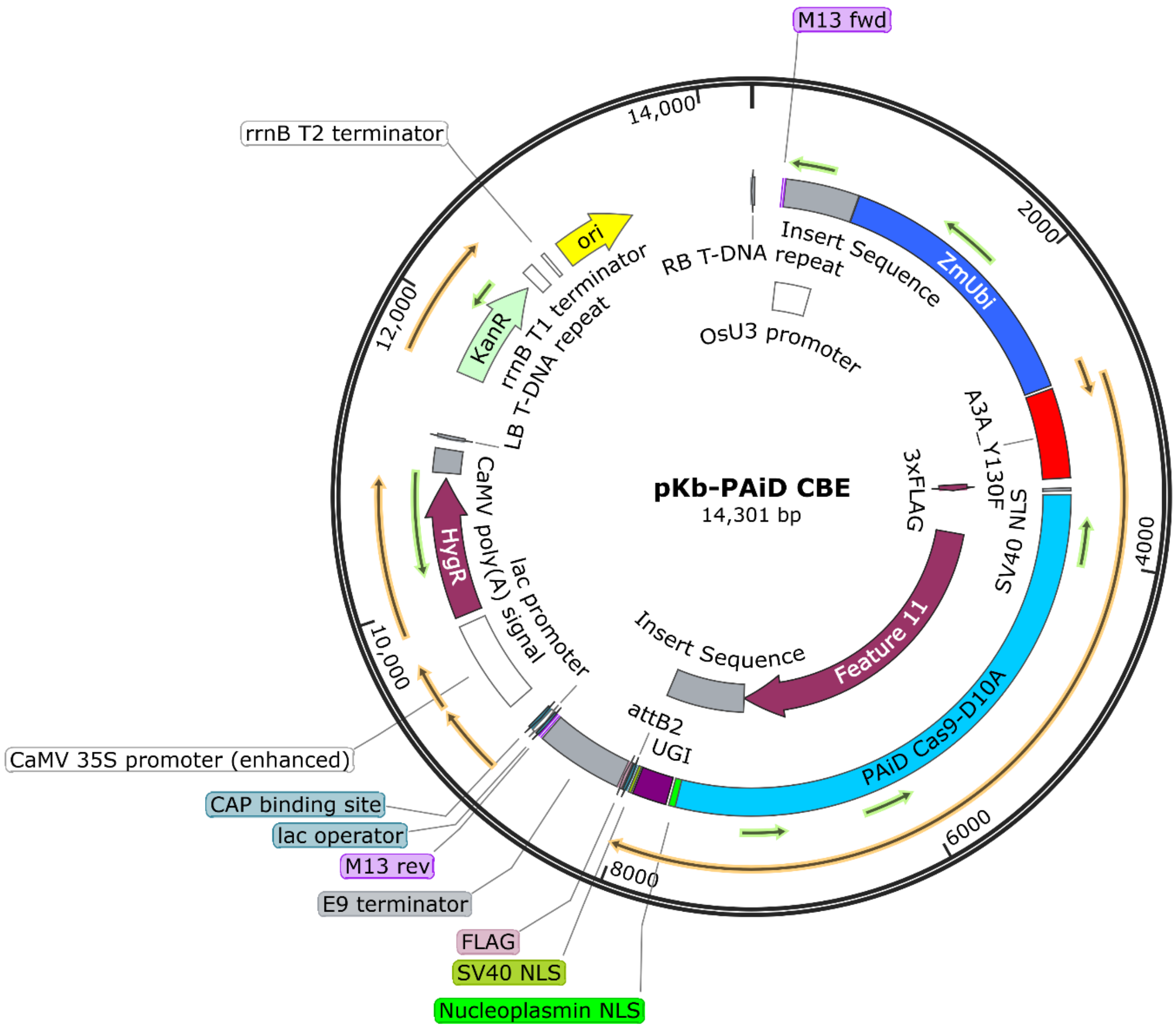


**Supplementary figure 7:** Schematic diagram showing different components of the pKb-PAiD-CBE mother vector, used for Cytosine Base Editing experiments in rice. The sgRNA component is driven under OsU3 promoter (An RNA PolIII promoter, terminated by PolyT) while the endonuclease-nickase is driven by ZmUbi promoter (An RNA PolII promoter, terminated by E9 terminator). The Cytosine deaminase has been fused to the N-terminal of the nickase PAiD, while the Uridine glycosylase inhibitor has been fused to the C-terminal of the nickase PAiD.

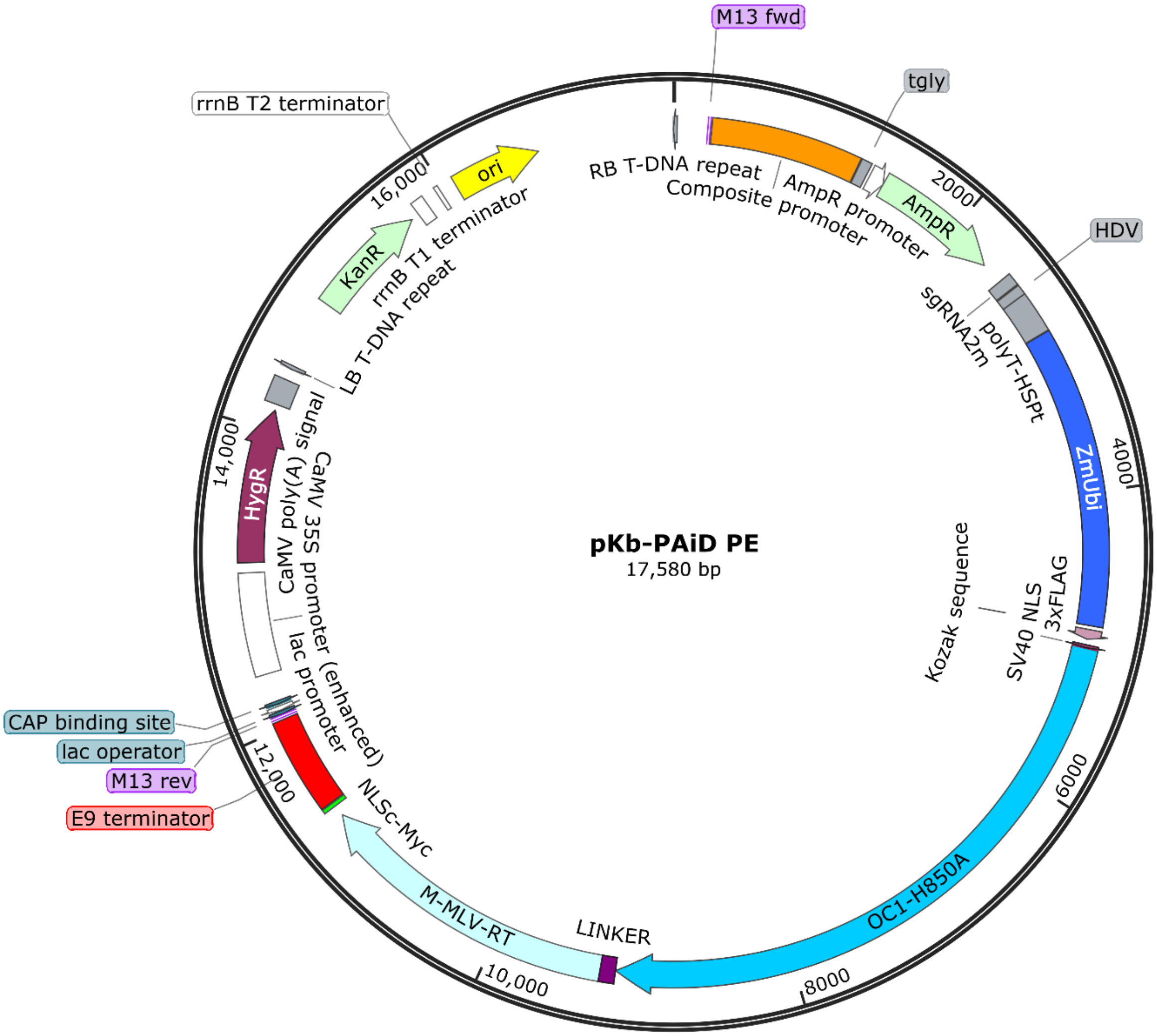


**Supplementary figure 8:** Schematic diagram showing different components of the pKb-PAiD PE mother vector, used for Prime Editing experiments in rice. The sgRNA2m component is driven under Composite Promoter while the endonuclease-nickase is driven by ZmUbi promoter (An RNA PolII promoter, terminated by E9 terminator). The M-MLV-RT has been fused to the C-terminal of the nickase PAiD.
